# Dual-acting Estrogen Receptor Modulator-Histone Lysine Demethylase Inhibitors

**DOI:** 10.64898/2026.05.26.728021

**Authors:** Dipak T. Walunj, Jeremiah O. Olugbami, Bocheng Wu, Ryan Kern, Alexis Johnston, Sonia Batan, Jabunnesa Khanom, Femi A. Egbeleke, Travis J. Nelson, Verjin Khodaverdian, Veronica Ventura, Nathaniel A. Hathaway, Muthusamy Thangaraju, Adegboyega K. Oyelere

## Abstract

Epigenetic dysfunction and the malfunction of endocrine proteins such as estrogen receptor alpha (ERα) are two key mechanisms for the sustenance of breast carcinoma. Agents that target ER signaling malfunctions are standard therapies for ERα-dependent breast cancer (BCa) subtypes. Small molecule inhibitors of epigenetic modifiers, histone lysine demethylases (KDMs), have shown promise as therapeutic agents for several BCa subtypes. We demonstrated herein that the integration of ERα antagonist (Tamoxifen) and agonist (17α-ethinylestradiol) ligands with deferiprone (DFP)-derived KDM inhibitor moiety furnished dual-acting agents Tam-KDMi and EED-KDMi, respectively. These agents showed robust on-target effects and potent BCa cells-selective anti-proliferative activities. The Tam-KDMi are cytotoxic to both ERα(+) and ERα(-) BCa cells, while the lack of ER-signaling inhibition conferred an enhanced ERα(-) cells cytotoxic to the EED-KDMi. Representative lead compounds Tam-KDMi **DW-116** and EED-KDMi **DW-088** and **DW-95** significantly reduce tumor growth in murine xenograft models of ERα(−) and ERα(+) BCas with TGI as high as 70%. Collectively, our data showed that **DW-116**, **DW-088** and **DW-95** have a high potential as leads for the development of new agents for the treatment of BCa subtypes regardless of the tumor ER expression status.

## 1. Introduction

Breast cancer (BCa) is second only to lung cancer as the leading cause of cancer deaths among US women.^1^ BCa subtypes are classified, based on gene expression patterns, as Luminal A, Luminal B, Basal, Claudin-low, and human epidermal growth factor receptor 2 (HER2).^2,3^ Most compounds developed clinically for BCa therapy exhibit greater efficacy for a subtype of BCa. In this regard, therapeutic interventions that capitalize on estrogen receptor (ER) signaling malfunctions, a driver of subtype that constitutes more than 70% of BCa cases,^4^ have enjoyed measured success in BCa therapy and/or chemoprevention. For example, estrogen-receptor modulators (ERMs) such as tamoxifen (Tam) and fulvestrant (Ful) are the first-line therapy for the treatment of hormone-dependent BCa. Other selective estrogen-receptor modulators (SERDs) are in development, and Elacestrant is the latest SERD to obtain clinical approval for the treatment of hormone-dependent BCa.^5^ However, despite initial benefits, most patients eventually relapse due to acquired resistance to these drugs.^6-8^ Additionally, there are no targeted treatment options for triple-negative breast cancer (TNBC), a BCa subtype lacking ER, HER2, and progesterone receptor (PR), which accounts for over 15-20% of BCa incidence.^3^

Another key mechanism that sustains BCa is epigenetic dysfunction.^9^ BCa viability, regardless of ER-expression status, has been shown to depend on an extensive network of epigenetic modifiers - histone mark writers, readers, and erasers. Bioinformatics and functional analysis have identified specific paralogs of amine oxidase and Jumonji family of histone lysine <u>d</u>e<u>m</u>ethylases (KDMs), <u>h</u>istone <u>m</u>ethyltransferases (HMTs), and <u>h</u>istone <u>d</u>e<u>ac</u>etylase (HDACs) as essential in supporting ERα signaling activation.^10^ Among these epigenetic modifiers, KDM1A, KDM3A, KDM5A, KDM5B, and KDM6A are exquisitely wired into ERα signaling and could be ideal targets for developing novel treatments for early and metastatic BCas. KDM1 is a flavin-dependent monoamine oxidase which demethylates histone H3 lysines 4 and 9 (H3K4 and H3K9) while members of the KDM5 and KDM6 families are 2-oxoglutarate- and Fe^2+^-dependent oxygenases that act as specific demethylases for H3K4 and H3K27me3, respectively.^11^ These KDMs regulate cell proliferation, differentiation, and stem cell self-renewal. Specifically, KDM1 is recruited to the ERα target gene promoters, where it modulates epigenetic changes via interactions with ER co-regulator protein PELP1.^12^ KDM5A and KDM5B are oncogenes that are overexpressed mostly in BCas and other tumors as well. KDM5A regulates many metastasis-related genes, which are required for BCa colonization in the lungs.^13^ KDM5B is overexpressed in ERα(+) BCas, functioning as a luminal lineage-driving oncogene and poor prognosis predictor.^14^ Similarly, KDM6A positively regulates gene expression programs associated with BCa proliferation and invasion.^15^

Members of the KDM subfamily are involved in other epigenetic reprograming events essential for the survival of BCas regardless of their receptor expression status. Ramadoss *et al* showed that KDM6B is required for TGF-β-induced EMT and BCa invasion via the H3K27me3 marks erasing activity of KDM6B.^16^ Subsequent analysis of the Oncomine database revealed that KDM6B level was markedly increased in invasive BCa compared with normal breast tissues, while a high expression of KDM6A is a predictor of poor prognosis in BCa patients.^17^ KDM5A and 5B are oncoproteins overexpressed in human BCa tumors and BCa cell lines (and other tumor cell lines), but not in normal adult breast tissue.^18,19^ Sharma *et al* have shown that increased expression of KDM5A promotes cancer stemness and enhanced resistance to anticancer agents.^20^ In addition, Lin *et al* have shown that KDM5A is required for BCa tumor formation in mouse models.^21^ Similarly, knockdown of KDM5B in BCa cells resulted in up-regulation of tumor suppressor genes including BRCA1, and decreased tumor formation potential *in vivo*.^22^ Dysfunction in the activities of other KDMs has been implicated in the development of several tumor types,^23,24^ suggesting the broad and significant roles of KDMs in the etiology of human cancers.

Based on the aforementioned literature evidence, KDMs have emerged as new anticancer drug targets with high therapeutic potential.^23^ While several KDM1A inhibitors are in clinical trials, the development of clinical candidates against the other KDMs is not as advanced.^25^ Genetic downregulation of KDM3A, KDM5A, KDM5B and KDM6A caused a significant decrease in the proliferation and invasiveness of BCa.^14,15,22,^ Pharmacological inhibition of each of these KDMs also caused BCa cell growth arrest *in vitro* and *in vivo*.^26^ There are, however, precedents for compensation among KDM paralogs in tumor models generated by selective deletion of a paralog member.^21^ Additionally, knockout mice of representative KDM paralogs presents a mild phenotype.^27,28.^ Compensation that may occur from KDM paralog-selective inhibition is not unique to cancer as a recent report noted that simultaneous inhibition of KDMs 6A and 7A is required to elicit a robust anti-inflammatory effect.^29^

We have previously discovered that clinically approved deferiprone (DFP) is a pan-selective inhibitor of KDMs implicated in BCa etiology; and a subsequent structure activity relationship (SAR) study furnished DFP-based KDM inhibitors that altered the velocity of HP1-mediated heterochromatin gene repression and displayed selective cytotoxicity against the BCa cell lines. Several of these compounds are preferentially more cytotoxic to the TNBC cell line and a representative compound significantly reduced tumor growth in xenograft mouse models.^30,53,58^ Encouraged by these findings, we investigated herein the integration of the DFP moiety into two structurally distinct ER modulator ligands as a strategy to further enhance the potency and/or selectivity of these DFP-based KDMi for both TNBC and ERα(+) BCas. We used Tam and 17α-ethinylestradiol (EED) as representative ER antagonist and ER agonist ligands, respectively, for this design. We designed the Tam-linked compounds (Tam-KDMi) by covalently linking the tertiary amino group of Tam to the aryl moiety of the aryl-triazolyl-DFP compounds identified from our prior study.^30,53,58^ Similarly, EED-linked DFP compounds (EED-KDMi) were envisioned to result from Huisgen cycloaddition reaction between the 17α-ethynyl moiety of EED and appropriate azidoalkyl-DFP (Fig. 1). This design was predicated on the analysis of the co-crystals of the active metabolite of Tam (PDB: 3ERT)^31^ and an analog of EED, TFMPV-E2 (PDB:2P15),^32^ bound to the ligand binding domain of ERα. These structures revealed that the tertiary amino group of Tam and the C-17 moiety of EED are solvent-exposed and may be suitable as conjugation locales for tethering Tam and EED to other pharmacophores with minimal disruption to ERα binding. We have shown that analogously modified Tam and EED retained ERα binding.^33^

**Figure 1:**
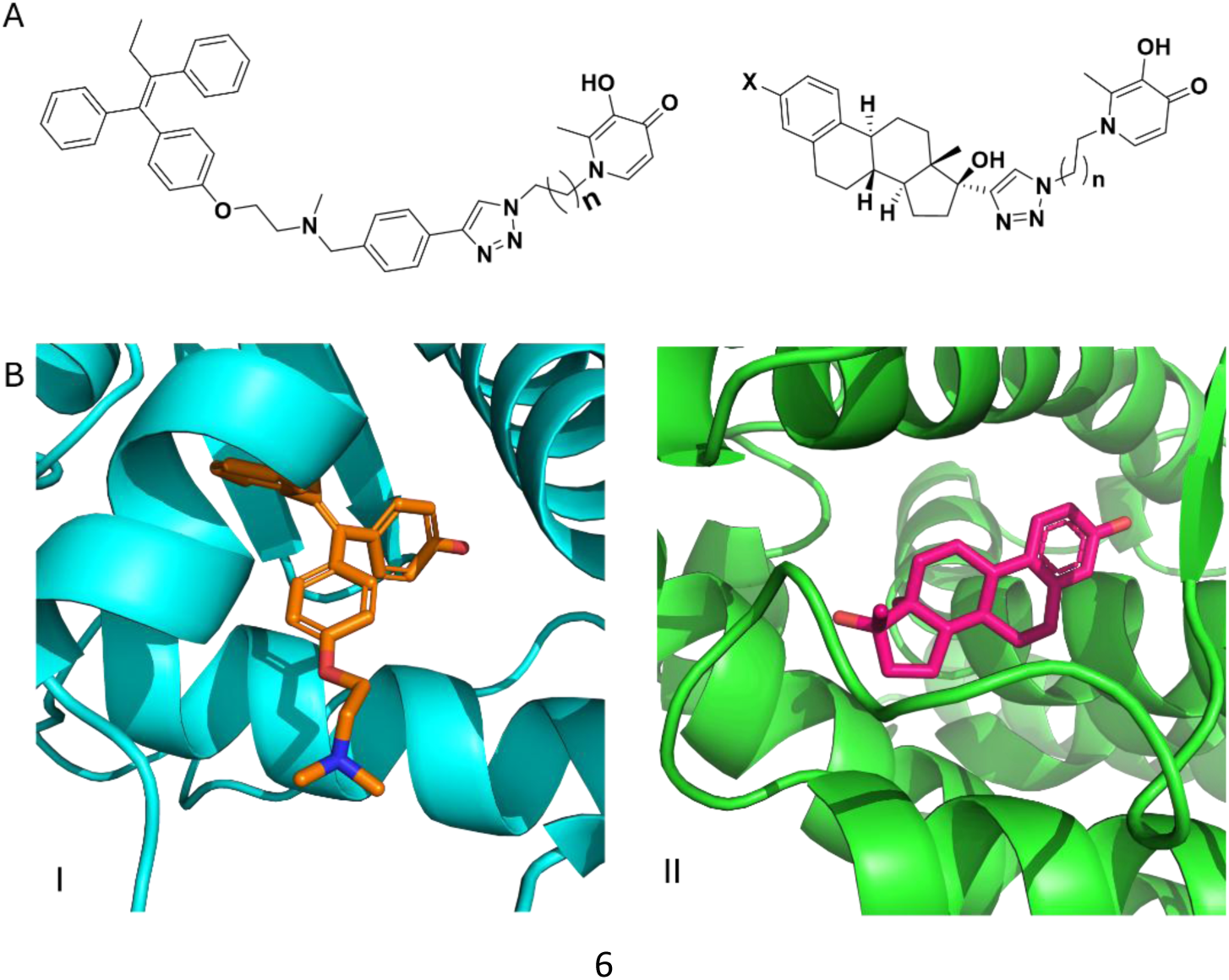
(A) Generalized structures of the designed Tam-KDMi and EED-KDMi. (B) Co-crystals of ERα and the active metabolite of Tam (PDB: 3ERT)^31^ (I) and EED analog, TFMPV-E2 (PDB:2P15)^32^ (II) revealed that the tertiary amino group of Tam and the C-17 moiety of EED are solvent-exposed, making them ideal modification points.

The designed Tam-KDMi and EED-KDMi were synthesized and evaluated for KDM inhibition, ERα binding and antiproliferative/antitumor activities in *in vitro* and *in vivo* models. Our results revealed that these dual-acting molecules possessed independent KDM inhibition and ERα binding activities. Evaluation of the intracellular markers revealed that the antiproliferative activities of these compounds are derived from strong on-target effects. The EED-KDMi are selectively cytotoxic to MDA-MB-231, a representative TNBC cell line, while MCF-7, a representative ERα(+) BCa cell line, is less responsive. Interestingly, Tam-KDMi are equally cytotoxic to both MDA-MB-231 and MCF-7 cell lines. We also found that representative Tam-KDMi **DW-116** and EED-KDMi **DW-088** and **DW-95** significantly reduce tumor growth in murine xenograft models of ERα(−) and ERα(+) BCas. Collectively, our data demonstrates that simultaneous inhibition of ER signaling and KDM by Tam-KDMi resulted in cytotoxicity to both ERα(+) and ERα(-) BCa cells while the lack of ER-signaling inhibition makes the EED-KDMi more cytotoxic to MDA-MB-231 cells. Given their in *vivo* efficacy, compounds **DW-116 DW-088** and **DW-95** have a high potential as leads for the development of new agents for the treatment of BCa subtypes regardless of the tumor ER expression status.

## 2. Results and Discussion

### 2.1 Design of Estrogen Modulator–KDMi Conjugates

Our interest in developing ER modulator-KDM dual inhibitors is predicated on literature precedence implicating KDM dysfunctions in the viability of BCas^4-8^ and our discovery of a cohort of DFP-based KDMi which inhibits a collection of KDM paralogs essential for BCas viability.^30,53,58^ To design the proposed compounds, we incorporated the DFP moiety, through methylene linkers of variable length, into the tertiary amino and 17α-ethynyl which are solvent exposed moieties of Tam and EED, respectively, when bound to ERα (Fig. 1). Analysis of their docked poses at the active site of KDM6a (PDB:3AVR) revealed that DFP and representative compounds **DW-088** and **DW-122** adopt poses in which they strongly chelate to the active site Fe^2+^ ion. In addition, DFP and the DFP moiety of **DW-088** and **DW-122** are within distances that could enable further stabilizing H-bonding interactions between their oxygen moieties and two key residues within the KDM6A active site – phenolic group of Tyr 1135 and OH group of Ser 1154 (Fig. 2a-c). The N-1 group of DFP is oriented out of the active site (Fig. 2a). This orientation allows further modifications to enhance KDM binding affinity and inhibitory activity as we previously demonstrated.^30^ Furthermore, we observed that the ER-binding moieties of **DW-088** and **DW-122** are accommodated within the outer rim leading to the active site of KDM6A where they are engaging in potentially stabilizing interactions that DFP lacks (Fig. 2d-f). It seems that, relative to the tamoxifen moiety of **DW-122**, the estradiol moiety of **DW-088** is better accommodated at the outer rim. This deduction is based on the improvement in the docking scores of the ER modulator-linked compounds being more negative compared to DFP (-4.972 kcal/mol). However, the dock score of the estradiol-derived compound **DW-088** (-8.979 kcal/mol) is more negative than that of the tamoxifen-derived compound **DW-122** (-6.881 kcal/mol). This observation suggests the importance of the KDM6A outer rim interactions in the overall binding affinities of **DW-088** and **DW-122**.

**Figure 2:**
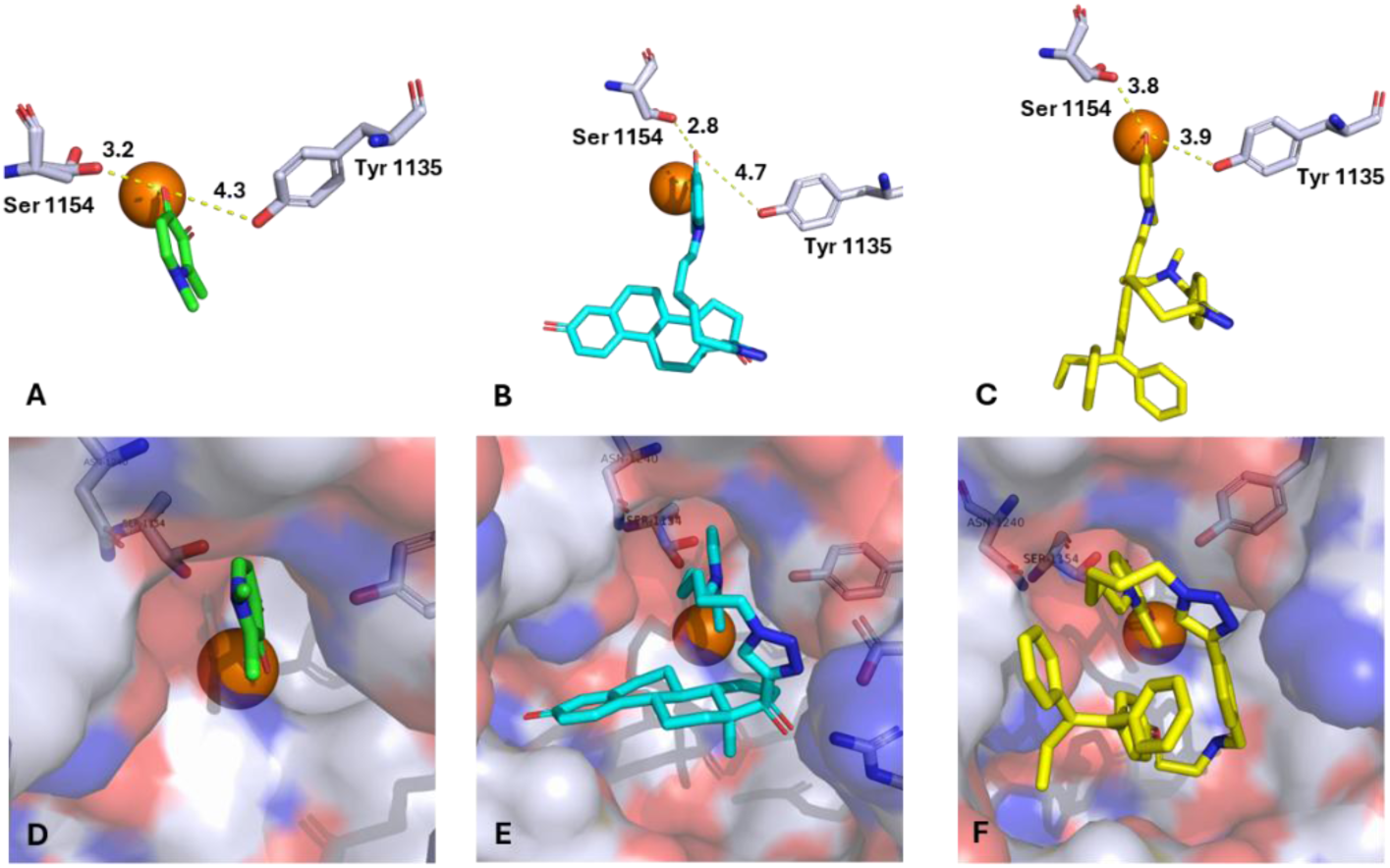
Molecular docking of DFP (green), **DW-088** (blue) and **DW-122** (yellow) **(A-C)** using KDM6a (PDB:3AVR) revealed that the chelation of the Fe^2+^ ion and H-bonding interactions with Ser 1154 and Tyr 1135 are stabilizing forces that are conserved regardless of the modification at the DFP N-1 position. **(D-F)** Surface presentation of the docked pose of DFP relative to those of **DW-088** and **DW-122** revealed that the ERα binding moieties of **DW-088** and **DW-122** are accommodated at the outer rim leading to KDM6a active site, allowing Fe chelation.

To evaluate the ERα interaction of the designed compounds, we docked the estradiol-derived compound **DW-088** against an ER agonist crystal structure (PDB: 2P15), while the tamoxifen-derived compound **DW-122** was docked against an ER antagonist crystal structure (PDB: 3ERT). Docking against 2P15 shows a nearly complete overlap between EED and the estradiol moiety of **DW-088** and adopting an orientation in which their 3-OH moiety is engaging in hydrogen bonding with the ARG-394 and GLU-353 residues found in the base of the binding pocket (Fig. 3a-b). In contrast, **DW-95**, which has a methoxy group at its C-3 position, cannot engage in H-bonding with ARG-394 and GLU-353 due to the methylation. Based on dock scores, EED (-10.487 kcal/mol) has the strongest affinity for 2P15, followed by **DW-088** (-8.956 kcal/mol) and **DW-95** (-7.932 kcal/mol). The difference in dock scores of **DW-088** and **DW-95**, can be explained by the loss of H-bonding to with ARG-394 and GLU-353 due to methylation. The drop in dock score between the **DW-088** and EED could be due to the possibility of energy penalty that results from the navigation of the DFP and linker moieties of **DW-088** through the narrow tunnel leading to the active site of ERα agonist form.

**Figure 3:**
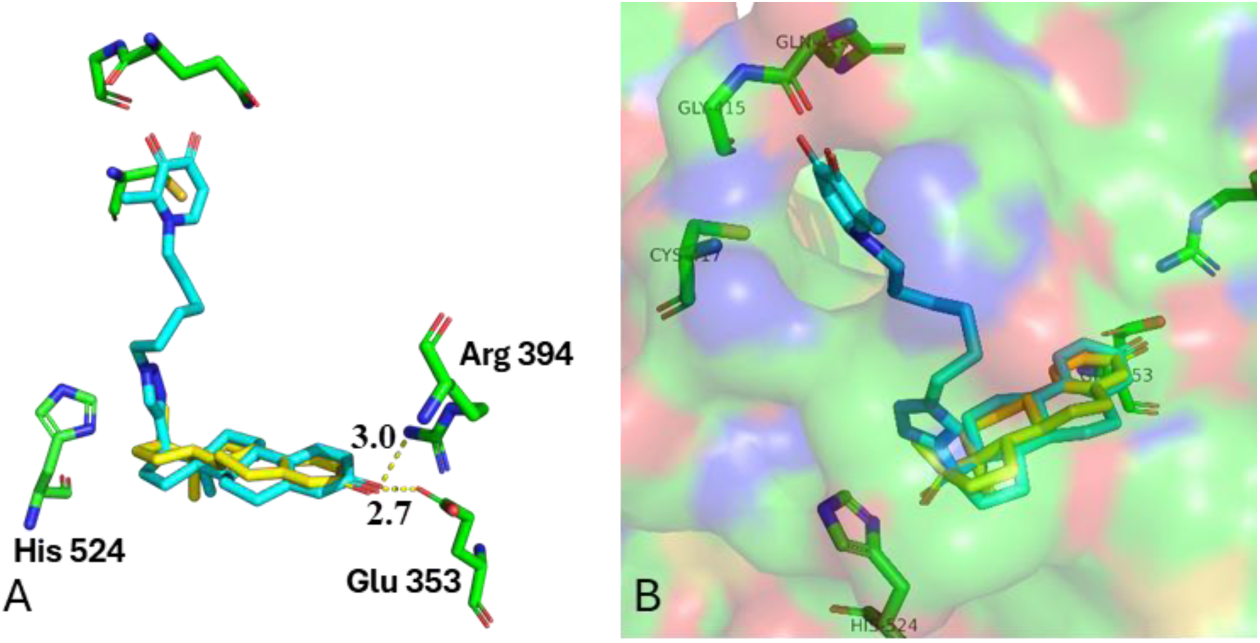
(A) Overlay of the docked poses of **DW-088** (blue) and EED (yellow) revealed near identical orientation of the steroid rings and hydrogen bonding with Arg-394 and Glu-353 at the base of the active site of ERα agonist form (PDB: 2P15). (B) The DFP and linker moieties of **DW-088** navigate through the narrow tunnel leading out of the active site to be oriented at the outer rim of ERα agonist binding mode.

Overlay of the docked outputs of tamoxifen and **DW-122** against 3ERT revealed a near identical overlap of tamoxifen and the tamoxifen moiety of **DW-122** (Fig. 4a). Due to the relative openness of the ER antagonist crystal structure, the alkyl linker and DFP moieties of **DW-122** have a high degree of flexibility, adopting a folded conformation that allows its DFP group to engage in potential H-bonding interactions with the backbones of Cys 530 and Val 534 (Fig. 4b). These extra interactions could explain the enhanced binding of **DW-122** (-10.183 kcal/mol) relative to that of tamoxifen (-9.02 kcal/mol).

**Figure 4:**
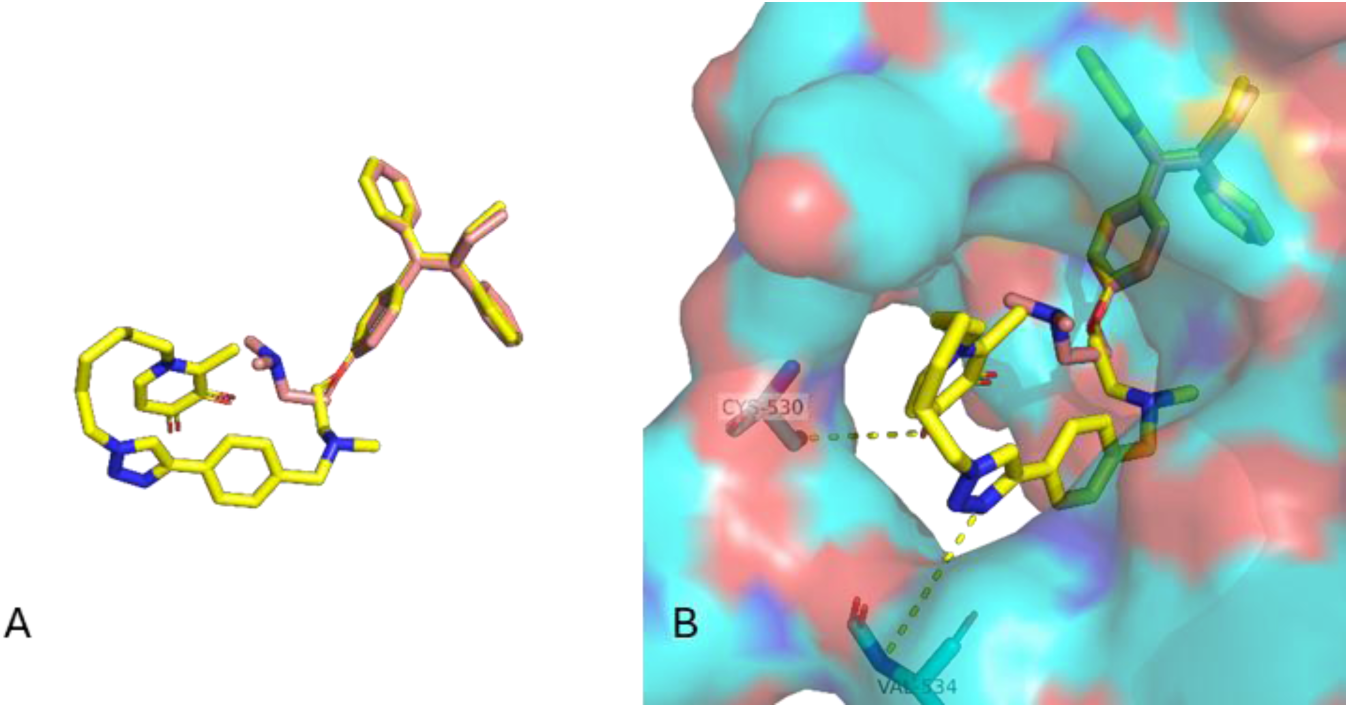
(A) Overlay of the docked poses adopted by **DW-122** (yellow) and tamoxifen (salmon) at the active site of the antagonist form of ERα (PDB: 3ERT) revealed strong overlap of tamoxifen and the tamoxifen moiety of **DW-122**. (B) Surface presentation of the docked pose of tamoxifen relative to that of **DW-122** revealed that the DFP and triazole moieties of **DW-122** are engaging in potentially stabilizing interactions with the backbones of Cys 530 and Val 534 within the tunnel leading to the active site of ERα in the agonist binding mode (PDB: 3ERT).

Encouraged by this *in silico* analysis, we synthesized all the designed compounds shown in Fig. 1 and investigated their effects on KDM and ERα signaling and anticancer activities using i*n vitro* and *in vivo* models.

### 2.2 Synthesis of Estrogen Modulator–KDMi Conjugates

The synthesis of the designed ER modulator–KDMi is accomplished as illustrated in Schemes 1, 2, and 3. First, the synthesis of the linker began with the preparation of PMB-protected maltol **1** through alkylation of maltol with 4-methoxybenzyl bromide in acetone, using potassium carbonate as a base.^30,53,58^ Subsequently, **1** was reacted with azido-amine intermediates **2** (**a-g**), with NaOH as base, to afford the maltol-azido intermediate compounds **3 (a-g)** in moderate yield using our established protocol, as shown in Scheme 1.^30,53,58.^ The synthesis of Tam-KDMi began with the demethylation of tamoxifen using chloroethyl chloroformate and methanol. The resulting desmethyl tamoxifen **4** was alkylated with 1-(bromomethyl)-4-ethynylbenzene using Hünig’s base to provide compound **5** according to our previously established protocol.^33^ Subsequently, copper(I)-catalyzed cycloaddition reaction of the **5** with maltol-azido compounds **3 (a-f)** resulted in the intermediate compounds **6 (a-f).** Finally, these intermediates were deprotected using TFA to obtain the desired Tam-KDMi in moderate to good yields (Scheme 2).

**Scheme 1:**
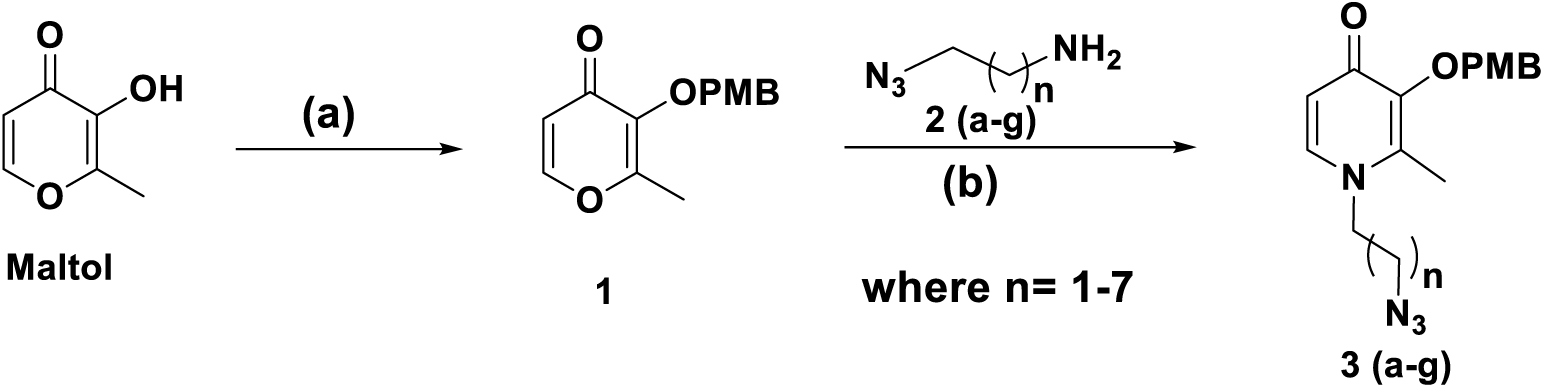
Synthesis of Linker. Reagent and conditions: (a) 4-methoxy benzyl bromide, K_2_CO_3_, Acetone, 56 ^°^C, 12 h, 86%; (b) NaOH, Water: Methanol (0.4:0.6),105 ^°^C, 24 h, 40-56 %.

**Scheme 2:**
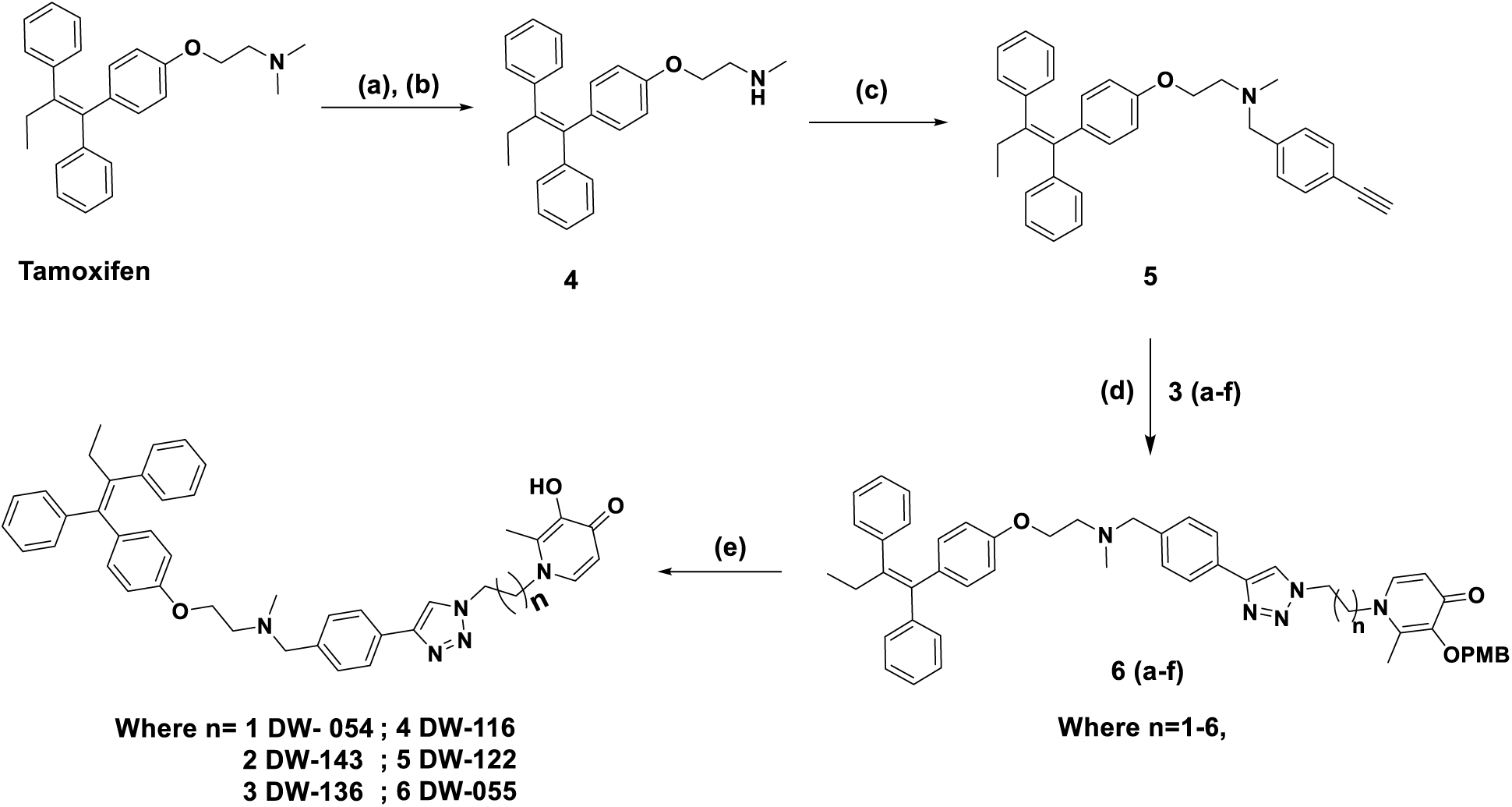
Synthesis of Tamoxifen-derived KDMi. Reagent and conditions: (a) 1 Chloroethylchloroformate,1,2 dichloroethane, 90 °C; (b) MeOH, reflux, 3 h, 94 % after two steps; (c) 1-(bromomethyl)-4-ethynylbenzene, DIPEA, CH_2_Cl_2_, rt, 24 h, 82 %; (d) CuI, DIPEA, THF: DMSO (1:1), rt, 15 h, (e) 10 % TFA in CH_2_Cl_2_, 0 ^°^C to rt, 1h, 50-70 % after two steps.

**Scheme 3:**
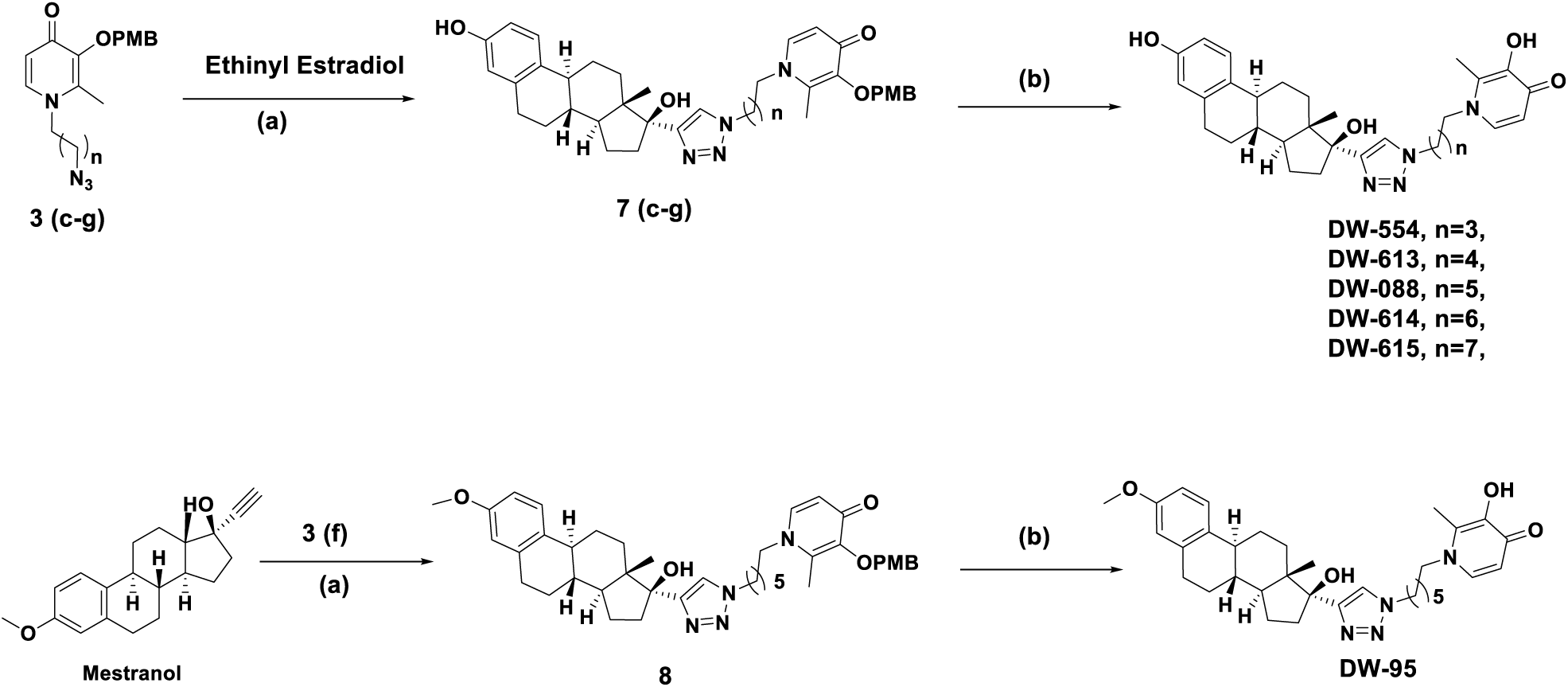
Synthesis of Steroid-derived KDMi. Reagent and conditions: (a) CuI, DIPEA, THF: DMSO (1:1), rt, 15 h, b) 10 % TFA in CH_2_Cl_2_, 0 °C to rt, 1 h, 48-71% after two steps.

The synthesis of the EED-KDMi began with copper(I)-catalyzed cycloaddition reaction of maltol-azido compound **3 (c-g)** and EED, resulting in the penultimate intermediates **7 (c-g)** in excellent yields. The PMB group was removed by treatment with 5% TFA in CH_2_Cl_2_, yielding the desired series of EED-KDMi in moderate to good yields (Scheme 3). Similarly, copper(I)-catalyzed cycloaddition reaction of maltol-azido **3f** with Mestranol afforded the desired intermediate compounds **8**, and the PMB group was removed by treatment with 5% TFA in CH_2_Cl_2_, yielding the desired compound **DW-95** in 71 % after two steps.

### 2.3 Target Validation – KDM inhibition

We used the chromatin *in vivo* assay (CiA) in mouse embryonic stem (mES) cells to investigate the effects of the Tam-KDMi and EED-KDMi on chromatin dynamics, which is a direct indicator of demethylase activities of KDMs within the cell. CiA measures heterochromatin formation speed by leveraging chemical induced proximity (CIP) to recruit the chromoshadow of heterochromatin protein 1 (HP1) to an *Oct4* locus modified to express an enhanced green fluorescent protein gene.^30^ We observed that Tam-KDMi and EED-KDMi caused inhibition of HP1-induced heterochromatin formation within 48 h of exposure to CiA mES cells with IC_50_s that are linker-length dependent (Table 1). Within the Tam-KDMi series, the five-methylene linked (C-5) compound **DW-116** most potently inhibited KDM activities with an IC_50_ of 161 nM while the C-4 linked compound **DW-136** is the least potent with an IC_50_ of 853 nM. Interestingly, the shorter linker compounds **DW-054** (IC_50_ = 417 nM) and **DW-143** (IC_50_ = 675 nM) and the longer linker compounds **DW-122** (398 nM) and **DW-055** (351 nM) were more potent than **DW-136**. Although they also exhibited linker length-dependent KDM inhibition activities, the EED-KDMi compounds have slightly weaker potency compared to the Tam-KDMi. Compound **DW-554**, the EED-KDMi with the shortest linker length (C-4) exhibited no KDM inhibition up to the maximum tested concentration of 10 μM. Potency increased with linker length up to C-6 (C-5 compound **DW-613**, IC_50_ =1.36 μM and C-6 compound **DW-088**, IC_50_ = 570 nM) and then dropped with further increase in linker length (C-7 compound **DW-614**, IC_50_ = 680 nM and C-8 compound **DW-615**, IC_50_ = 2.72 μM). Interestingly, the methylation of the EED 3-OH group attenuated KDM inhibition as the methyl-EED compound **DW-095** (IC_50_ = 966 nM) is slightly less potent than **DW-088**, the corresponding EED analog with the same linker length (Table 1). Collectively, this data strongly demonstrates that the Tam-KDMi and EED-KDMi possess intracellular KDM inhibitory activity and it is supportive of our *in silico* predictions; thus, validating our design.

**Table 1:** KDM inhibition activity of Tam-KDMi and EED-KDMi.

| Compound | IC<sub>50</sub> (μM) ± SD |
| --- | --- |
| **DW-054** | 0.417 ± 0.09 |
| **DW-055** | 0.351 ± 0.16 |
| **DW-116** | 0.161 ± 0.07 |
| **DW-122** | 0.398 ± 0.10 |
| **DW-136** | 0.853 ± 0.04 |
| **DW-143** | 0.676 ± 0.04 |
| **DW-554** | ND |
| **DW-613** | 1.358 ± 0.14 |
| **DW-088** | 0.570 ± 0.14 |
| **DW-614** | 0.680 ± 0.12 |
| **DW-615** | 2.720 ± 0.10 |
| **DW-095** | 0.966 ± 0.04 |

ND = less than 50% inhibition at 10 μM, SD = standard deviation

### 2.4 Target Validation – ERα-binding affinity

Next, we investigated the binding interactions of the Tam-KDMi and EED-KDMi with ERα based on cell-free assays. As shown in Table 2 and Suppl Info Figure S1, all the Tam-KDMi binds to ERα with nanomolar range binding affinities (200 nM to 590 nM) and stronger than that of tamoxifen (810 nM). Among the cohort of Tam-based compounds, it seems chain length does not play a significant role on the compounds’ ERα binding affinity. Specifically, the C-2, C-4 and C-5 compounds, **DW-054**, **DW-136** and **DW-116** respectively, have nearly identical binding affinity while the other compounds – **DW-143** (C-3), **DW-122** (C-6) and **DW-055** (C-7) – are only about 2- to 2.6-fold weaker. The stronger binding of the Tam-KDMi relative to tamoxifen agrees with our *in silico* docking predictions that the DFP group of the Tam-KDMi could participate in potentially stabilizing H-bonding interactions with the backbones of Cys 530 and Val 534 (Fig. 4b). While most of the EED-KDMi bind to ERα with similar binding affinities, which are comparable to that of tamoxifen, they have much weaker binding relative to EED (binding affinity = 30 nM). Among longer linker EED-KDMi, there seems to be a preference for the even chain compounds (comparing even chain compounds **DW-088** and **DW-615** with the odd chain **DW-613**). The methyl-EED compound **DW-095** has the weakest ERα binding affinity among the EED-KDMi. Despite having the same methylene linkers as **DW-088**, the presence of a methyl group in **DW-095**, instead of OH group in the other EED-KDMi, may account for the decline in ERα binding affinity of **DW-095**, an outcome that was predicted by our molecular docking analysis. This data revealed that Tam-KDMi and EED-KDMi retained the ERα binding attribute of their parent tamoxifen and EED. However, the consequence of integration of the DFP moiety on ERα binding is opposite, enhancing ERα binding for Tam-KDMi and significantly reducing ERα binding for EED-KDMi.

**Table 2:** ERα-binding affinity of Tam-KDMi and EED-KDMi.

| Compound | IC<sub>50</sub> (μM) |
| --- | --- |
| **DW-054** | 0.22 |
| **DW-143** | 0.59 |
| **DW-136** | 0.20 |
| **DW-116** | 0.24 |
| **DW-122** | 0.43 |
| **DW-055** | 0.43 |
| **DW-554** | 1.48 |
| **DW-613** | 1.38 |
| **DW-088** | 1.05 |
| **DW-614** | 1.91 |
| **DW-615** | 1.05 |
| **DW-095** | 5.77 |
| **Tam** | 0.81 |
| **EED** | 0.03 |

### 2.5 Tam-KDMi and EED-KDMi demonstrate strong cytotoxic effects against cancer cell lines

We proceeded to investigate the antiproliferative effects of Tam-KDMi and EED-KDMi using the MTT assay. Initially, we focused on BCa cell lines MCF-7 (ERα(+)), MDA-MB-231(ERα (-)), lung cancer cell line A549, and VERO, a normal kidney epithelial cell line. For comparison of bioactivity, we incorporated the parent compounds (EED and Tam) as reference compounds. The cells were treated with 1% DMSO solution of each compound for 72 h, and cell viability was determined using the MTT assay. As shown in Table 3, the parent compounds - DFP, EED, and tamoxifen - inhibited the proliferation of the tested cells with IC_50_ values that are in close agreement with the literature data.^30,33-37^ Interestingly, Tam-KDMi exhibit significantly higher antiproliferative activity against all the cell lines relative to the parent DFP and tamoxifen, with IC_50_ in the mid-nanomolar to low micromolar range and potency enhancement as high as 35-fold relative to tamoxifen. Comparing the tested cell lines, Tam-KDMi show mild preferential toxicity against MCF-7 and MDA-MB-231 as evidenced by their slightly higher *in vitro* therapeutic indexes (IVTI) relative to tamoxifen (Table 3). Tamoxifen has been reported to elicit toxicity to the TNBC cancer cell line MDA-MB-231 via ERα-independent mechanisms.^39,40^ It is likely that the enhanced potency of Tam-KDMi is the result of synergistic inhibition of KDM and ERα and perturbation of these tamoxifen ERα-independent mechanisms. Although the EED-KDMi are more potent than the EED and DFP building blocks, they are less cytotoxic than Tam-KDMi. Similar to Tam-KDMi, however, EED-KDMi are much more cytotoxic against MCF-7 and MDA-MB-231 than A549 and Vero cells. Also, their antiproliferative effects showed linker length-dependence that peaked with the C-6 compound **DW-088**. The somewhat lower antiproliferative effects of the EED-KDMi cohort, relative to Tam-KDMi, are compensated for by their much higher cancer cell selective toxicity. In this regard, **DW-088** exhibited the highest cancer cell selective toxicity with IVTI as high as 59.2 (Table 3). Paradoxically, the methyl-EED compound **DW-095** is a clear exception among the EED-KDMi, being the most cytotoxic and least selective. Although **DW-095** inhibits KDM activity in CiA with a high nanomolar IC_50_ (Table 1), it binds to ERα with the weakest affinity among the compounds disclosed herein (Table 2). The enhanced antiproliferative effects could likely be due to mechanisms other than KDM inhibition and ERα binding.

**Table 3.**
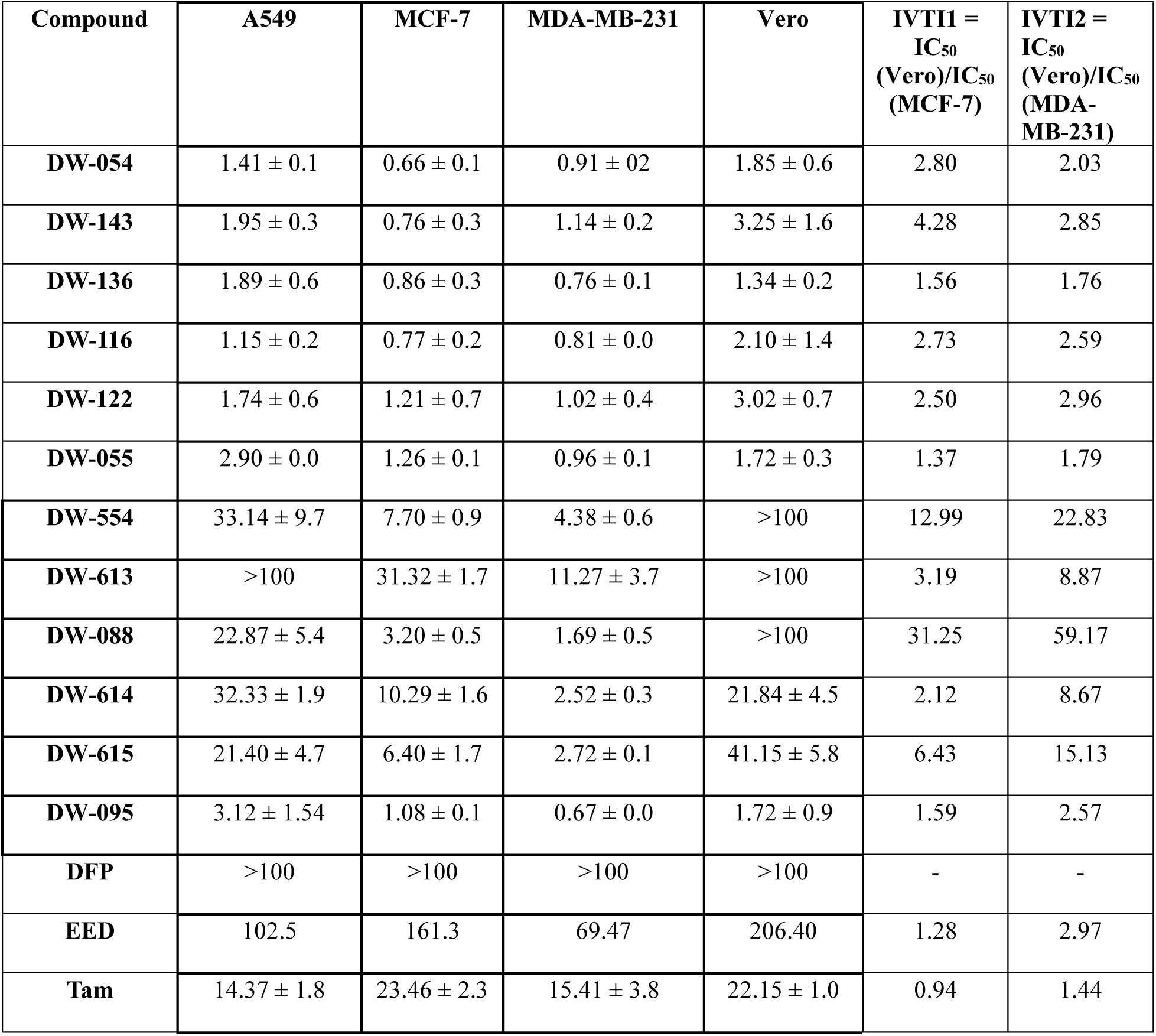
Antiproliferative activity of Tam-KDMi and EED-KDMi.

In light of these observations, we extended our study to four additional cancer cell lines – two prostate (DU-145 and LNCaP), two liver (Hep-G2 and SK-HEP-1) to investigate the therapeutic potential of these compounds beyond TNBC and ERα(+) BCas. We chose these cell lines because EED and tamoxifen have demonstrated conflicting, yet promising therapeutic effects against prostate and liver cancers.^41-44^ Gratifyingly, we observed that Tam-KDMi, EED-KDMi, and the template compounds inhibited the proliferation of these four cell lines with an inhibition pattern that closely resembled their effects against the BCas cell lines MCF-7 and MDA-MB-231 (Suppl info Table S1 and Figs. S2-S3). This data suggests that the KDM inhibition activity and/or perturbation of the ER-independent pathways targeted by the template compounds tamoxifen and EED significantly contributed to the antiproliferative effects of these compounds. More importantly, the data demonstrate that the disclosed Tam-KDMi and EED-KDMi possess selective toxicity against diverse cancer cell lines. Furthermore, based on IVTI, both sets of Tam-KDMi and EED-KDMi exhibit higher IVTI values than tamoxifen in both MCF-7 and MDA-MB-231 cells.

### 2.6 Tam-KDMi and EED-KDMi compromise the clonogenic ability of MDA-MB-231 cells

To determine the influence of representative Tam-KDMi and EED-KDMi on the ability of a single cell to become a colony of cells, we carried out a clonogenic/colony-formation assay using MDA-MB-231 as a representative cancer cell line. This assay assesses the reproductive capacity of the treated cells.^45^ For this experiment, we tested with Tam-KDMi **DW-122** and EED-KDMi **DW-088** at three different concentrations ranging from 0.5× IC_50_ to 2× IC_50_. DFP and Tam were used as control compounds. We determined the colony formation efficiency using both colony number counting and absorbance-based quantification.^46^ In agreement with its enhanced cytotoxicity, we observed that Tam is more effective than DFP in attenuating clonogenicity of MDA-MB-231 cells. Although both Tam-KDMi **DW-122** and EED-KDMi **DW-088** significantly inhibited the clonogenicity of MDA-MB-231 cells, **DW-122** is more effective at suppressing clonogenicity at lower concentrations (Fig. 5a-c). The seeming disparity observed between colony number counting and absorbance-based quantifications for **DW-088** (Fig. 5b vs Fig. 5c) at the lowest concentration may be associated with the fact that a colony is considered as a group of at least 50 cells. The implication here is that, although there might be colonies found, cells not making up a colony still existed. This scenario is not applicable to **DW-122** as colony quantification data are in excellent agreement with the absorbance-based quantification. Overall, both representatives of Tam-KDMi **DW-122** and EED-KDMi **DW-088** elicited anticlonogenic activity with **DW-122** being more effective.

**Figure 5:**
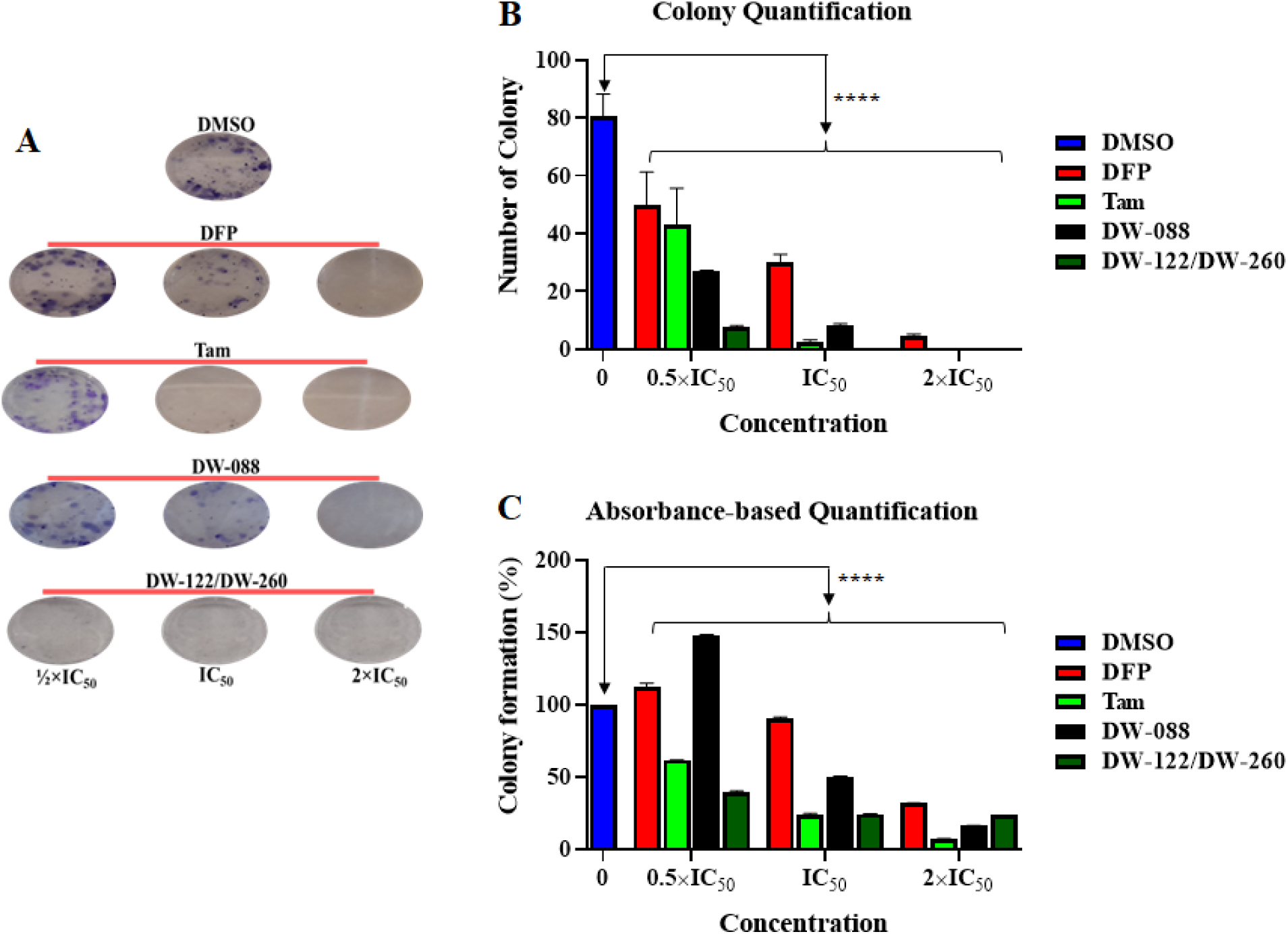
Assessment of the effects of Tam-KDMi **DW-122** and EED-KDMi **DW-088** on the colony formation ability of MDA-MB-231. Cells were treated for 72 h with the respective concentrations of ½× IC_50_, 1× IC_50_, and 2× IC_50_ values of each compound. The treatment media were removed after 72 h and the cells were subsequently maintained on fresh media for 10 days. \*\*\*\**p* < 0.0001.

### 2.7 Tam-KDMi and EED-KDMi modulate cell cycle progression in MCF-7 and MDA-MB-231 cells

To further probe the mechanism of action of the disclosed compounds, we investigated the effects of representative Tam-KDMi (**DW-116** and **DW-122**), EED-KDMi (**DW-088** and **DW-095**), and tamoxifen on cell cycle progression. We used MCF-7 and MDA-MB-231 cells for this experiment. We observed that none of the tested compounds significantly altered cell cycle progression in both cell lines after 24 h treatment at their respective IC_50_ relative to the negative (DMSO) control (Suppl info Figs. S4-S5). However, on increasing the concentrations of treatment to 2×IC_50_, we observed significant changes in different phases of the cell cycle. While Tam did not significantly alter MCF-7 cell population at G1 phase, its effect was more pronounced at G2/M phase, as reported previously.^47^ We also found that **DW-116**, **DW-122,** and **DW-088** significantly arrested cells in S-phase of the cell cycle with **DW-122** further significantly arresting cells in G2/M phase (Fig. 6). A direct implication of the S-phase arrest is that these compounds act through prevention of DNA synthesis and inhibition of cell proliferation with a likelihood of induction of cell death.^44^ Furthermore, **DW-095**, an EED-KDMi, displayed similar trends as tamoxifen on cell cycle progression. In the case of MDA-MB-231 cells, we also observed that the new compounds elicited an increase in S phase cell population reaching a significant level, specifically for **DW-122** (Fig. 7). We reported previously that DFP, presumably through its KDM inhibition activity, can significantly induce an S-phase arrest.^53^ Therefore, it is likely that the cell cycle arrest induced by the new compounds may be derived principally from their KDM inhibition activities. This inference is supported by other literature observations indicating that inhibition/knockdown of certain KDM paralog members could result in S-phase arrest of the cell cycle.^49^ Additionally, **DW-095** significantly induced an increase in cell population in G2/M phase, which may account for its higher cytotoxic effects in MDA-MB-231 cells when compared with MCF-7 cells. Importantly, these drugs also resulted in induction of apoptosis (i.e. sub-G1 cells) in both cell lines, evidenced by the less than 100% total cell population in the treated groups.

**Figure 6:**
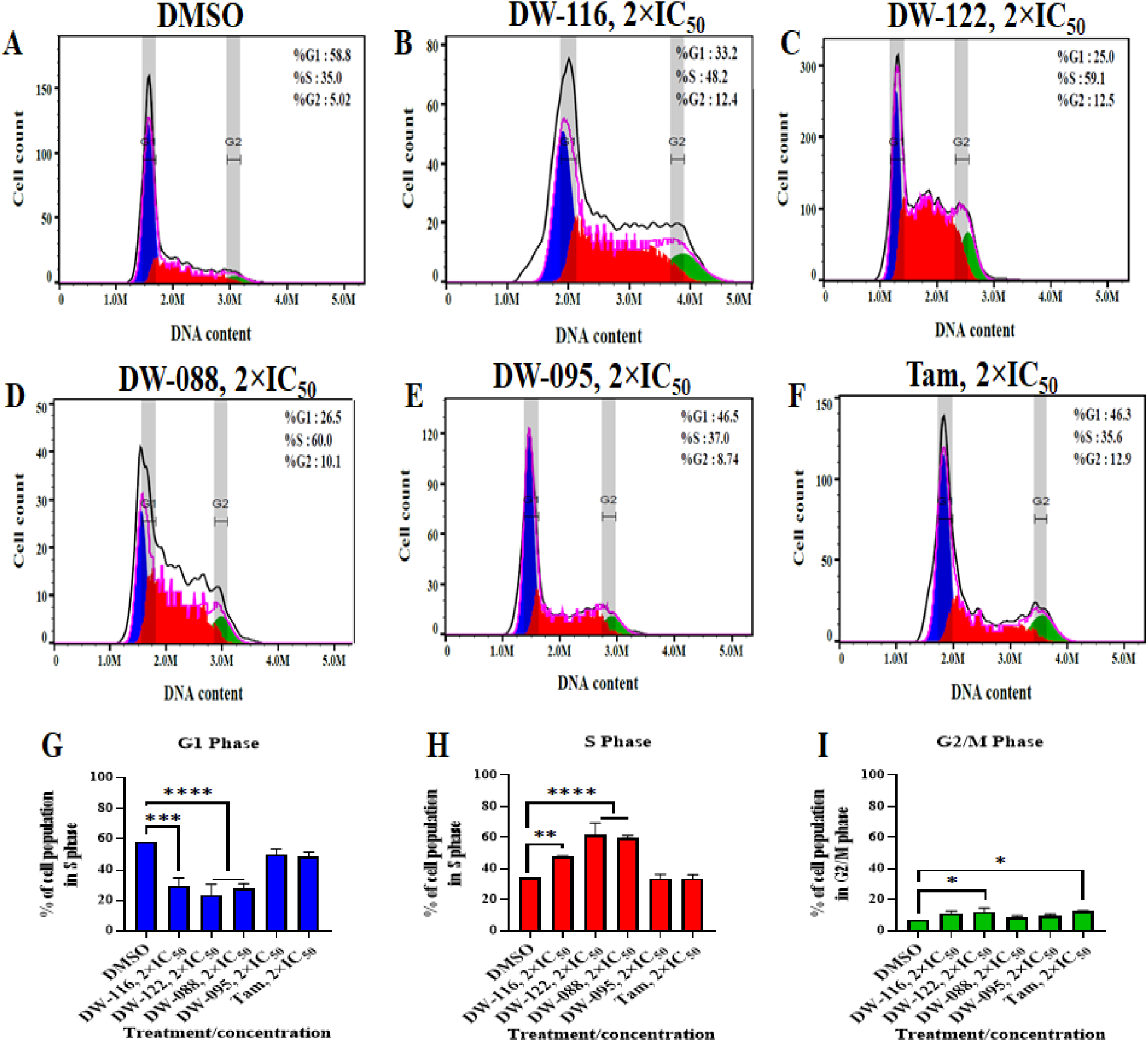
Effects of representative Tam-KDMi (**DW-116** and **DW-122**), EED-KDMi (**DW-088** and **DW-095**), and tamoxifen on cell cycle progression in MCF-7 cell line treated for 24 h at 2×IC_50_ concentrations of the respective compounds. Histograms of the negative control (DMSO, **A**) and cells treated with the indicated compounds display the distribution of cell populations in G0/G1, S, and G2/M phases of the cell cycle (**B**-**F**). The effects of treatment on the phases of the cell cycle are presented as bar charts (**G**-**I**) based on at least two independent experiments. \**p* < 0.05; \*\**p* < 0.01; \*\*\**p* < 0.001; \*\*\*\**p* < 0.0001.

**Figure 7:**
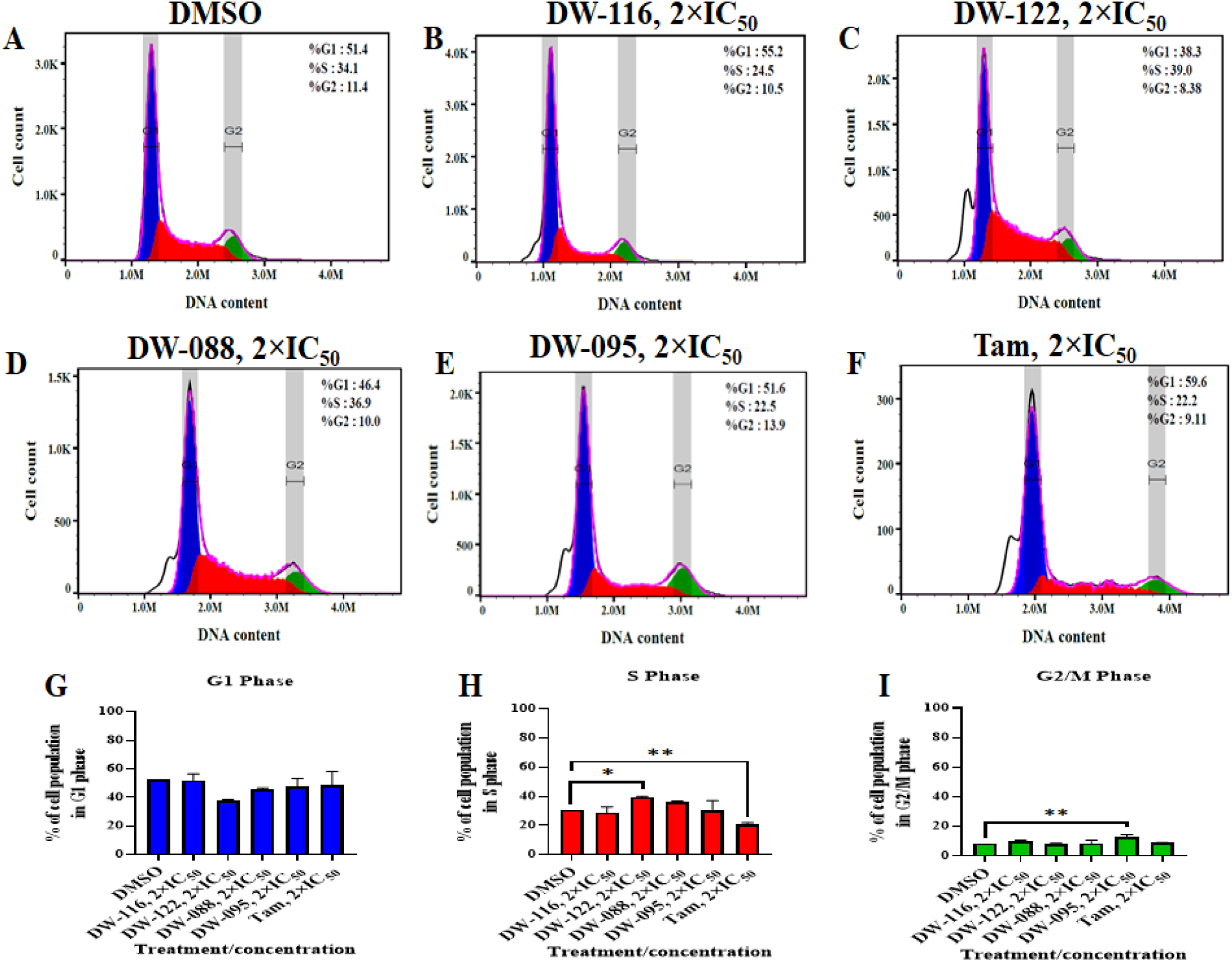
Effects of representative Tam-KDMi (**DW-116** and **DW-260**), EED-KDMi (**DW-088** and **DW-095**), and tamoxifen on cell cycle progression in MDA-MB-231 cell line treated for 24 h at 2×IC_50_ concentrations of the respective compounds. Histograms of the negative control (DMSO, **A**) and cells treated with the indicated compounds display the distribution of cell populations in G0/G1, S, and G2/M phases of the cell cycle (**B**-**F**). The effects of treatment on the phases of the cell cycle are further presented as bar charts (**G**-**I**) based on at least two independent experiments. \**p* < 0.05; \*\**p* < 0.01.

### 2.8 Tam-KDMi and EED-KDMi modulate cell proliferative pathways in MCF-7 and MDA-MB-231 cells

To further establish the mechanism(s) of the cytotoxic activity of the Cle-based KDMi, we used immunoblot assays to determine the effects of **DW-116**, **DW-122**, **DW-088,** and **DW-095** on the activities of key cancer cell-sustaining proliferative pathways. For instance, hypoxia inducible factor-1α (HIF-1α) plays vital roles in the sustenance of cancer cells under hypoxic conditions.^56^ In fact, it is considered a master regulator of oxygen homeostasis.^56^ We observed that the Tam-KDMi, **DW-116** and **DW-122,** downregulated the expression levels of HIF-1α in MCF-7 and MDA-MB-231 cells. However, the EED-KDMi, **DW-088,** and **DW-095** showed a tendency to increase the expression levels of HIF-1α; albeit a similar attribute has been noted for estradiol in MCF-7 cells. ^57^

Egl-9 family hypoxia inducible factor 3 (EGLN3 or PHD3), is a prolyl hydroxylase that senses oxygen levels in cells to regulate HIF-1α expression.^52-56^ We observed that EED does not affect EGLN3 levels in both cancer cell lines. DFP caused next to no effects on the expression levels of EGNL3 in MCF-7 cells, while it elicited downregulatory effects in MDA-MB-231 cells, as we have observed before. ^58^ All the tested Tam-KDMi and EED-KDMi influenced EGLN3 expression following the same trend as DFP (Fig. 8). Previous reports have shown that the downregulation of EGLN3 is associated with tumor aggressiveness and survival since EGLN3 normally acts as a tumor suppressor protein. ^57^

**Figure 8:**
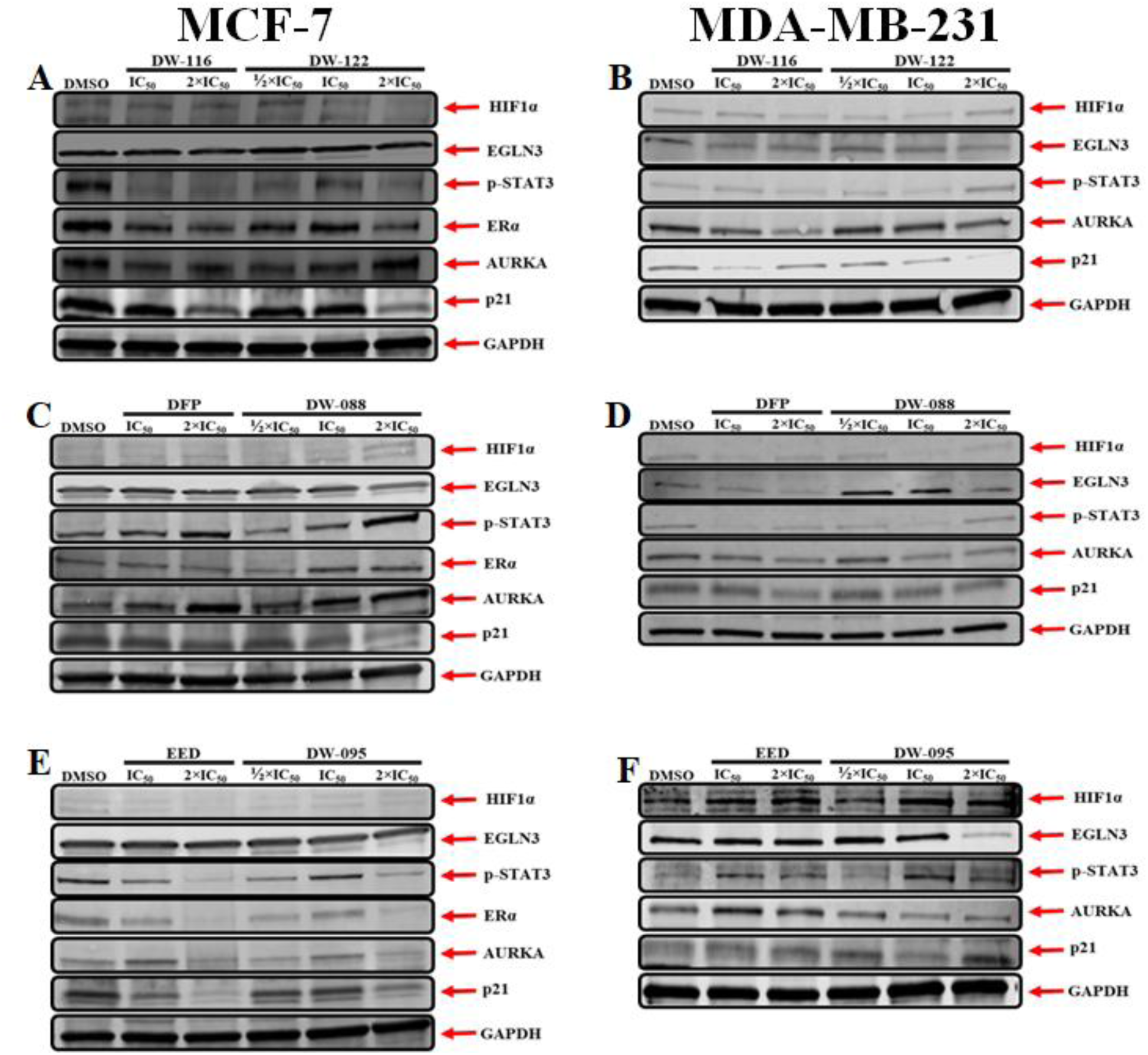
Western blot analysis of the expression of proteins involved in cell proliferation, cell motility, apoptosis and cell cycle regulation in MCF-7 and MDA-MB-231 cells treated with **DW-116**, **DW-122**, **DW-088**, **DW-095**, EED, and DFP. Cells were treated for 24 h and total cell extracts were subsequently analyzed by immunoblot analysis. Representative gel bands are shown above. The respective full gels and densitometric quantifications are presented in the Suppl info Figures S6-S17.

Phosphorylated signal transducer and activator of transcription-3 (p-STAT3) is relevant in cancer cell proliferation and metastasis, thus promoting tumor growth.^59^ We observed that both Tam-KDMi, **DW-116** and **DW-122**, elicited a significant tendency toward downregulating p-STAT3. In contrast, DFP upregulated the levels of p-STAT3 in MCF-7 cells, while it showed opposite effects in MDA-MB-231 cells (Fig. 8). This disparity in the effects of DFP may be an indication of cell-based differences. In both cell lines, we observed that EED caused a repression of p-STAT3 levels. However, EED-KDMi **DW-088** and **DW-095** did not have similar effects on the levels of p-STAT3 – **DW-088** and **DW-095** showed tendencies to respectively upregulate and downregulate p-STAT3 in both cell lines (Fig. 8). These differences between the two compounds may be among the factors responsible for the higher cytotoxicity **DW-095** in all the cell lines that we screened in this study.

Next, we probed for the effects of the tested compounds on ERα expression levels in MCF-7. We observed that DFP, EED, both Tam-KDMi **DW-116** and **DW-260,** and EED-KDMi **DW-095** downregulated the expression levels of ERα (Fig. 8). Paradoxically, due to unknown mechanisms, EED-KDMi **DW-088** displayed a tendency to upregulate ERα expression. Previous reports have demonstrated that estrogens can downregulate the expression of estrogen receptors in MCF-7 cells.^60-64^

Aurora kinase A (AURKA) plays vital roles in promoting the proliferation and survival of cancer cells, and its upregulation has been implicated in the induction of tamoxifen resistance in ERα(+) BCa.^63,64^ We observed that DFP and Tam-KDMi **DW-088** expressed differential effects on this protein in MCF-7 and MDA-MB-231 cells after 24 h of treatment – an upregulation in MCF-7 cells, while a downregulation in MDA-MB-231 cells. While EED downregulated AURKA in MCF-7 cells, it elicited no statistical effect on AURKA levels in MDA-MB-231 cells. Interestingly, we found that both EED-KDMi **DW-095** and Tam-KDMi **DW-122** showed a fairly similar trend in MCF-7 cells, downregulating AURKA in both cell lines at their respective 2× IC_50_ (Fig. 8).

Moreover, we evaluated the effects of the test compounds on the expression of cyclin-dependent kinase inhibitor 1 (CDKN1A/p21). Normally, p21 functions as a tumor suppressor by inducing cell cycle arrest and apoptosis. Depending on its location in a cell – either in the cytoplasm or nucleus – p21 can promote or prevent apoptosis, and its expression has been linked with promotion of cell survival through inhibition of apoptosis, mostly after DNA damage^.71^ We observed that all of the tested compounds caused downregulation of p21 expression in MCF-7 cells. The effects of these compounds on p21 expression in MDA-MB-231 cells are slightly different. EED elicited no statistically relevant effect on p21; DFP, **DW-122** and **DW-088** downregulated p21 while the downregulatory effects shown by **DW-095** at IC_50_ seemed to be annulled at its 2× IC_50_ (Fig. 8). Despite the diversity of their effects, it is likely that the perturbations caused by the tested Tam-KDMi and EED-KDMi on these oncogenic pathways/targets, tilted the balance in favor of cancer cell death.

### 2.9 RNA seq analysis revealed the effects of Tam-KDMi and EED-KDMi on the transcriptome

We used RNA-sequencing (RNA-seq) analysis to further probe the transcriptome-level effects of Tam-KDMi (**DW-260**) and EED-KDMi (**DW-088** and **DW-095**) in MCF-7 and MDA-MB-231 cells (Fig. 9). Hallmark gene set enrichment analysis (GSEA) was conducted first using the human molecular signature database (MSigDB) hallmark collection composed of 50 gene sets.^60^ Significance was defined as gene sets with a p-value < 0.05 and a false discovery rate (FDR) < 0.25 and enrichment was compared using normalized enrichment score (NES). In MCF-7 cells, both **DW-088** and **DW-260** at 2x IC_50_ positively enriched hypoxia, glycolysis, and mTORC1 gene sets similar to DFP, which we have shown to be a pan-selective KDMi.^53,58^ The effects on MDA-MB-231 cells are consistent with that of DFP, revealing that the E2F targets (NES = -2.2, **DW-088**, -3.7, **DW-095**, -2.0, **DW-260**) and G2M checkpoint (NES = -2.5, **DW-088**, -4.5, **DW-095**, -2.4, **DW-260**) gene sets were negatively enriched and the hypoxia gene set was positively enriched (NES = +2.3, **DW-088**, +2.4, **DW-095**, +2.6, **DW-260**) by all compounds at 2x IC_50_ while mitotic spindle was negatively enriched by **DW-088** (NES = -1.6) and **DW-095** (NES = -2.0) at 2x IC_50_.

**Figure 9:**
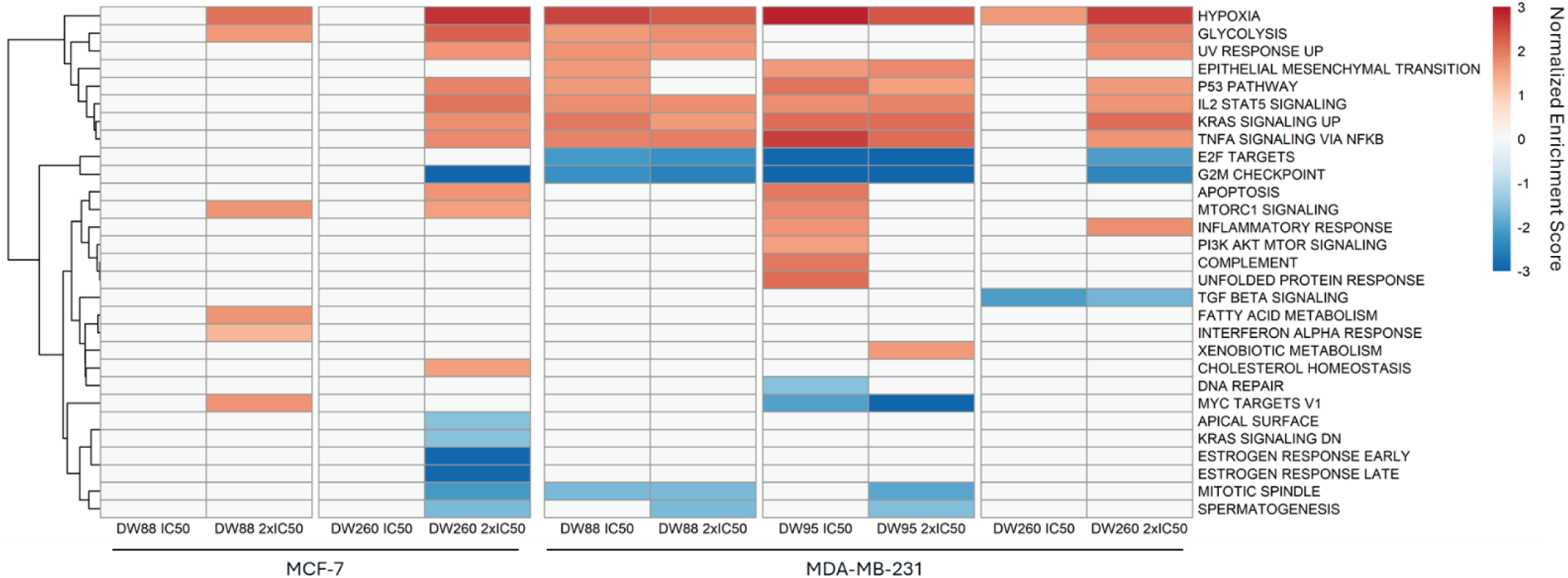
Hallmark Gene Set Enrichment Analysis (GSEA) of treatment to ERα+ (MCF-7) and triple-negative (MDA-MB-231) breast cancer cell lines. Heatmap of normalized enrichment scores (NES) of significantly enriched hallmark gene sets following resulting from EED-KDMi (**DW-088** and **DW-095**) and Tam-KDMi (**DW-260**) treatment to MCF-7 and MDA-MB-231 cells at IC_50_ and 2x IC_50_ (p < 0.05, FDR < 0.25). In MCF-7 cells, **DW-088** and **DW-260** at IC_50_ did not significantly enrich hallmark gene sets. Hypoxia (NES= +2.1, +2.8), glycolysis (NES= +1.7, +2.2), and mTORC1 signaling (NES= +1.7, +1.6) were positively enriched in MCF-7 cells by **DW-088** and **DW-260** at 2x IC_50_, respectively. IL-2-STAT5 signaling and KRAS signaling up (NES= +2.1, +1.8) were positively enriched by **DW-260** at 2x IC_50_ and estrogen response early and estrogen response late were negatively enriched by **DW-260** at 2x IC_50_ in MCF-7 cells. In MDA-MB-231 cells, hypoxia (NES= +2.3, +2.4, +2.6), IL-2-STAT5 signaling (NES= +1.7, +1.9, +1.7), KRAS signaling up (NES= +1.7, +2.1, +2.1), E2F targets (NES= -2.2, -3.7, -2.0), and G2M checkpoint (NES= -2.5, -4.5, -2.4) were enriched by **DW-088**, **DW-095**, and **DW-260** at 2x IC_50_, respectively. Mitotic spindle was negatively enriched by **DW-088** (NES= -1.6) and DW-095 (NES= -2.0) at 2x IC_50_.

Independently, IL-2-STAT5 signaling and KRAS signaling were positively enriched by **DW-088** (+1.7, +1.7), **DW-095** (+1.9, +2.1), and **DW-260** (+1.7, +2.1) in MDA-MB-231 and by **DW-260** in MCF-7 (+2.1, +1.8). Upregulation of genes in the IL-2-STAT5 and KRAS signaling pathways have been found to reduce metastasis in TNBC models and could contribute to the biological activity of these compounds.^65,66^ Additionally, both estrogen response gene sets were negatively enriched by Tam-KDMi **DW-260** (NES= -2.9, ER early, -2.9, ER late) in MCF-7 cells (Fig. 9).

Gene ontology biological processes (GOBP) gene sets significantly enriched (p < 0.01, FDR < 0.1) were determined by GSEA of the MSigDataBase GOBP collection. **DW-088**, **DW-095**, and **DW-260** negatively enriched cell cycle related gene sets including mitotic spindle organization (NES= -2.7, **DW-088**, -3.4, **DW-095**, -2.1, **DW-260**), chromosome organization (NES= -4.1, **DW-95**, -2.4, **DW-260**), and DNA recombination (NES= -1.9, **DW-088**, -2.2, **DW-95**, -2.2, **DW-260**) in MDA-MB-231 cells while minimal effect was seen in MCF-7 cells (Suppl info Fig S18).

Consistent with DFP data, HIF1A was downregulated and EGLNs 1 and 3, responsible for promoting HIF1A degradation, were upregulated in MCF-7 by **DW-260** (-1.1, +1.0, +2.0) and in MDA-MB-231 by **DW-088** (-2.1, +0.9, +3.0), **DW-95** (-2.1, +0.9, +2.4), and **DW-260** (-1.7, +1.1, +2.6) (Fig. 10).

**Figure 10:**
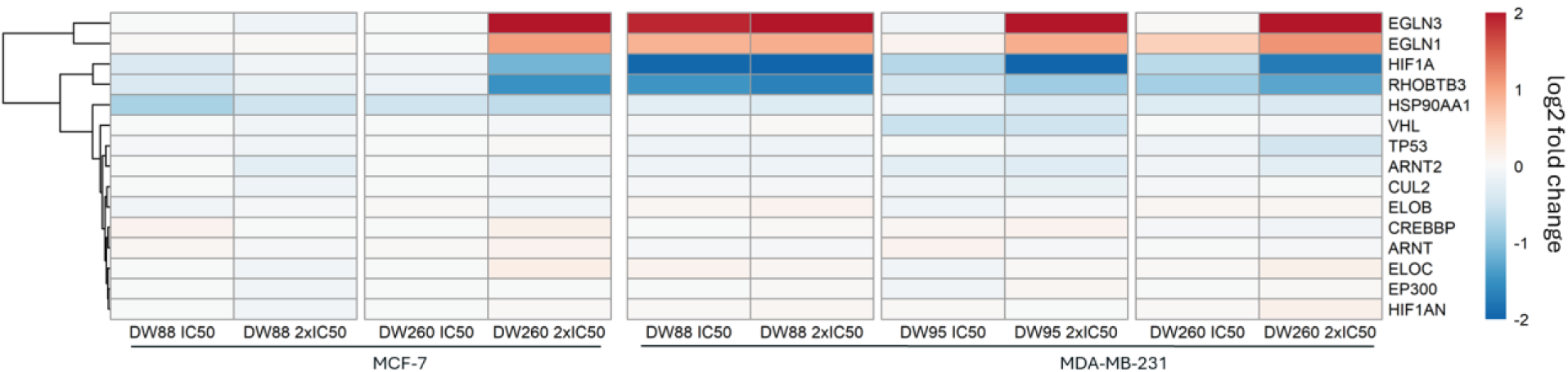
Effects of EED-KDMi (**DW-088** and **DW-095**) and Tam-KDMi (**DW-260**) on HIF1A and interacting genes in ERα+ (MCF-7) and triple-negative (MDA-MB-231) breast cancer cell lines. Log2 fold change heatmap of HIF1A interactors showing the regulation of HIF1A, EGLN1, EGLN3, and RHOBTB3 by **DW-260** 2x IC_50_ in MCF-7 and MDA-MB-231 cells and **DW-088** and **DW-095** in MDA-MB-231 cells. HIF1A downregulation at 2x IC_50_ by **DW-088** (log2 fold change= -2.1, MDA-MB-231), **DW-095** (log2 fold change= -2.1, MDA-MB-231), and **DW-260** (log2 fold change= -1.1, MCF-7; -1.7, MDA-MB-231). EGLNs 1 and 3 upregulation at 2x IC_50_ by **DW-088** (log2 fold change= +0.9, +3.0, MDA-MB-231), **DW-095** (log2 fold change= +0.9, +2.4, MDA-MB-231), and **DW-260** (log2 fold change= +1.0, +2.0, MCF-7; +1.1, +2.6, MDA-MB-231). RHOBTB3 downregulation at 2x IC_50_ by **DW-088** (log2 fold change= -1.7, MDA-MB-231), **DW-095** (log2 fold change= -0.8, MDA-MB-231), and **DW-260** (log2 fold change= -1.5, MCF-7; -1.3, MDA-MB-231).

Finally, analysis of genes implicated in KDM inhibition ^67^ revealed an upregulation of tumor suppressors CDKN1A (p21) and CDKN1C by **DW-088** (+2.5, +2.5) and **DW-095** (+2.1, +3.3) in MDA-MB-231 and by **DW-260** in both MCF-7 (+1.4, +0.2) and MDA-MB-231 (+2.8, +1.9) cell lines (Fig. 11). Cell cycle promoters CCNB1, AURKA, and CDCA7 were downregulated by **DW-088** (-1.9, -1.7, -1.3), **DW-95** (-2.2, -2.3, -2.6), and **DW-260** (-1.3, -1.2, -1.6) in MDA-MB-231 while **DW-260** also downregulated CCNB1 (-2.0), and AURKA (-1.4) in MCF-7 cells. It is worth noting that there are disparities between this RNA-seq data and the Western blot data described above. The lack of correlation between the RNA-seq and Western blot data is not unexpected as the two techniques use different cellular targets to measure gene products’ expression.^68,69^ Nevertheless, this RNA-seq data collectively indicates that the tested compounds induced transcriptome level changes that compromised the viability of MCF-7 and MDA-MB-231 cells.

**Figure 11:**
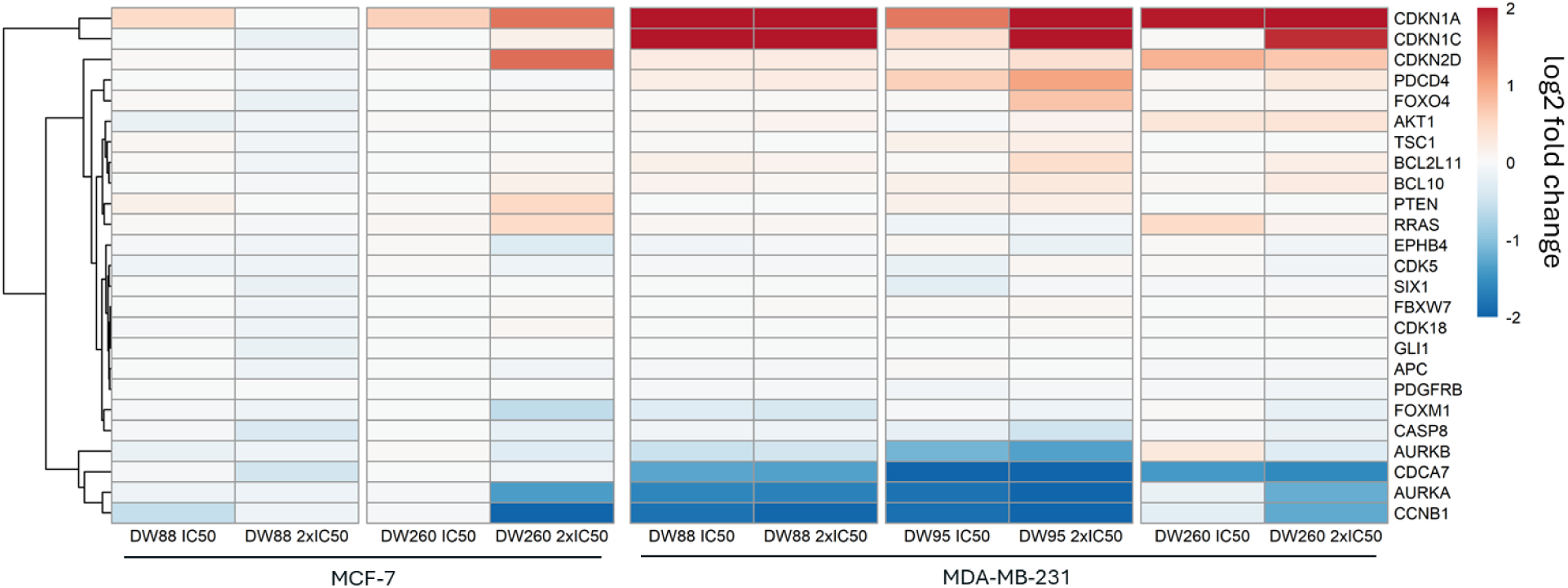
Effects of EED-KDMi (**DW-088** and **DW-095**) and Tam-KDMi (**DW-260**) on genes implicated in KDM inhibition in ERα+ (MCF-7) and triple-negative (MDA-MB-231) breast cancer cell lines. Upregulation of CDKN1A (p21) and CDKN1C by **DW-088** (log2 fold change= +2.5, +2.5) and **DW-095** (log2 fold change= +2.1, +3.3) at 2x IC_50_ in MDA-MB-231 cells and by **DW-260** at 2x IC_50_ in MCF-7 (log2 fold change= +1.4, +0.2) and MDA-MB-231 (log2 fold change= +2.8, +1.9). Downregulation of CCNB1, AURKA, and CDCA7 at 2x IC_50_ by **DW-088** (log2 fold change= -1.9, -1.7, -1.3, MDA-MB-231), **DW-095** (log2 fold change= -2.2, -2.3, -2.6, MDA-MB-231), and **DW-260** (log2 fold change= -1.3, -1.2, -1.6, MDA-MB-231). Downregulation of CCNB1 and AURKA by **DW-260** at 2x IC_50_ in MCF-7 (log2 fold change= - 2.0, -1.4).

### 2.10 In vivo efficacy of Tam-KDMi and EED-KDMi in murine xenograft models of BCa

Encouraged by the *in vitro* data described above demonstrating their on-target effects and cytotoxicity to both ERα(+) and ERα(-) BCa cells, we investigated the antitumor potential of Tam-KDMi (**DW-116**) and EED-KDMi (**DW-088** and **DW-095**) in four murine models – one TNBC (MDA-MB-231) and two ERα(+) (T47D and ZR75.1) xenograft models; and one syngeneic (AT3) mouse model – reflective of various clinical presentations of BCa. For this study, we used T47D and MDA-MB-231 models to test the *in vivo* efficacy of **DW-116,** while **DW-088** and **DW-095** were evaluated against ZR75.1 and MDA-MB-231 xenograft and AT3 syngeneic models. Xenografts were established in 8-week-old mice, while we used 12-week-old mice for the syngeneic model. Treatment commenced after tumors reached ∼250 mm^3^ and test compounds were injected intraperitoneally (i.p) at 25 mg/kg, three times a week, for 6 weeks. The control animals received vehicle only, and tumor volume was measured once a week for 6 weeks. We observed that these compounds drastically and robustly reduce tumor growth in all of these models, with tumor growth inhibition (TGI) as high as 70% relative to the untreated control (Fig. 12). More specifically, Tam-KDMi **DW-116** inhibited tumor growth in both ERα(+) and ERα(-) models but its tumor growth inhibition effect is more pronounced against the ERα(+) T47D model (Fig. 12a). In contrast, the EED-KDMi **DW-088** and **DW-095** showed near equal tumor growth inhibition in both ERα(+) and ERα(-) models with a slightly improved efficiency against the ERα(-) model (Fig. 12b-c). Interestingly, **DW-088** and **DW-095** are more efficacious against the syngeneic AT3 model, showing near total tumor growth inhibition (Fig. 12d). This *in vivo* efficacy data illustrated several important qualities of the Tam-KDMi and EED-KDMi. First, **DW-116** is a unique tamoxifen-based agent that shows efficacy in both ER (+) and ER(-) BCa models. Also, AT3 is a mouse cell line derived from an autochthonous tumor considered to model human tumors more closely than transplanted tumors and a useful model for tumor physiology that is representative of typical disease states in humans.^70^ The enhancement of the efficiency of **DW-088** and **DW-095** in the syngeneic AT3 model is strongly indicative of their potential in other clinically relevant models with intact immune systems. Overall, this data provides clear evidence of the potential of Tam-KDMi and EED-KDMi as therapeutic agents for BCa regardless of the tumor’s receptor expression status.

**Figure 12:**
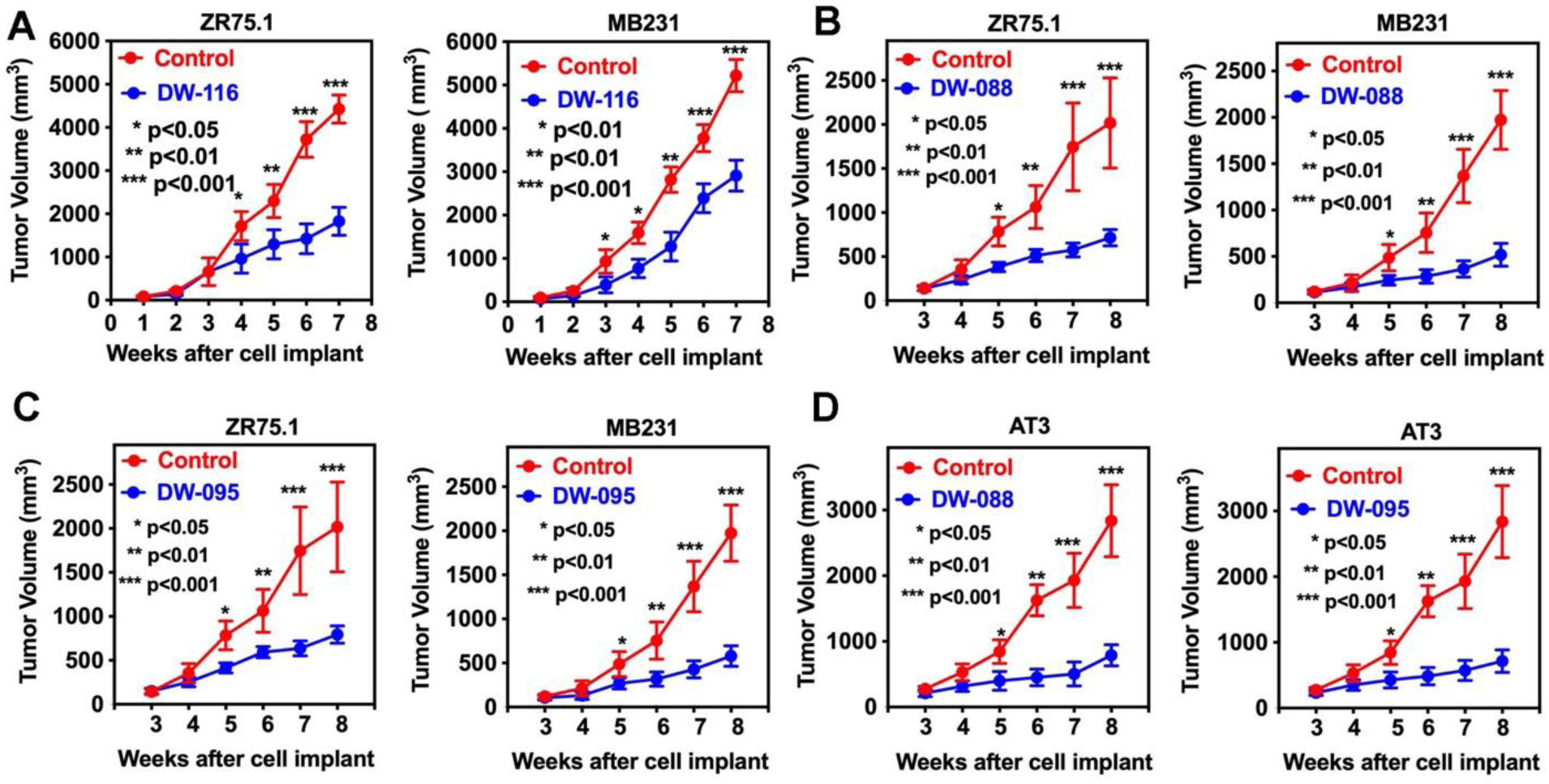
I*n vivo* efficacy of Tam-KDMi **DW-116**, EED-KDMi **DW-088,** and **DW-095** in mouse xenografts and syngeneic models. Test agents were administered intraperitoneally (i.p) at 25 mg/kg. (A) Effects of **DW-116** on tumor growth in ZR75.1 and MDA-MB-231 xenografts. (B-C) Effects of **DW-088** and **DW-095** on tumor growth in ZR75.1 and MDA-MB-231 xenografts. (D) Effects of **DW-088** and **DW-095** on tumor growth in the syngeneic AT3 model. Shown in the Y-axis are tumor volumes (mm3), and on the X-axis are weeks after tumor implant.

## 3. Conclusion

Two vital mechanisms for the sustenance of breast carcinoma are epigenetic dysfunction and the malfunction of endocrine proteins such as ERα. We demonstrated herein that the integration of ERα antagonist (Tam) and agonist (EED) ligands with DFP-derived KDM inhibitor moiety furnished dual-acting agents Tam-KDMi and EED-KDMi, respectively. These agents showed robust on-target effects and potent BCa cells-selective anti-proliferative activities. We observed that simultaneous inhibition of ER signaling and KDM by Tam-KDMi resulted in cytotoxicity to both ERα(+) and ERα(-) BCa cells, while the lack of ER-signaling inhibition conferred an enhanced MDA-MB-231 cells cytotoxic to the EED-KDMi. Moreover, the exemplary compounds Tam-KDMi **DW-116** and EED-KDMi **DW-088** and **DW-95** significantly reduce tumor growth in murine xenograft models of ERα(−) and ERα(+) BCas with TGI as high as 70%. Collectively, **DW-116**, **DW-088** and **DW-95** have a high potential as leads for the development of new agents for the treatment of BCa subtypes regardless of the tumor ER expression status.

## 4. Method and materials

Anhydrous solvents and reagents, high-performance liquid chromatography (HPLC) grade or American Chemical Society (ACS) grade solvents were purchased from Sigma-Aldrich (St. Louis, MO, USA), Across, VWR International (Radnor, PA, USA), or Thermo Fisher Scientific (Waltham, MA, USA) and were used without further purification. Analytical TLC was conducted using Analtech silica gel plates (60 F254), while purification was achieved using Analtech preparative TLC plates (UV 254, 2000 µm). Spot visualization was facilitated using UV light in conjunction with anisaldehyde/iodine stain. For column chromatography, 200–400 Mesh silica gel was utilized. HPLC analyses of products were carried out using Phenomenex Luna 91 5 µm C8(2) 100 Å LC column (4.6×250 mm) using Agilent 1260 Infinity II HPLC system at detection wavelength of 254 nm. Water (solvent A) and MeCN (solvent B) with 0.1 % TFA were used as the mobile phase at a flow rate of 0.5 mL·min^-1^. The chromatography elution profile was as follows: 5% solvent B from 0 to 5 min, linear gradient to 100% solvent B from 5 to 18 min, 100% solvent B from 18 to 22 min, linear gradient to 5% solvent B from 22 to 24 min, and 5% solvent B from 24 to 30 min. Sample concentrations were 250 μM – 1mM, injecting 30 μL. Nuclear magnetic resonance (NMR) spectra were acquired using Varian-Gemini 400 MHz, Bruker 500 MHz, or 700 MHz magnetic resonance spectrometers. ^1^H NMR spectra were referenced in parts per million (ppm) relative to the residual peaks of CHCl_3_ (7.24 ppm) in CDCl_3_, CHD_2_OD (4.78 ppm) in CD_3_OD, or DMSO-*d5* (2.49 ppm) in DMSO-*d6*. Similarly, ^13^C spectra were referenced relative to the central peak of the CDCl_3_ triplet (77.0 ppm), CD_3_OD septet (49.3 ppm), or DMSO-*d6* septet (39.7 ppm), employing complete hetero decoupling. Data from the ‘fid’ file were processed using MestReNova LITE (version 5.2.5−5780) software. High-resolution mass spectra were recorded at the mass spectrometry facility of the Georgia Institute of Technology in Atlanta

### Cell culture

All cell lines were obtained from the American Type Culture Collection. HepG2, SK-HEP-1, DU145 were cultured in MEM (Corning) supplemented with L-glutamine; MCF-7 was cultured in DMEM (Corning) without phenol red supplemented with L-glutamine; MDA-MB-231, A549, and Vero were cultured in DMEM (Corning), while LNCaP was cultured using RPMI-1640 supplemented with L-glutamine. All media were supplemented with 10% FBS and 1% penicillin/streptomycin. mES CiA cells were cultured in DMEM (Corning) with 15% FBS supplemented with 100 units/mL penicillin/streptomycin, non-essential amino acids (NEAA; Gibco 11140-050), 10 mM HEPES buffer (Corning, 25-060-CI), 55 µM β-mercaptoethanol, leukemia inhibitory factor (LIF), 7.5µg/mL blasticidin (InvivoGen, ant-bl-1), and 1.5 µg/mL puromycin (InvivoGen, ant-pr-1) on gelatin-coated tissue culture treated plates.

### ER-binding Assay

Estrogen receptor competitive binding of the tested compounds was assessed with PolarScreen™ Estrogen Receptor Alpha Competitor Assay Kit, Green (#A15883, Thermo Scientific), based on the manufacturer’s instructions.

### MTT cell viability assay

Using standard protocol, with minor modifications, cells, at a density of 4.5 × 10^3^/well and a medium volume of 100 μL, were seeded in 96-well transparent tissue culture plates and allowed to adhere for 24 h. Thereafter, the culture medium was aspirated, and the cells were treated for 72 h with 100 μL of varying concentrations (0.5 – 100 μM) of the compounds, dissolved in dimethyl sulfoxide (DMSO) but diluted with the respective culture media to ensure a final maximum DMSO concentration of 1%. Subsequently, 10 μL of MTT reagent (5 mg/mL) was added to the culture medium, and the plates were incubated for 3 h, after which the MTT reagent/medium was carefully aspirated. Finally, the resulting formazan crystals were dissolved in 100% DMSO (100 μL/well), and absorbance values were determined at 570 nm using multimode plate reader (Tecan Infinite M200 Pro, Männedorf, Switzerland), and the percent cell viability was computed with respect to untreated controls. IC_50_ values were determined using Prism GraphPad 10.

### Clonogenic assay

Clonogenicity was determined using the methods of Liu et al.^46^ and Franken et al.^45,^ with some modifications. Cells were seeded in 12-well plates at a density of 200 - 500 cells per well (in 1.0/1.5 mL media) to avoid contact within the expected clones, and the cells were allowed to adhere for 24 h. Subsequently, cells were treated as indicated and incubated for 72 h. Thereafter, the treated culture media were replaced with fresh media to allow cells to grow for 10 days, with media being changed every third day. Afterward, the plate was washed twice with sufficient PBS, fixed with 400 µL paraformaldehyde (4%) at 37°C for 20 min (alternatively, the colonies were fixed with ice-cold 100% methanol for 5 - 10 min), aspirated, and stained with 0.5 - 1 mL crystal violet (0.1%, Sigma Aldrich, St. Louis, MO) for 20 min after aspirating the fixative. Excess crystal violet solution was washed thrice with dH_2_O, and the fixed cells were allowed to dry for at least 24 h. Colonies were scored using a stereomicroscope and representative images taken for quantification using ImageJ software (National Institutes of Health). For further quantitative analysis^46^, 1.5 mL 33% acetic acid was added to each well, and the plate was shaken for 1 h. Then, 100 µL was taken from each well to a 96-well plate for absorbance reading at 560 nm using a microplate reader (TECAN mPro200, USA).

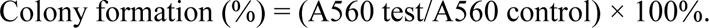

### Chromatin *in vivo* assay

Mouse embryonic stem cells (mES) with an *Oct4* allele replaced with eGFP were modified via lentiviral infection to stably contain the chromatin *in vivo* assay (CiA) components consisting of plasmids N118 (LV EF-1α-Gal-FKBPx1-HA-PGK Blast) and N163 (nLV EF-1α-HP1α (CS)-Frbx2(Frb+FrbWobb)-V5-PGK-Puro), as previously described. The mES CiA cells were then seeded in gelatin-coated 96-well plates at a density of 10,000 cells per well. The following day, the cell culture media were removed and replaced with media containing either a DMSO vehicle, 6 nM rapamycin, and a DMSO vehicle, or 6 nM rapamycin and a varied concentration of a Tam-KDMi or EED-KDMi compound dissolved in DMSO; vehicle-only and rapamycin with vehicle were considered as positive and negative controls, respectively. Concentrations of EED-KDMi were varied between 1-10 µM per well, except **DW-554** (which was dosed at 1, 5, and 10 µM). Concentrations of Tam-KDMi were varied between 0.25-1 µM. All conditions, including controls, were measured using biological triplicates. After 48 h of incubation, the media was removed and cells were collected via trypsinization with 0.25% trypsin-EDTA, quenches with media and PBS, and then transferred to a 96-well untreated U-bottom plate. Cell fluorescence was then analyzed using an Attune NxT Acoustic Focusing Flow Cytometer with an autosampler (ThermoFisher) using a 488 nM laser with a 530/30 filter for GFP signal and a 637 nM laser with a 670/14 filter for autofluorescence. Observed mES cells were gated to exclude dead, autofluorescent, or conjoined cells, and were then further gated based on GFP fluorescence using a bifurcating gate to gate to generate two populations expressed as percentages (%GFP+ and %GFP-). A mean %GFP+ value was found for each control condition. Then, a % inhibition value was generated for each sample well containing rapamycin and a concentration Tam-KDMi or EED-KDMi compound; % inhibition was defined as 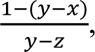 where x is the %GFP+ of the well, y is the mean %GFP+ of the positive control, and z is the mean %GFP+ of the negative control.

### Cell cycle analysis

This analysis was carried out by adapting the manufacturer’s instructions (Canvax Biotech, S.L., Spain) with some modifications. Briefly, cells (1 × 10^6^ cells/well in 2 mL of medium) were seeded in six-well plates and allowed to attach for 24 h before treatment for another 24 h. Subsequently, cells were dissociated from the substrate using a cell scrapper, and the cell suspension was centrifuged at 500 ×g for 5 min at 4^0^C, with the supernatant aspirated and discarded. Cells were then washed in 1 mL ice-cold PBS and centrifuged again with the supernatant aspirated and discarded. Next, nucleic acid labeling was initiated by initially fixing the cells with 1 mL ice cold 70% ethanol added dropwise to the cell pellet while vortexing and the samples were stored on ice for at least 30 min. Thereafter, without disrupting the pellet, ethanol was carefully removed by centrifugation at 2500 ×g for 5 min at 4°C, and the cells were subsequently washed in 1 mL PBS. Again, PBS was removed by centrifugation at 2500 ×g for 5 min, and then the cells were resuspended in 200 μL of the staining solution (freshly prepared by mixing PBS with propidium iodide and RNase A in a ratio of 50:1:1, respectively), which should be protected from exposure to light. In preparation for flow cytometry analysis (using CytoFLEX S Flow Cytometer), the tubes containing the stained cells were incubated in the dark for 30 min at 37°C and finally placed on ice before analysis.

### Western blot

Cells were seeded into 6-well plates at a density of 1 x 10^^6^ cells per well and allowed to incubate for 24 h. Following treatment for 24 h, cells were lysed using radioimmunoprecipitation assay (RIPA) buffer (150 μL) supplemented with phosphatase and protease inhibitors on ice for 15 min. The lysates were sonicated for 90 s and then centrifuged at 16000 ×*g* for 10 min, and the supernatants were collected. Total protein concentrations were determined using a BCA protein assay kit. Based on the protein concentrations, the lysates were normalized to achieve equal protein concentrations; the denatured lysate was loaded onto TGX MIDI 4–15% gels and electrophoresed at 150 V for 70 min. Subsequently, the gel was transferred onto a Turbo PDVF membrane, blocked with 5% BSA, and incubated overnight at 4 °C with the desired primary antibodies. On the following day, the membrane was washed with TBS-T, incubated with secondary antibodies. After removal of the secondary antibodies, and membrane washing steps, images of the immunoblots were scanned with LI-COR’s Odyssey CLx® Imager (software Image Studio™ software v. 5.2) and the signal intensities were quantified using ImageJ software.

### Synthesis procedure and characterization

#### Procedure for the Synthesis of ((4-methoxybenzyl)oxy)-2-methyl-4H-pyran-4-one (PMB-maltol) **(1)**

Maltol (10 g, 1 equiv.) was dissolved in dry acetone (100 mL) under an argon atmosphere. To this mixture, K_2_CO_3_ (16.42 g, 1.5 equiv.) was added. The reaction mixture was stirred at room temperature for 15 min. Then, paramethoxybenzyl bromide (12.72 mL, 1.1 equiv.) was added dropwise. The mixture was then heated under reflux for 12 h. After the reaction was complete, as indicated by TLC, the mixture was quenched with water (300 mL) and extracted with dichloromethane (CH_2_Cl_2_) (2 × 200 mL). The organic phase was dried over Na_2_SO_4_ and then filtered. The solvent was removed using a rotary evaporator, and the crude material was purified by column chromatography using a mixture of methanol and dichloromethane (1:10, v/v), affording compound **1** in 16.85 g (86%) as a brown liquid. ^1^H NMR (700 MHz, CDCl_3_) δ 7.57 (d, J = 5.6 Hz, 1H), 7.29 (d, J = 8.6 Hz, 2H), 6.84 (d, J = 8.7 Hz, 2H), 6.33 (d, J = 5.6 Hz, 1H), 5.08 (s, 2H), 3.78 (s, 3H), 2.04 (s, 3H). ^13^C NMR (176 MHz, CDCl_3_) δ 175.2, 159.9, 159.7, 153.4, 143.6, 130.8, 129, 117.1, 113.8,73.1, 55.3, 14.8.

#### General procedure for the synthesis of PMB-Maltol-Azides 3 (a-g)

A mixture of corresponding azido amines (1.5 equiv.), PMB-maltol (1 equiv.) and sodium hydroxide (0.5 equiv.) was prepared in a mixture of ethanol and water (3:2). This mixture was heated in a sealed tube at 105 °C for 24 h. The completion of the reaction was monitored using thin-layer chromatography (TLC). The mixture was quenched with 50 mL of water and then extracted with dichloromethane (twice, 20 mL each). The organic phases were combined, dried over sodium sulfate (Na_2_SO_4_), and filtered. The solvent was removed using a rotary evaporator, and the crude material was further purified using silica gel column chromatography, eluting with a 1:12 (v/v) mixture of MeOH and CH_2_Cl_2_, which afforded the desired PMB-Maltol-Azide compounds **3 (a-g)** as brown liquid.

#### 1-(2-Azidoethyl)-3-((4-methoxybenzyl) oxy)-2-methylpyridin-4(1H)-one (**3a)**

Following the general procedure using ((4-methoxybenzyl) oxy)-2-methyl-4H-pyran-4-one (**1**) (500 mg), 2-azidoethan-1-amine **2a** (262 mg), and NaOH (40.6 mg) gave **3a** (312 mg, 48 % yield) as a brown liquid. ^1^H NMR (400 MHz, CDCl_3_) δ 7.30 (d, *J* = 8.8 Hz, 2H), 7.20 (d, *J* = 7.5 Hz, 1H), 6.84 (d, *J* = 8.7 Hz, 2H), 6.43 (d, *J* = 7.6 Hz, 1H), 5.16 (s, 2H), 3.87 (t, *J* = 5.7 Hz, 2H), 3.79 (s, 3H), 3.52 (t, *J* = 5.7 Hz, 2H), 2.05 (s, 3H). ^13^C NMR (176 MHz, CDCl_3_) δ 173.8, 159.6, 146, 140.5, 138.5, 131, 129.7, 117.6, 113.7, 72.7, 55.3, 52.2, 51.1, 29.8, 12.7.

#### 1-(3-Azidopropyl)-3-((4-methoxybenzyl)oxy)-2 methylpyridin-4(1H)-one **(3b)**

Following the general procedure using ((4-methoxybenzyl) oxy)-2-methyl-4H-pyran-4-one **(1)** (500 mg), 3-azidopropan-1-amine **2b** (304 mg), and NaOH (40.6 mg) gave **3b** (362 mg, 54 % yield) as a brown liquid. ^1^H NMR (400 MHz, CDCl_3_) δ 7.28 (d, J = 8.8 Hz, 2H), 7.18 (d, J = 7.5 Hz, 1H), 6.82 (d, J = 7.8 Hz, 2H), 6.40 (d, J = 7.5 Hz, 1H), 5.13 (s, 2H), 3.82 (d, J = 6.2 Hz, 2H), 3.78 (s, 3H), 3.30 (t, J = 6.6 Hz, 2H), 2.04 (s, 3H), 1.87 – 1.79 (m, 2H). ^13^C NMR (176 MHz, CDCl_3_) δ 173.4, 159.5, 140.8, 130.9, 129.7, 129.2, 128.3, 72.5, 55.3, 50.5, 47.6, 29.8, 12.4.

#### 1-(4-Azidobutyl)-3-((4-methoxybenzyl)oxy)-2-methylpyridin-4(1H)-one **(3c)**

Following the general procedure using ((4-methoxybenzyl) oxy)-2-methyl-4H-pyran-4-one **(1)** (500 mg), 4-azidobutan-1-amine **2c** (347 mg), and NaOH (40.6 mg) gave **3 c** (391 mg, 56 % yield) as a brown liquid. ^1^H NMR (700 MHz, CDCl_3_) δ 7.30 (d, *J* = 6.2 Hz, 2H), 7.16 (d, *J* = 7.4 Hz, 1H), 6.83 (d, *J* = 8.8 Hz, 2H), 6.40 (d, *J* = 4.0 Hz, 1H), 5.16 (s, 2H), 3.78 (s, 3H), 3.75 (t, 2H), 3.31 (t, J = 5.9 Hz, 2H), 2.05 (s, 3H), 1.73 – 1.65 (m, 2H), 1.55 – 1.47 (m, 2H). ^13^C NMR (176 MHz, CDCl_3_) δ 173.5, 146, 140.7, 138, 130.9, 129.8, 117.4, 113.6, 72.5, 55.3, 53.2, 50.8, 28.1, 25.8, 12.5.

#### 1-(5-Azidopentyl)-3-((4-methoxybenzyl)oxy)-2-methylpyridin-4(1H)-one **(3d)**

Following the general procedure using ((4-methoxybenzyl) oxy)-2-methyl-4H-pyran-4-one **(1)** (500 mg), 5-azidopentan-1-amine (**2d**) (390 mg), and NaOH (40.6 mg) gave **3d** (407 mg, 56 % yield) as a brown liquid. ^1^H NMR (400 MHz, CDCl_3_) δ 7.31 (d, J = 8.7 Hz, 2H), 7.16 (d, J = 7.5 Hz 1H), 6.84 (d, J = 8.8 Hz, 2H), 6.41 (d, J = 7.5 Hz, 1H), 5.16 (s, 2H), 3.79 (s, 3H), 3.74 (t, 2H), 3.29 (t, J = 6.6 Hz, 2H), 2.06 (s, 3H), 1.69 – 1.55 (m, 4H), 1.40 – 1.31 (m, 2H). ^13^C NMR (176 MHz, CDCl_3_) δ 173.4, 146, 140.7, 138.1, 130.9, 117.3, 113.6, 72.5, 53.6, 50.9, 30.4, 29.7, 28.5, 23.6, 12.5.

#### 1-(6-Azidohexyl)-3-((4-methoxybenzyl)oxy)-2-methylpyridin-4(1H)-one **(3e)**

Following the general procedure using ((4-methoxybenzyl) oxy)-2-methyl-4H-pyran-4-one **(1)** (500 mg), 6-azidohexan-1-amine **2e** (432 mg), and NaOH (40.6 mg) gave **3e** (419 mg, 55 % yield) as a brown liquid. ^1^H NMR (400 MHz, CDCl_3_) δ 7.30 (d, *J* = 8.8 Hz, 2H), 7.18 (d, *J* = 7.6 Hz, 1H), 6.82 (d, *J* = 7.8 Hz, 2H), 6.46 (d, *J* = 7.6 Hz, 1H), 5.15 (s, 2H), 3.78 (S, 3H), 3.72 (d, *J* = 4.1 Hz, 2H), 3.26 (t, *J* = 6.7 Hz, 2H), 2.05 (s, 3H), 1.66 – 1.53 (m, 4H), 1.33 (m, 4H). ^13^C NMR (176 MHz, CDCl_3_) δ 173.3, 159.6, 145.9, 141, 138.1, 130.9, 129.8, 117.2, 113.7, 72.6, 55.3, 53.8, 51.2, 30.7, 28.7, 26.4, 26.

#### 1-(7-Azidoheptyl)-3-((4-methoxybenzyl)oxy)-2-methylpyridin-4(1H)-one **(3f)**

Following the general procedure using ((4-methoxybenzyl) oxy)-2-methyl-4H-pyran-4-one **(1)** (500 mg), 7-azidoheptan-1-amine **2f** (475 mg), and NaOH (40.6 mg) gave **3f** (404 mg, 51 % yield) as a brown liquid. ^1^H NMR (400 MHz, CDCl_3_) δ 7.31 (d, *J* = 5.8 Hz, 2H), 7.18 (d, *J* = 6.8 Hz, 1H), 6.83 (d, *J* = 7.4 Hz, 2H), 6.45 (d, *J* = 7.0 Hz, 1H), 5.16 (s, 2H), 3.79 (s, 3H), 3.72 (t, *J* = 7.2 Hz, 2H), 3.26 (t, *J* = 6.8 Hz, 2H), 2.05 (s, 3H), 1.66 – 1.52 (m, 4H), 1.32 (m, 6H). ^13^C NMR (176 MHz, CDCl_3_) δ 173.4, 159.6, 146, 140.9, 138.1, 130.9, 129.9, 117.3, 113.6, 72.6, 53.9, 51.4, 30.8, 28.8, 26.3, 12.6.

#### 1-(8-Azidooctyl)-3-((4-methoxybenzyl)oxy)-2-methylpyridin-4(1H)-one **(3g)**

Following the general procedure using ((4-methoxybenzyl) oxy)-2-methyl-4H-pyran-4-one **(1)** (500 mg), 8-azidooctan-1-amine **2g** (518 mg), and NaOH (40.6 mg) gave **3g** (392 mg, 48 % yield) as a brown liquid ^1^H NMR (700 MHz, CDCl_3_) δ 7.28 (d, *J* = 8.6 Hz, 2H), 7.14 (d, *J* = 7.5 Hz, 1H), 6.80 (d, *J* = 8.7 Hz, 2H), 6.36 (d, *J* = 7.5 Hz, 1H), 5.12 (s, 2H), 3.75 (s, 3H), 3.68 (t, *J* = 7.6 Hz, 2H), 3.22 (t, *J* = 6.9 Hz, 2H), 2.02 (s, 3H), 1.61 – 1.51 (m, 4H), 1.34 – 1.18 (m, 8H). ^13^C NMR (176 MHz, CDCl_3_) δ 173.3, 145.9, 140.7, 138.1, 117.2, 113.5, 72.4, 55.8, 30.7, 29, 28.9, 28.7, 28.7, 12.4.

#### Procedure for the synthesis of 4-(4-Ethynylbenzyl)−Tamoxifen **(5**)

**Step I –** N-Desmethyltamoxifen: The synthetic procedure was adapted from our previously established protocol.^29^ Briefly, to a solution of tamoxifen (1 g, 2.32 mmol) in 1,2-dichloroethane (15 mL) at 0 °C was added 1-chloroethyl chloroformate (332 mg, 2.32 mmol). Stirring continued at 0 °C for 15 min, followed by heating under reflux for 24 h. TLC revealed a near quantitative conversion to a higher Rf intermediate. The solvent was evaporated off, and the oily residue was dissolved in dry methanol. The mixture was then refluxed for 3 h, during which a reconversion to a lower Rf product was observed. The solvent was evaporated to give 0.910 g (94 %) of N-desmethyltamoxifen as a white solid.

**<u>Step-II –</u>** <u>Alkylation of N-desmethyltamoxifen</u>: To a stirring solution of N-desmethyltamoxifen (0.9 g, 1 equiv.) and 1-(bromomethyl)-4-ethynylbenzene (0.513 g, 1.05 equiv.) in CH_2_Cl_2_ (10 mL), Hunig’s base (1.283 mL, 3 equiv.) was added and stirring continued for 15 h. The completion of the reaction was indicated by TLC. The reaction mixture was quenched with 10% NaHCO_3_ solution (30 mL) and extracted with CH_2_Cl_2_ (3×30 mL). The organic layer was washed with saturated brine (1 × 30 mL) and dried over Na_2_SO_4_. Solvent was evaporated off, and crude material was purified on silica gel chromatography, eluting with ethyl acetate: hexanes (2:1), v/v, to afford to give 0.98 g (82%) 4-(4-ethynylbenzyl)−Tamoxifen **(5)** as a white−brown solid. ^1^H NMR (700 MHz, CDCl_3_) δ 7.46 (s, 2H), 7.40 – 7.37 (m, 2H), 7.32 – 7.27 (m, 5H), 7.23 – 7.19 (m, 2H), 7.17 (d, J = 9.8 Hz, 3H), 6.81 (d, J = 6.7 Hz, 2H), 6.58 (d, J = 6.7 Hz, 2H), 3.99 (t, J = 5.9 Hz, 2H), 3.61 (s, 2H), 3.09 (s, 1H), 2.78 (t, J = 5.9 Hz, 2H), 2.50 (q, J = 7.5 Hz, 2H), 2.31 (s, 3H), 0.97 (t, J = 7.5 Hz, 3H). ^13^C NMR (176 MHz, CDCl_3_) δ 156.8, 141.4, 138.3, 135.6, 132.4, 132.1, 131.9, 129.8, 129.5, 129, 128.2, 128.0, 126.8, 126.6, 126.1, 120.8, 83.7, 77, 66, 62.3, 55.9, 43, 29.1, 13.6.

#### General Procedure for the Synthesis of Tam-KDMi

4-(4-Ethynylbenzyl)−Tamoxifen **5** (100 mg, 1 equiv.) and the corresponding PMB-Maltol-Azide **3 (a-f)** (1 equiv.) were dissolved in 4 mL of dry THF and DMSO (in a 1:1 ratio) under an argon atmosphere. Argon was bubbled through the solution for 10 min before and after the addition of copper(I) iodide (4 mg, 0.1 equivalent) and diisopropyl ethylamine (0.043 mL, 1.2 equiv.). The reaction mixture was stirred at room temperature for 15 h. To quench the reaction, water (50 mL) was added, and the mixture was extracted with CH_2_Cl_2_ (2 × 20 mL). The combined organic layers were sequentially washed with a 4:1 mixture of NH_4_Cl and NH_4_OH (10 mL), then with 15 mL of water, and finally with brine solution (10 mL). The organic phases were combined, dried over Na_2_SO_4_, and filtered. The solvent was removed using a rotary evaporator, and the crude material was purified via silica gel column chromatography, eluting with MeOH: CH_2_Cl_2_ (1:12, v/v). This process yielded the intermediate PMB-protected compounds **6 (a-f)** as solid. Each intermediate compound was deprotected by stirring in a 10% solution of TFA in CH_2_Cl_2_ (2 mL) at room temperature for 1 h. The resulting crude material was further purified using silica gel column chromatography, eluting with MeOH: CH_2_Cl_2_ (2:10, v/v), which afforded the desired Tam-KDMi as red solid.

#### (Z)-1-(2-(4-(4-(((2-(4-(1,2-Diphenylbut-1-en-1 yl)phenoxy)ethyl)(methyl)amino)methyl)phenyl)-1H-1,2,3-triazol-1-yl)ethyl)-3-hydroxy-2-methylpyridin-4(1H)-one **(DW-54)**

Following the general procedure using 4-(4-ethynylbenzyl)−Tamoxifen **5** (100 mg), 1-(2-azidoethyl)-3-((4-methoxybenzyl) oxy)-2-methylpyridin-4(1H)-one (**3a**) (67mg), CuI (4 mg) and DIPEA (0.043 mL) gave **DW- 54** (71 mg, 50 % yield after two steps) as a brown solid. ^1^H NMR (700 MHz, CDCl_3_) δ 7.67 (s, 2H), 7.33 (t, *J* = 7.4 Hz, 4H), 7.29 – 7.18 (m, 5H), 7.13 (dd, *J* = 20.3, 7.3 Hz, 5H), 6.76 (d, *J* = 8.4 Hz, 2H), 6.54 (d, *J* = 8.5 Hz, 2H), 6.22 (s, 1H), 4.46 (s, 2H), 3.96 (t, *J* = 5.9 Hz, 2H), 3.58 (s, 2H), 3.25 (s, 2H), 2.75 (t, *J* = 5.8 Hz, 2H), 2.45 (q, *J* = 7.3 Hz, 2H), 2.28 (s, 6H), 0.91 (d, *J* = 7.3 Hz, 3H). ^13^C NMR (176 MHz, CDCl_3_) δ 170, 156.8, 148.3, 146.6, 143.9, 142.5, 141.4, 139.4, 138.3, 135.6, 131.9, 129.8, 129.7, 129.5, 128.6, 128.2, 127.9, 127.5, 126.6, 126.1, 125.8, 113.4, 66, 62.3, 55.8, 42.9, 29.8, 29.1, 13.7.12.1. HRMS (ESI) m/z Calcd. for C_42_H_43_N_5_O_3_ [M+H] ^+^: 666.3444, found 666.3447.

#### (Z)-1-(3-(4-(4-(((2-(4-(1,2-Diphenylbut-1-en-1-yl)phenoxy)ethyl)(methyl)amino)methyl)phenyl)-1H-1,2,3-triazol-1-yl)propyl)-3-hydroxy-2-methylpyridin-4(1H)-one (**DW-143**)

Following the general procedure using 4-(4-ethynylbenzyl)−Tamoxifen **5** (100 mg), 1-(3-azidopropyl)-3-((4-methoxybenzyl)oxy)-2 methylpyridin-4(1H)-one **(3b)** (70 mg), CuI (4 mg) and DIPEA (0.043 mL) gave **DW-143** (76 mg, 52 % yield after two steps) as a brown solid. ^1^H NMR (700 MHz, CDCl_3_) δ 7.75 (s, 2H), 7.38 (d, J = 7.9 Hz, 2H), 7.33 (t, J = 7.6 Hz, 3H), 7.23 (s, 4H), 7.16 (t, J = 7.5 Hz, 2H), 7.10 (d, J = 8.0 Hz, 3H), 6.76 (d, J = 6.7 Hz, 2H), 6.54 (d, J = 6.6 Hz, 2H), 6.37 (s, 1H), 4.43 (s, 2H), 3.99 (d, J = 28.8 Hz, 2H), 3.96 (d, J = 5.9 Hz, 2H), 3.61 (s, 2H), 2.77 (t, J = 6.0 Hz, 2H), 2.45 (q, J = 7.4 Hz, 2H), 2.32 (m, 5H), 2.30 (s, 3H), 0.92 (t, J = 7.4 Hz, 3H). ^13^C NMR (176 MHz, CDCl_3_) δ 169.7, 156.8, 148.3, 143.9, 141.4, 139.3, 138.2, 137.3, 135.6, 131.9, 129.8, 129.7, 129.5, 129, 128.2, 127.9, 126.6, 126.1, 125.7, 119.9, 113.4, 111.5, 66, 62.3, 55.8, 50.6, 46.6, 42.9, 31, 29.1, 13.7, 11.9. HRMS (ESI) m/z Calcd. For C_43_H_45_N_5_O_3_ [M+H] ^+^: 680.3601, found 680.3606.

#### (Z)-1-(4-(4-(4-(((2-(4-(1,2-Diphenylbut-1-en-1-yl)phenoxy)ethyl)(methyl)amino)methyl)phenyl)-1H-1,2,3-triazol-1-yl)butyl)-3-hydroxy-2-methylpyridin-4(1H)-one (**DW-136**)

Following the general procedure using 4-(4-ethynylbenzyl)−Tamoxifen **5** (100 mg), 1-(4-azidobutyl)-3-((4-methoxybenzyl)oxy)-2-methylpyridin-4(1H)-one **(3c)** (73 mg), CuI (4 mg) and DIPEA (0.043 mL) gave **DW-136** (84 mg, 57 % yield after two steps) as a brown solid. ^1^H NMR (700 MHz, CDCl_3_) δ 7.77 (s, 2H), 7.43 – 7.32 (m, 4H), 7.27 (dd, *J* = 18.0, 6.6 Hz, 4H), 7.22 – 7.07 (m, 6H), 6.79 (d, *J* = 8.6 Hz, 2H), 6.56 (d, *J* = 8.7 Hz, 2H), 6.36 (s, 1H), 4.44 (s, 2H), 3.99 (s, 2H), 3.98 (d, *J* = 5.9 Hz, 2H), 3.61 (s, 2H), 2.78 (t, *J* = 5.9 Hz, 2H), 2.47 (q, *J* = 7.4 Hz, 2H), 2.35 (s, 3H), 2.31 (s, 3H), 2.02 (m, 2H), 1.79 (m, 2H), 0.94 (t, *J* = 7.4 Hz, 3H). ^13^C NMR (176 MHz, CDCl_3_) δ 169.6, 156.7, 148, 146.5, 143.8, 142.4, 141.4, 139.1, 138.2, 136.8, 135.6, 131.9, 129.7, 129.6, 129.5, 129.1, 128.1, 127.9, 126.6, 126, 125.6, 119.8, 113.4, 111.4, 66, 62.3, 55.8, 42.9, 29.8, 29, 27.9, 27.1, 13.6, 11.9. HRMS (ESI) m/z Calcd. for C_44_H_47_N_5_O_3_ [M+H]^+^: 694.3757, found 694.3759.

#### (Z)-1-(5-(4-(4-(((2-(4-(1,2-Diphenylbut-1-en-1-yl)phenoxy)ethyl)(methyl)amino)methyl)phenyl)-1H-1,2,3-triazol-1-yl)pentyl)-3-hydroxy-2-methylpyridin-4(1H)-one **(DW-116)**

Following the general procedure using 4-(4-ethynylbenzyl)−Tamoxifen **5** (100 mg), 1-(5-azidopentyl)-3-((4-methoxybenzyl)oxy)-2-methylpyridin-4(1H)-one **(3d)** (76 mg), CuI (4 mg) and DIPEA (0.043 mL) gave **DW-116** (94 mg, 62 % yield after two steps) as a brown solid. ^1^H NMR (700 MHz, CDCl_3_) δ 7.76 (s, 2H), 7.34 (dd, *J* = 15.3, 7.9 Hz, 4H), 7.28 – 7.22 (m, 4H), 7.19 – 7.05 (m, 6H), 6.75 (s, 2H), 6.54 (d, *J* = 6.4 Hz, 2H), 6.35 (s, 1H), 4.41 (s, 2H), 3.97 (t, *J* = 5.9 Hz, 2H), 3.82 (s, 2H), 3.60 (s, 2H), 2.77 (t, *J* = 6.0 Hz, 2H), 2.45 (q, *J* = 7.4 Hz, 2H), 2.36 (s, 3H), 2.30 (s, 3H), 1.99 (m, 2H), 1.76 (m, 2H), 1.39 (m, 2H), 0.92 (t, *J* = 7.4 Hz, 3H). ^13^C NMR (176 MHz, CDCl_3_) δ 169.6, 156.8, 147.9, 146.4, 143.9, 142.5, 141.4, 139, 138.3, 136.8, 135.6, 131.9, 129.8, 129.6, 129.5, 129.3, 128.1, 127.9, 127.8, 126.6, 126.1, 125.6, 119.5, 113.4, 111.2, 66, 62.3, 55.8, 53.6, 49.9, 42.9, 41, 30.4, 29.9, 29.1, 23.4, 15.3, 11.9. HRMS (ESI) m/z Calcd. For C_45_H_49_N_5_O_3_ [M+H]^+^: 708.3914, found 708.3918.

#### (Z)-1-(6-(4-(4-(((2-(4-(1,2-Diphenylbut-1-en-1-yl)phenoxy)ethyl)(methyl)amino)methyl)phenyl)-1H-1,2,3-triazol-1-yl)hexyl)-3-hydroxy-2-methylpyridin-4(1H)-one **(DW-122)**

Following the general procedure using 4-(4-ethynylbenzyl)−Tamoxifen **5** (100 mg), 1-(6-azidohexyl)-3-((4-methoxybenzyl)oxy)-2-methylpyridin-4(1H)-one (**3e**) (79 mg), CuI (4 mg) and DIPEA (0.043 mL) gave **DW-122** (108 mg, 70 % yield after two steps) as a brown solid. ^1^H NMR (700 MHz, CDCl_3_) δ 7.77 (s, 2H), 7.37 (dd, J = 16.3, 9.2 Hz, 4H), 7.30 – 7.24 (m, 4H), 7.23 – 7.09 (m, 6H), 6.78 (d, J = 8.9 Hz, 2H), 6.56 (d, J = 6.6 Hz, 2H), 6.38 (s, 1H), 4.42 (s, 2H), 3.99 (t, J = 5.9 Hz, 2H), 3.85 (s, 2H), 3.62 (s, 2H), 2.79 (t, J = 6.0 Hz, 2H), 2.47 (q, J = 7.4 Hz, 2H), 2.38 (s, 3H), 2.31 (s, 3H), 1.98 (m, 2H), 1.73 (m, 2H), 1.40 (m, 4H), 0.94 (t, J = 7.4 Hz, 3H). ^13^C NMR (176 MHz, CDCl_3_) δ 169.5, 156.6, 147.8, 146.4, 143.8, 142.4, 141.4, 138.2, 136.9, 135.78, 132.0, 129.9, 129.8, 129.6, 128.2, 128.0, 126.6, 126.1, 125.7, 119.7, 113.4, 111.2, 65.7, 62.1, 55.7, 53.8, 50.1, 42.6, 30.8, 30.1, 29.7, 29.1, 26.1, 25.8, 13.6, 11.9. HRMS (ESI) m/z Calcd. for C_46_H_51_N_5_O_3_ [M+H]^+^: 722.4070, found 722.4073.

#### (Z)-1-(7-(4-(4-(((2-(4-(1,2-diphenylbut-1-en-1-yl)phenoxy)ethyl)(methyl)amino)methyl)phenyl)-1H-1,2,3-triazol-1-yl)heptyl)-3-hydroxy-2-methylpyridin-4(1H)-one (**DW-55**)

Following the general procedure using 4-(4-ethynylbenzyl)−Tamoxifen **5** (100 mg), 1-(7-azidoheptyl)-3-((4-methoxybenzyl)oxy)-2-methylpyridin-4(1H)-one (**3f**) (82 mg), CuI (4 mg) and DIPEA (0.043 mL) gave **DW-55** (89 mg, 57 % yield after two steps) as a brown solid. ^1^H NMR (700 MHz, CDCl_3_) δ 7.77 (s, 2H), 7.41 – 7.30 (m, 4H), 7.28 – 7.22 (m, 4H), 7.18 – 7.07 (m, 6H), 6.77 (d, *J* = 8.7 Hz, 2H), 6.54 (d, *J* = 8.7 Hz, 2H), 6.35 (s, 1H), 4.38 (s, 2H), 3.99 (s, 2H), 3.81 (s, 2H), 3.63 (s, 2H), 2.80 (s, 2H), 2.45 (q, *J* = 7.5 Hz, 2H), 2.35 (s, 3H), 2.31 (s, 3H), 1.93 (m, 2H), 1.68 (m, 2H), 1.34 (m, *J* = 13.9 Hz, 6H), 0.92 (t, *J* = 7.4 Hz, 3H). ^13^C NMR (176 MHz, CDCl_3_) δ 169.4, 156.6, 147.6, 146.3, 143.8, 141.3, 138.2, 136.7, 135.6, 131.9, 129.7, 129.4, 128.1, 128, 127.9, 126.5, 126, 125.6, 119.5, 113.4, 111.1, 65.8, 62.2, 55.7, 53.8, 50.2, 42.7, 30.7, 30.1, 29, 28.5, 26.3, 13.6, 11.9. HRMS (ESI) m/z Calcd. for C_47_H_53_N_5_O_3_ [M+H]^+^: 736.4227, found 736.4229.

#### General Procedure for the Synthesis of EED-KDMi

17α-Ethynylestradiol (1 equiv.) and the corresponding PMB-Maltol-Azide **3 (c-g)** (1 equiv.) were dissolved in 4 mL of dry THF and DMSO (in a 1:1 ratio) under an argon atmosphere. Argon was bubbled through the solution for 10 min before and after the addition of copper(I) iodide (0.1 equivalent) and DIPEA (1.2 equiv.). The reaction mixture was stirred at room temperature for 15 h. Completion of the reaction was monitored by TLC. To quench the reaction, water (50 mL) was added, and the mixture was extracted with CH_2_Cl_2_ (2 × 20 mL). The combined organic layers were sequentially washed with a 4:1 mixture of NH_4_Cl and NH_4_OH (10 mL), then with 15 mL of water, and finally with brine solution (10 mL). The organic phases were combined, dried over Na_2_SO_4_, and filtered. The solvent was removed using a rotary evaporator, and the crude material was purified via silica gel column chromatography, eluting with MeOH: CH_2_Cl_2_ (1:12, v/v). This process yielded the corresponding intermediate PMB-protected compound **7 (c-g)** as a solid. Each intermediate compound was deprotected by stirring in a 10% solution of TFA in CH_2_Cl_2_ (2 mL) at room temperature for 1 h. The resulting crude material was further purified using silica gel column chromatography, eluting with MeOH: CH_2_Cl_2_ (2:10, v/v), which afforded the desired EED-KDMi as brown or red solid.

#### 1-(4-(4-((8R,9S,13S,14S,17S)-3,17-dihydroxy-13-methyl-7,8,9,11,12,13,14,15,16,17-decahydro-6H-cyclopenta[a]phenanthren-17-yl)-1H-1,2,3-triazol-1-yl)butyl)-3-hydroxy-2-methylpyridin-4(1H)-one **(DW-554)**

Following the general procedure using 17α-ethynylestradiol (100 mg), 1-(4-azidobutyl)-3-((4-methoxybenzyl) oxy)-2-methylpyridin-4(1H)-one **(3c)** (116 mg), CuI (7 mg) and DIPEA (0.068 mL) gave **DW-554** (90 mg, 51 % yield after two steps) as a brown solid. ^1^H NMR (700 MHz, CD_3_OD+CDCl_3_) δ 7.69 (s, 1H), 7.66 (s, 1H), 7.48 (d, *J* = 7.2 Hz, 1H), 7.00 (d, *J* = 8.6 Hz, 1H), 6.53 (d, *J* = 8.5 Hz, 1H), 6.48 (s, 1H), 6.36 (d, *J* = 7.2 Hz, 1H), 4.43 (t, *J* = 6.9 Hz, 2H), 4.01 (t, *J* = 7.7 Hz, 2H), 3.35 (s, 1H), 2.87 – 2.63 (m, 2H), 2.45 (dd, *J* = 20.7, 10.6 Hz, 1H), 2.38 (s, 3H), 2.14 – 2.03 (m, 2H), 1.98 (m, 2H), 1.94 – 1.85 (m, 2H), 1.73 (s, 1H), 1.66 – 1.49. (m, 4H), 1.49 –1.34 (m, 2H), 1.30 (d, *J* = 18.2 Hz, 4H), 1.01 (s, 3H), 0.61 (d, *J* = 12.9 Hz, 1H). ^13^C NMR (176 MHz, CD_3_OD+CDCl_3_) δ 170, 155.2, 138.5, 138.1, 132.1, 131.6, 126.8, 123.3, 115.8, 113.3, 112.4, 82.8, 54, 50, 47.9, 44.3, 40.4, 38, 33.8, 30.3, 28.3, 28.2, 27.6, 27, 24.2, 14.6, 11.9. HRMS (ESI) m/z Calcd. for C_30_H_38_N_4_O_4_ [M+H] ^+^: 519.2966, found 519.2969.

#### 1-(5-(4-((8R,9S,13S,14S,17S)-3,17-dihydroxy-13-methyl-7,8,9,11,12,13,14,15,16,17-decahydro-6H-cyclopenta[a]phenanthren-17-yl)-1H-1,2,3-triazol-1-yl)pentyl)-3-hydroxy-2-methylpyridin-4(1H)-one **(DW-613)**

Following the general procedure using 17α-ethynylestradiol (100 mg), 1-(5-azidopentyl)-3-((4-methoxybenzyl) oxy)-2-methylpyridin-4(1H)-one (**3d**) (120 mg), CuI (7 mg) and DIPEA (0.068 mL) gave **DW-613** (96 mg, 53 % yield after two steps) as a brown solid. ^1^H NMR (700 MHz, CD_3_OD+CDCl_3_) δ 7.52 (s, 1H), 7.29 (d, *J* = 7.2 Hz, 1H), 6.97 (d, *J* = 8.5 Hz, 1H), 6.53 (d, *J* = 8.6 Hz, 1H), 6.49 (s, 1H), 6.35 (d, *J* = 7.2 Hz, 1H), 4.34 (d, *J* = 18.5 Hz, 2H), 3.87 (t, *J* = 7.6 Hz, 2H), 2.82 – 2.68 (m, 2H), 2.40 (m *J* = 9.4, 4.4 Hz, 1H), 2.35 (s, 3H), 2.12 – 2.02 (m, 2H), 1.93 (d, *J* = 7.4 Hz, 2H), 1.90 – 1.82 (m, 2H), 1.73 (d, *J* = 8.3 Hz, 2H), 1.55 (d, *J* = 11.2 Hz, 2H), 1.51 (s, 1H), 1.45 – 1.23 (m, 6H), 0.99 (s, 3H), 0.60 (t, *J* = 13.0 Hz, 1H). 13C NMR (176 MHz, CD_3_OD+CDCl_3_) δ 169.3, 154.5, 154.3, 146.2, 138.1, 137.2, 131.6, 130.3, 126.3, 122.2, 115.4, 112.8, 111.8, 82.4, 54.1, 53.6, 49.9, 47.4, 43.6, 39.6, 37.5, 33.1, 30.2, 27.5, 26.4, 23.7, 23.4, 14.1, 11.8. HRMS (ESI) m/z Calcd. for C_31_H_40_N_4_O_4_ [M+H] ^+^: 533.3122, found 533.3121.

#### 1-(6-(4-((8R,9S,13S,17S)-3,17-dihydroxy-13-methyl-7,8,9,11,12,13,14,15,16,17-decahydro-6H-cyclopenta[a]phenanthren-17-yl)-1H-1,2,3-triazol-1-yl)hexyl)-3-hydroxy-2-methylpyridin-4(1H)-one (**DW-088)**

Following the general procedure using 17α-ethynylestradiol (1 g), 1-(6-azidohexyl)-3-((4-methoxybenzyl)oxy)-2-methylpyridin-4(1H)-one (**3e**) (1.25 g), CuI (64 mg) and DIPEA (0.68 mL), 20 mL of dry THF and DMSO gave **DW-088** (1.126 g, 61 % yield after two steps) as a brown solid. ^1^H NMR (700 MHz, CD_3_OD+CDCl_3_) δ 7.55 (s, 1H), 7.32 (d, J = 7.2 Hz, 1H), 6.95 (d, J = 8.6 Hz, 1H), 6.56 – 6.44 (m, 2H), 6.35 (d, J = 7.1 Hz, 1H), 5.31 (s, 1H), 4.34 (t, J = 7.0 Hz, 2H), 3.89 (t, J = 7.6 Hz, 2H), 2.83 – 2.65 (m, 2H), 2.62 (s, 1H), 2.46 – 2.38 (m, 1H), 2.36 (s, 3H), 2.07 (q, J = 11.0 Hz, 2H), 1.96 – 1.83 (m, 4H), 1.74 – 1.60 (m, 2H), 1.60 – 1.48 (m, 3H), 1.43 (d, J = 11.1 Hz, 1H), 1.40 – 1.25 (m, 6H), 0.99 (s, 3H), 0.62 (t, J = 12.3 Hz, 1H). ^13^C NMR (176 MHz, CD_3_OD+CDCl_3_) δ 169.4, 154.7, 154.4, 146.4, 138.2, 137.4, 131.7, 130.7, 126.4, 122.4, 115.5, 112.9, 111.9, 82.4, 66.2, 54.4, 53.8, 50.3, 47.5, 43.8, 40.3, 39.8, 37.6, 33.3, 30.9, 30.2, 29.9, 27.7, 26.6, 26.3, 26, 23.8, 14.4, 11.9. HRMS (ESI) m/z Calcd. for C_32_H_42_N_4_O_4_ [M+H]^+^: 547.3279, found 547.3275.

#### 1-(7-(4-((8R,9S,13S,17S)-3,17-dihydroxy-13-methyl-7,8,9,11,12,13,14,15,16,17-decahydro-6H-cyclopenta[a]phenanthren-17-yl)-1H-1,2,3-triazol-1-yl)heptyl)-3-hydroxy-2-methylpyridin-4(1H)-one **(DW-614)**

Following the general procedure using 17α-ethynylestradiol (100 mg), 1-(7-azidoheptyl)-3-((4-methoxybenzyl)oxy)-2-methylpyridin-4(1H)-one (**3f**) (130 mg), CuI (7 mg) and DIPEA (0.068 mL) gave **DW-614** (103 mg, 54 % yield after two steps) as a brown solid. ^1^H NMR (700 MHz, CD_3_OD+CDCl_3_) δ 7.56 (s, 1H), 7.29 (d, *J* = 7.2 Hz, 1H), 6.99 (d, *J* = 8.5 Hz, 1H), 6.55 (d, *J* = 8.7 Hz, 1H), 6.51 (s, 1H), 6.37 (d, *J* = 7.2 Hz, 1H), 4.34 (t, *J* = 7.1 Hz, 2H), 3.87 (t, *J* = 7.7 Hz, 2H), 2.83 – 2.68 (m, 2H), 2.46 – 2.39 (m, 1H), 2.37 (s, 3H), 2.09 (q, *J* = 10.3 Hz, 2H), 1.94 (d, *J* = 5.4 Hz, 1H), 1.93 – 1.84 (m, 4H), 1.72 – 1.63 (m, 2H), 1.63 – 1.51 (m, 3H), 1.45 (d, *J* = 21.9 Hz, 1H), 1.36 (m, *J* = 20.8 Hz, 3H), 1.29 (m, 4H), 1.01 (s, 3H), 0.64 (t, J = 13.0 Hz, 1H). ^13^C NMR (176 MHz, CD_3_OD+CDCl_3_) δ 172.2, 154.7, 154.2, 146.3, 138.2, 137.3, 131.6, 130.7, 126.3, 122.3, 115.5, 112.9, 111.8, 82.4, 60.9, 54.5, 50.4, 47.5, 43.8, 39.7, 37.4, 33.3, 30.9, 30.3, 29.9, 28.7, 27.7, 26.5, 26.5, 23.7, 21, 14.3, 11.9. HRMS (ESI) m/z Calcd. for C_33_H_44_N_4_O_4_ [M+H+]: 561.3435, found 561.3439.

#### 1-(8-(4-((8R,9S,13S,17S)-3,17-dihydroxy-13-methyl-7,8,9,11,12,13,14,15,16,17-decahydro-6H-cyclopenta[a]phenanthren-17-yl)-1H-1,2,3-triazol-1-yl)octyl)-3-hydroxy-2-methylpyridin-4(1H)-one **(DW-615)**

Following the general procedure using 17α-ethynylestradiol (100 mg), 1-(8-azidooctyl)-3-((4-methoxybenzyl) oxy)-2-methylpyridin-4(1H)-one (**3 g**) (67mg), CuI (7 mg), and DIPEA (0.068 mL) gave **DW-615** (94 mg, 48 % yield after two steps) as a brown solid. ^1^H NMR (700 MHz, CD_3_OD+CDCl_3_) δ 7.56 (s, 1H), 7.28 (d, *J* = 7.2 Hz, 1H), 7.01 (d, *J* = 8.2 Hz, 1H), 6.55 (d, *J* = 20.9 Hz, 2H), 6.40 (d, *J* = 7.0 Hz, 1H), 4.37 (t, *J* = 6.9 Hz, 2H), 3.86 (t, *J* = 7.6 Hz, 2H), 2.91 – 2.69 (m, 2H), 2.66 (s, 1H), 2.46 (d, *J* = 14.0 Hz, 1H), 2.39 (s, 2H), 2.20 – 1.97 (m, 3H), 1.97 – 1.78 (m, 5H), 1.78 – 1.41 (m, 5H), 1.31 (m, 8H), 1.04 (s, 3H), 0.72 – 0.56 (m, 1H). ^13^C NMR (176 MHz, CD_3_OD+CDCl_3_) δ 169.3, 154.6, 154, 138.2, 137.1, 131.6, 130.5, 129.4, 128.5, 126.3, 122.2, 115.4, 112.8, 111.8, 82.1, 54.4, 43.6, 39.7, 37.5, 33.2, 31, 30.3, 29.8, 29.2, 28.9, 27.6, 26.5, 26.4, 26.4, 23.7, 14.3, 11.9. HRMS (ESI) m/z Calcd. for C_34_H_46_N_4_O_4_ [M+H+]: 575.3591, found 575.3596.

#### · 3- Hydroxy-1-(6-(4-((8R,9S,13S,17S)-17-hydroxy-3-methoxy-13-methyl 7,8,9,11,12,13,14,15,16,17-decahydro-6H-cyclopenta[a]phenanthren-17-yl)-1H-1,2,3-triazol-1-yl)hexyl)-2-methylpyridin-4(1H)-one (**DW-95**)

Mestranol (1 g, 1 equiv.) and 1-(6-azidohexyl)-3-((4-methoxybenzyl)oxy)-2-methylpyridin-4(1H)-one (**3e**) (1.193 g, 1 equiv.) were dissolved in 20 mL of dry THF and DMSO (in a 1:1 ratio) under an argon atmosphere. Argon was bubbled through the solution 10 min before and after the addition of CuI (61 mg,0.1 equivalent) and DIPEA (0.657 mL, 1.2 equiv.). The reaction mixture was stirred at room temperature for 15 h. Completion of the reaction was monitored by TLC. To quench the reaction, water (50 mL) was added, and the mixture was extracted with CH_2_Cl_2_ (3 × 50 mL). The combined organic layers were sequentially washed with a 4:1 mixture of NH_4_Cl and NH_4_OH (20 mL), then with 15 mL of water, and finally with brine solution (10 mL). The organic phases were combined, dried over Na_2_SO_4_, and filtered. The solvent was removed using a rotary evaporator, and the crude material was purified via silica gel column chromatography, eluting with MeOH: CH_2_Cl_2_ (1:12, v/v). This process yielded the intermediate compound **8** as a solid. The intermediate compound was deprotected by stirring in a 10% solution of TFA in CH_2_Cl_2_ (10 mL) at room temperature for 1 h. The resulting crude material was further purified using silica gel column chromatography, eluting with MeOH: CH_2_Cl_2_ (2:10, v/v), which afforded **DW- 95** in 1.287 g (71%) as a red solid. ^1^H NMR (700 MHz, CDCl_3_) δ 7.43 (s, 1H), 7.18 (s, 1H), 7.07 (d, *J* = 8.7 Hz, 1H), 6.65 (d, *J* = 8.8 Hz, 1H), 6.59 (s, 1H), 6.34 (s, 1H), 33 (s, 2H), 3.84 (s, 2H), 3.74 (s, 3H), 2.88 – 2.77 (m, 2H), 2.60 (s, 3H), 2.38 (s, 1H), 2.34 (s, 2H), 2.12 (d, *J* = 13.1 Hz, 2H), 1.92 (m, 4H), 1.69 (s, 2H), 1.58 (d, 2H), 1.44 (d, 2H), 1.35 (m, 4H), 1.02 (s, 3H), 0.64 (s, 1H). ^13^C NMR (176 MHz, CDCl_3_) δ 169.4, 157.4, 153.9, 146.3, 138, 136.8, 132.5, 128.3, 126.2, 121.3, 113.8, 111.5, 82.3, 55.2, 53.8, 49.9, 48.5, 47.3, 43.5, 40.9, 39.5, 37.9, 33, 30.7, 30, 29.9, 27.4, 26.3, 26.1, 25.8, 23.5, 14.3, 11.9. HRMS (ESI) m/z Calcd. For C_33_H_44_N_4_O_4_ [M+H+]: 561.3435, found 561.3437.

## Supporting information

KDMi_Tamox_Steroid_Suppl Info_BioRxiv

## Author Contributions

¶These authors contributed equally to the manuscript.

## Author Contributions

D.T.W., J.O.O., and B.W. contributed equally to the manuscript. Experimental Design: D.T.W., J.O.O., B.W., R.K., A.J., S.B., J.K., F.A.E., T.J.N., N.A.H., M.T., and A.K.O. Data Generation: D.T.W., J.O.O., B.W., R.K., A.J., S.B., F.A.E., T.J.N., V.K., and V.V. Data Analysis: D.T.W., J.O.O., B.W., R.K., A.J., S.B., F.A.E., and T.J.N. Supervision: N.A.H., M.T. and A.K.O. Manuscript Writing: D.T.W., J.O.O., B.W., R.K., A.J., S.B., F.A.E., T.J.N., N.A.H., M.T., and A.K.O. Manuscript Review: All Funding: N.A.H., M.T. and A.K.O.

## Notes

The authors declare no competing financial interest.

## Acknowledgements

This project was financially supported by NIH grants R01CA252720 (A.K.O), R01CA275840 (M. T), and T32CA244125 (UNC Integrated Translational Oncology Program to UNC/TJN). The UNC Flow Cytometry Core Facility (RRID: SCR_019170) is supported in part by a Cancer Center Core Support Grant (P30CA016086) to the UNC Lineberger Comprehensive Cancer Center.

## ABBREVIATIONS

AURKA: Aurora kinase A
BCa: Breast cancer
CIP: chemical induced proximity
CiA: chromatin *in vivo* assay
CDKN1A/p21: cyclin-dependent kinase inhibitor 1
DFP: Deferiprone
ERα: estrogen receptor alpha
EED: 17α-ethinylestradiol
FDR: false discovery rate
Ful: Fulvestrant
GOPB: Gene ontology biological processes
GSEA: gene set enrichment analysis
HDACs: histone deacetylases
KDMs, HP1: heterochromatin protein 1; histone lysine demethylases
HMTs: histone methyltransferases
HER2: human epidermal growth factor receptor 2
HIF-1α: hypoxia inducible factor-1α
IVTI: *in vitro* therapeutic indexes
mES: mouse embryonic stem
NES: normalized enrichment score
PR: progesterone receptor
RNA-seq: RNA-sequencing
SERDs: selective estrogen-receptor modulators
SAR: structure activity relationship
Tam: Tamoxifen
TNBC: triple-negative breast cancer
TGI: tumor growth inhibition.

## Table of Contents Graphic

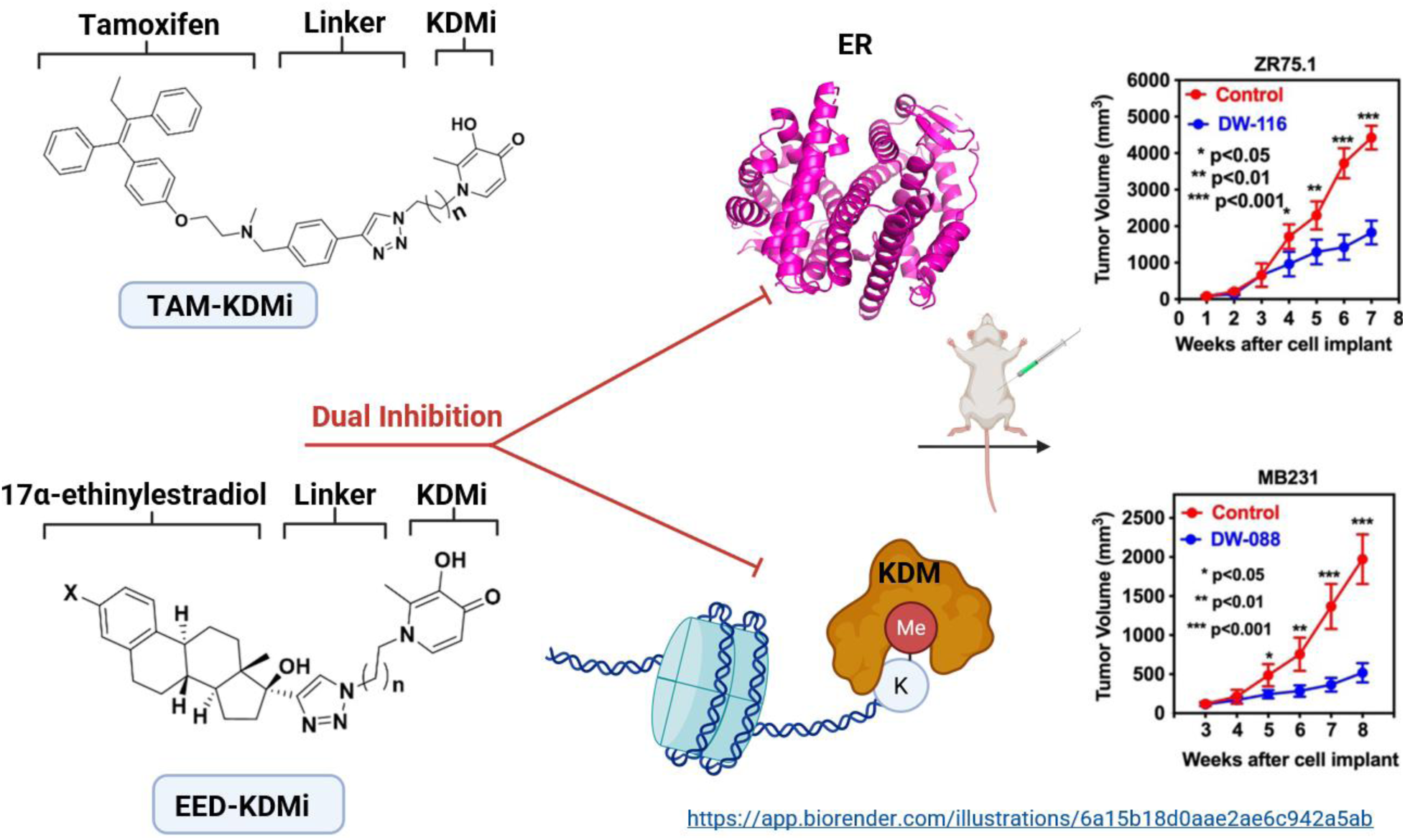

