## Supplementary material for "Dual-acting Estrogen Receptor Modulator-Histone Lysine Demethylase Inhibitors": KDMi_Tamox_Steroid_Suppl Info_BioRxiv

| Sr. No. | Table of Contents | Page No. |
| --- | --- | --- |
| **1** | ERα-binding affinity of Tam-KDMi and EED-KDMi | **S4** |
| **2** | Antiproliferative activity of Tam-KDMi and EED-KDMi | **S5** |
| **3** | Representative concentration-response antiproliferative curves of cells treated with Tam-KDMi conjugates (a.–h.). | **S6** |
| **4** | Representative concentration-response antiproliferative curves of cells treated with EED-KDMi conjugates (a.–h.). | **S7** |
| **5** | Effects of representative Tam-KDMi (**DW-122**), EED-KDMi (**DW-088** and **DW-095**), and tamoxifen on cell cycle progression in MCF-7 cell line | **S8** |
| **6** | Effects of representative Tam-KDMi (**DW-122**), EED-KDMi (**DW-088** and **DW-095**), and tamoxifen on cell cycle progression in MDA-MB-231 cell line | **S9** |
| **7** | Representative full gels (a.–c.) of MCF-7 cells treated with compounds **DW-116** and **DW-260 (DW-122)** | **S10** |
| **8** | Western blot densitometric quantifications (a.–f.) of MCF-7 breast cancer cells treated with compounds **DW-116** and **DW-260 (DW-122)** | **S11** |
| **9** | Representative full gels (a.–c.) of MCF-7 cells treated with deferiprone (DFP) and compound **DW-088** | **S12** |
| **10** | Western blot densitometric quantifications (a.–f.) of MCF-7 breast cancer cells treated with deferiprone (DFP) and compound **DW-088** | **S13** |
| **11** | Representative full gels (a–c) of MCF-7 cells treated with ethinyl estradiol (EED) and compound **DW-095** | **S14** |
| **12** | Western blot densitometric quantifications (a.–f.) of MCF-7 breast cancer cells treated with ethinyl estradiol (EED) and compound **DW-095** | **S15** |
| **13** | Representative full gels (a.–c.) of MDA-MB-231 cells treated with compounds **DW-116** and **DW-260 (DW-122)** | **S16** |

| | | |
|---|---|---|
| **14** | Western blot densitometric quantifications (a.–f.) of MDA-MB-231 breast cancer cells treated with compounds **DW-116** and **DW-260 (DW-122)** | **S17** |
| **15** | Representative full gels (a.–c.) of MDA-MB-231 cells treated with deferiprone (DFP) and compound **DW-088** | **S18** |
| **16** | Western blot densitometric quantifications (a.–f.) of MDA-MB-231 breast cancer cells treated with deferiprone (DFP) and compound **DW-088** | **S19** |
| **17** | Representative full gels (a–c) of MDA-MB-231 cells treated with ethinyl estradiol (EED) and compound **DW-095** | **S20** |
| **18** | Western blot densitometric quantifications (a.–f.) of MDA-MB-231 breast cancer cells treated with ethinyl estradiol (EED) and compound **DW-095** | **S21** |
| **19** | Heatmap of Gene Ontology Biological Process gene set enrichment analysis of MCF-7 and MDA-MB-231 breast cancer cells treated with **DW-088**, **DW-095**, and **DW-260**. | **S22** |
| **20** | $^{1}$H and $^{13}$C NMR Spectra | **S23-S43** |
| **21** | HPLC-UV Tracers | **S44-S47** |

### Supplementary Information

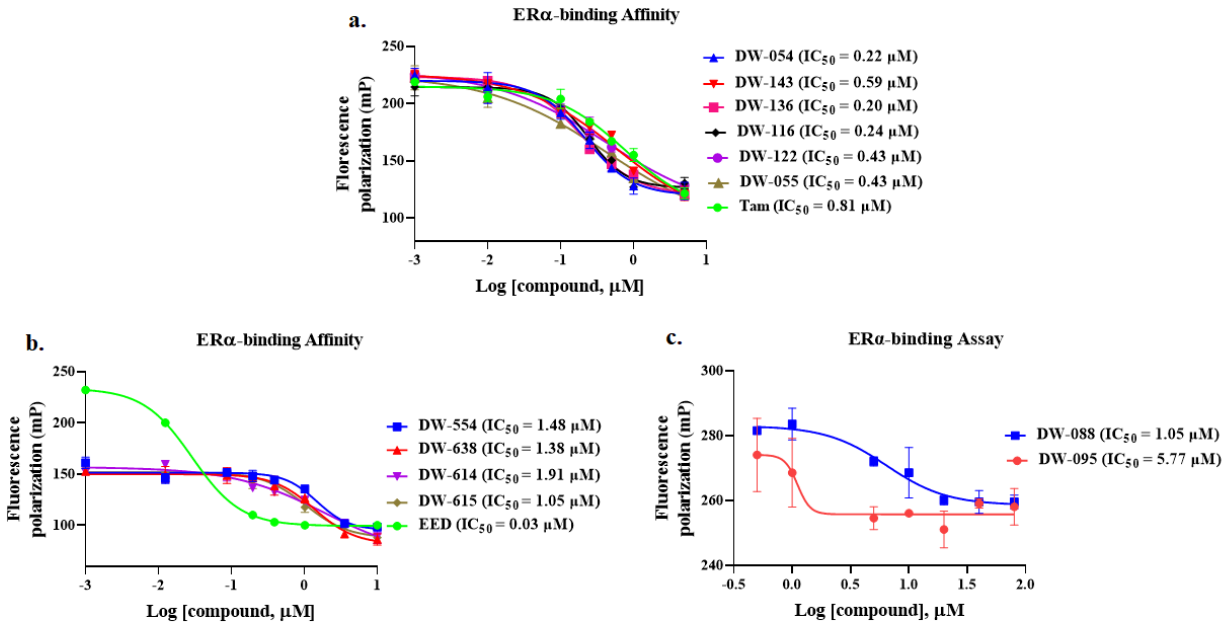


**Figure S1:** ERα-binding affinity of Tam-KDMi and EED-KDMi. Concentration-response curves and the median inhibitory concentration ($IC_{50}$) values of ERα binding affinity of tamoxifen (Tam), 17α-ethinylestradiol (EED) along with both Tam-KDMi and EED-KDMi (**a.–c.**) based on PolarScreen ER competitor assay.

**Table S1.** Antiproliferative activity of Tam-KDMi and EED-KDMi ($IC_{50}$, μM)

| Compound | DU-145 | LNCaP | HepG2 | SK-HEP-1 |
|---|---|---|---|---|
| **DW-054** | 1.27 ± 0.3 | 0.93 ± 0.4 | 1.28 ± 0.6 | 1.80 ± 1.0 |
| **DW-143** | 1.83 ± 1.2 | 1.12 ± 0.8 | 2.39 ± 1.2 | 2.72 ± 0.2 |
| **DW-136** | 1.28 ± 0.6 | 1.03 ± 0.7 | 2.27 ± 1.0 | 1.84 ± 0.7 |
| **DW-116** | 1.00 ± 0.1 | 1.12 ± 0.0 | 1.61 ± 0.2 | 1.57 ± 0.4 |
| **DW-122/DW-260** | 1.27 ± 0.5 | 0.98 ± 0.6 | 1.16 ± 0.3 | 1.90 ± 1.3 |
| **DW-055** | 1.51 ± 0.4 | 1.10 ± 0.0 | 0.96 ± 0.3 | 0.96 ± 0.1 |
| **DW-554** | 6.00 ± 2.3 | 10.8 ± 2.3 | 13.61 ± 4.5 | 38.56 ± 3.1 |
| **DW-613** | 58.31 ± 7.8 | >100 | 38.34 ± 11.2 | >100 |
| **DW-088** | 3.58 ± 1.1 | 7.50 ± 1.0 | 3.36 ± 0.0 | 1.82 ± 0.6 |
| **DW-614** | 5.96 ± 1.0 | 7.80 ± 3.3 | 11.82 ± 0.2 | >50 |
| **DW-615** | 4.60 ± 0.1 | 34.60 ± 0.9 | 8.31 ± 0.2 | 42.07 ± 4.8 |
| **DW-095** | 2.20 ± 0.7 | 0.90 | 1.40 ± 0.3 | 6.90 |
| **DFP** | >100 | >100 | >100 | >100 |
| **EED** | 104.6 | 50.49 | 55.06 | 134.9 |
| **Tam** | 22.03 ± 3.7 | 18.01 ± 4.1 | 11.58 ± 0.9 | 15.64 ± 3.7 |

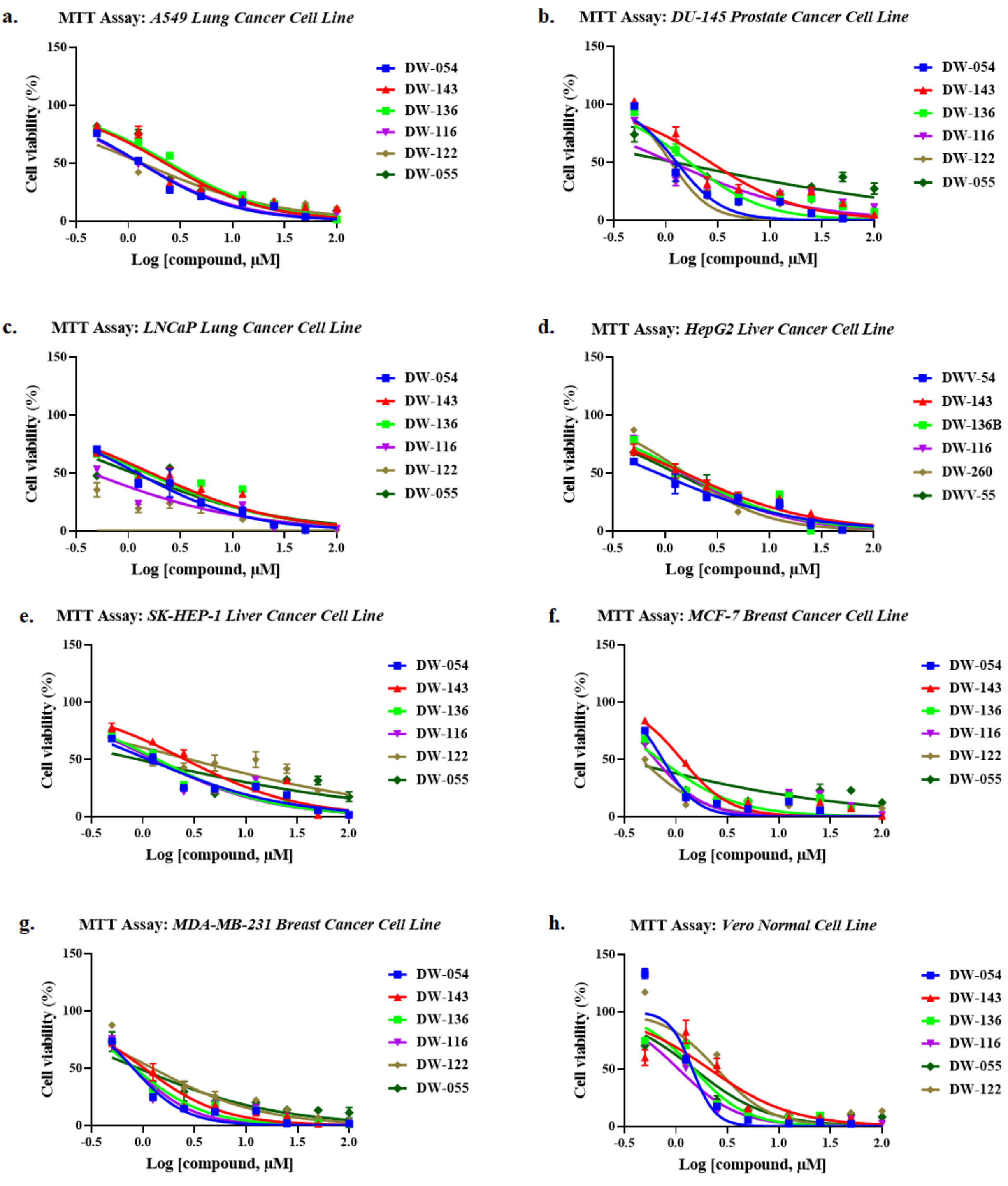


**Figure S2:** Representative concentration-response antiproliferative curves of cells treated with Tam-KDMi conjugates **(a.–h.).** The cells were treated for 72 h to determine the respective $IC_{50}$ values of the compounds using MTT cell viability assay.

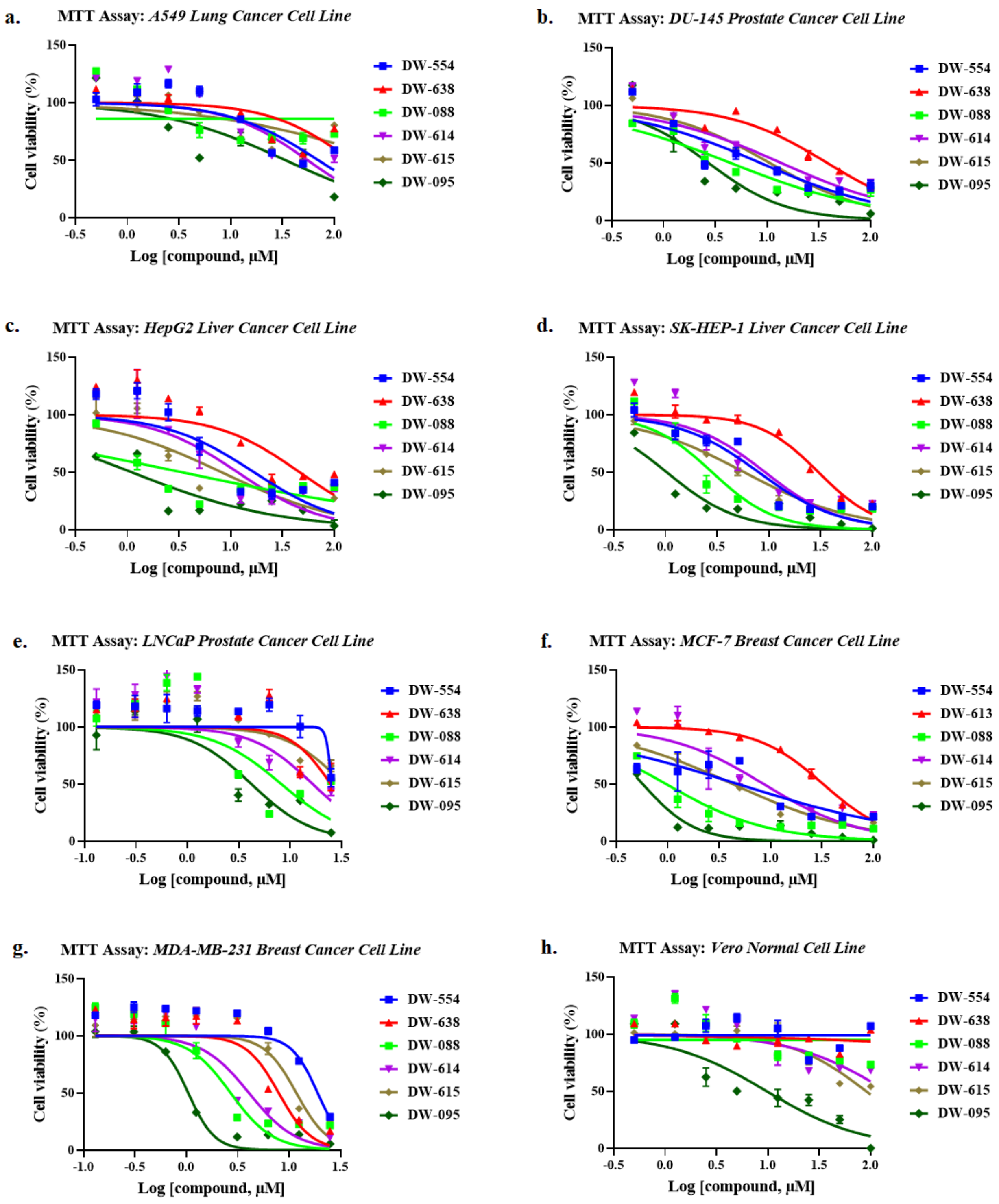


**Figure S3:** Representative concentration-response antiproliferative curves of cells treated with EED-KDMi conjugates **(a.–h.).** The cells were treated for 72 h to determine the respective $IC_{50}$ values of the compounds using MTT cell viability assay.

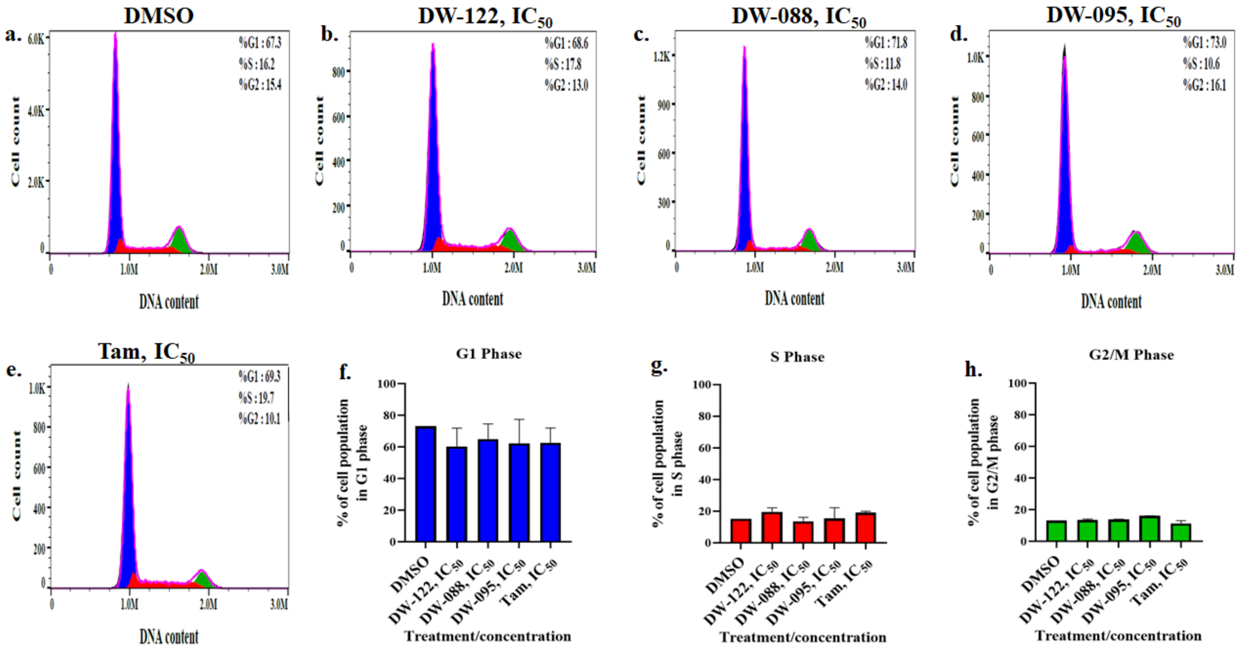


**Figure S4:** Effects of representative Tam-KDMi (**DW-122**), EED-KDMi (**DW-088** and **DW-095**), and tamoxifen on cell cycle progression in MCF-7 cell line treated for 24 h at $IC_{50}$ concentrations of the respective compounds. Histograms of the negative control (DMSO, **a**) and cells treated with the indicated compounds display the distribution of cell populations in G0/G1, S, and G2/M phases of the cell cycle (**b**-**e**). The effects of treatment on the phases of the cell cycle are further presented as bar charts (**f**-**h**) based on at least two independent experiments.

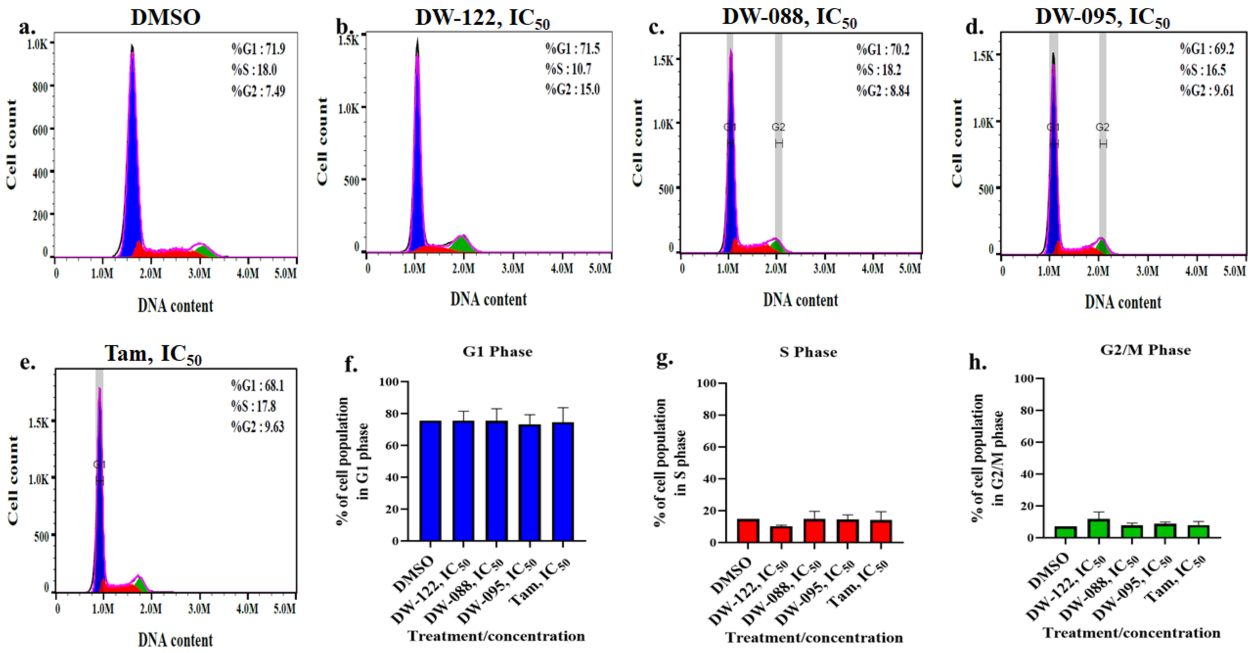


**Figure S5:** Effects of representative Tam-KDMi (**DW-122**), EED-KDMi (**DW-088** and **DW-095**), and tamoxifen on cell cycle progression in MDA-MB-231 cell line treated for 24 h at $IC_{50}$ concentrations of the respective compounds. Histograms of the negative control (DMSO, **a**) and cells treated with the indicated compounds display the distribution of cell populations in G0/G1, S, and G2/M phases of the cell cycle (**b**-**e**). The effects of treatment on the phases of the cell cycle are further presented as bar charts (**f**-**h**) based on at least two independent experiments.

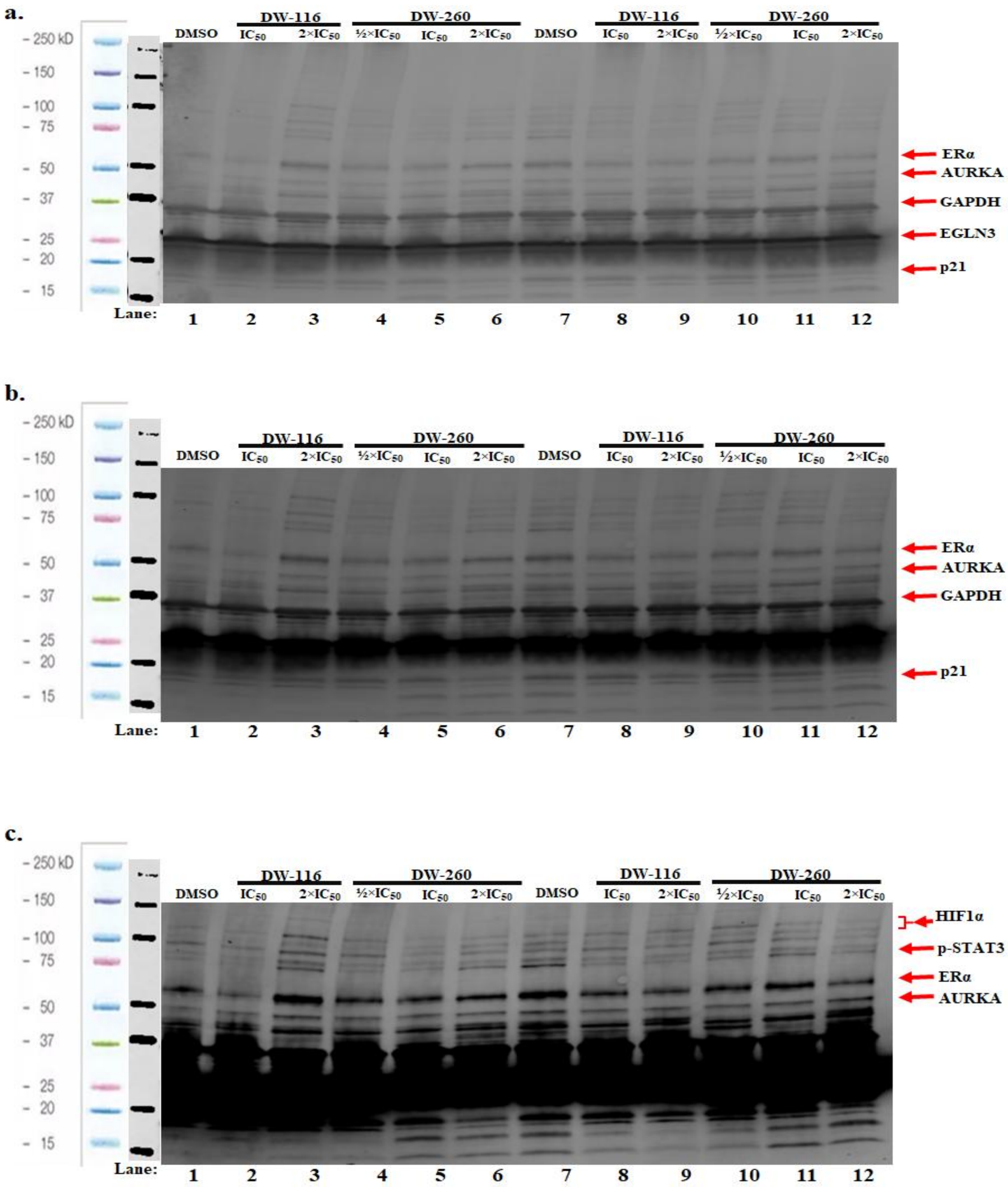


**Figure S6:** Representative full gels (**a**–**c**) of MCF-7 cells treated with compounds **DW-116** and **DW-260 (DW-122)** for 24 h, as indicated above.

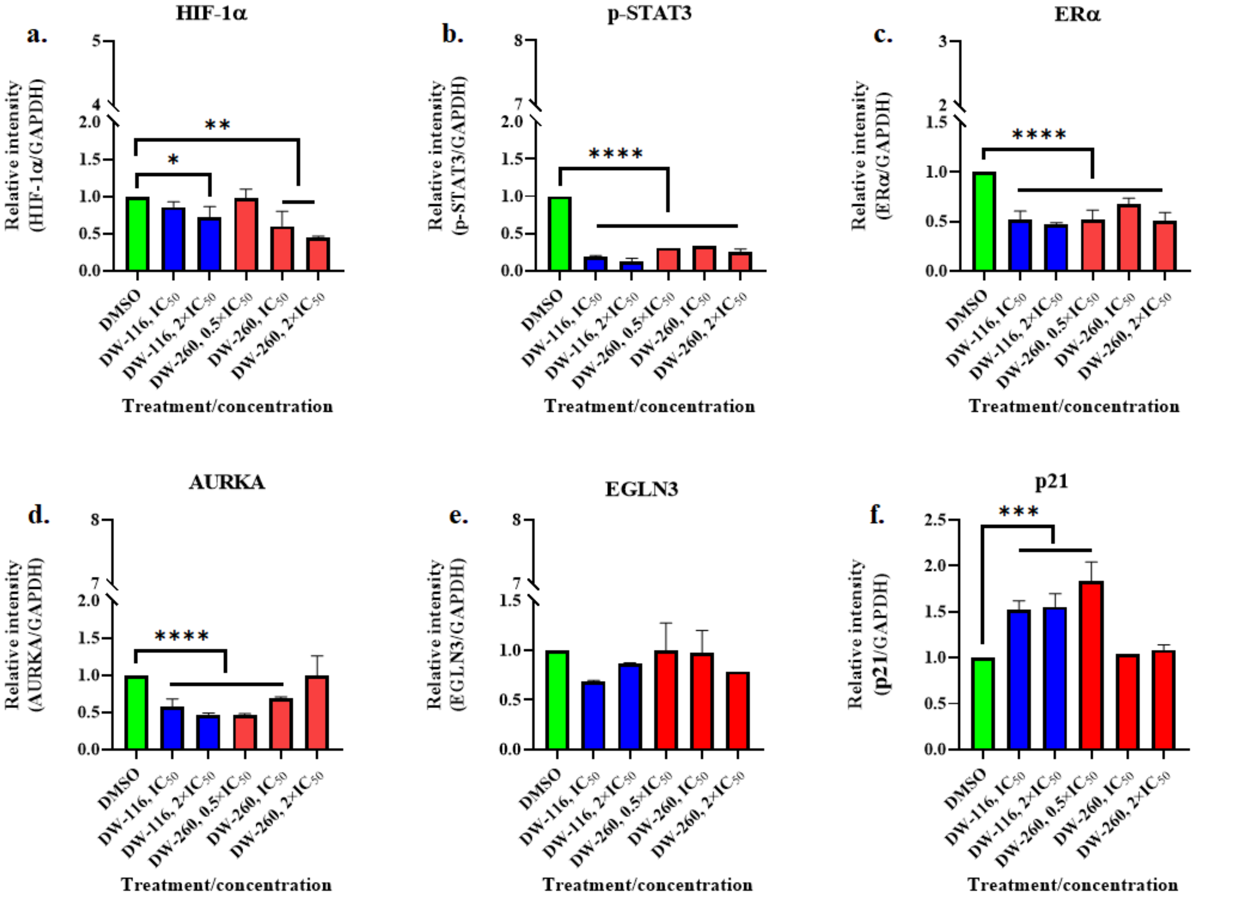


**Figure S7:** Western blot densitometric quantifications (**a**-**f**.) of MCF-7 breast cancer cells treated with compounds **DW-116** and **DW-260 (DW-122)** for 24 h. The quantifications are based on at least two independent experiments (bars show means plus standard deviations; *$p < 0.05$; **$p < 0.01$; ****$p < 0.0001$).

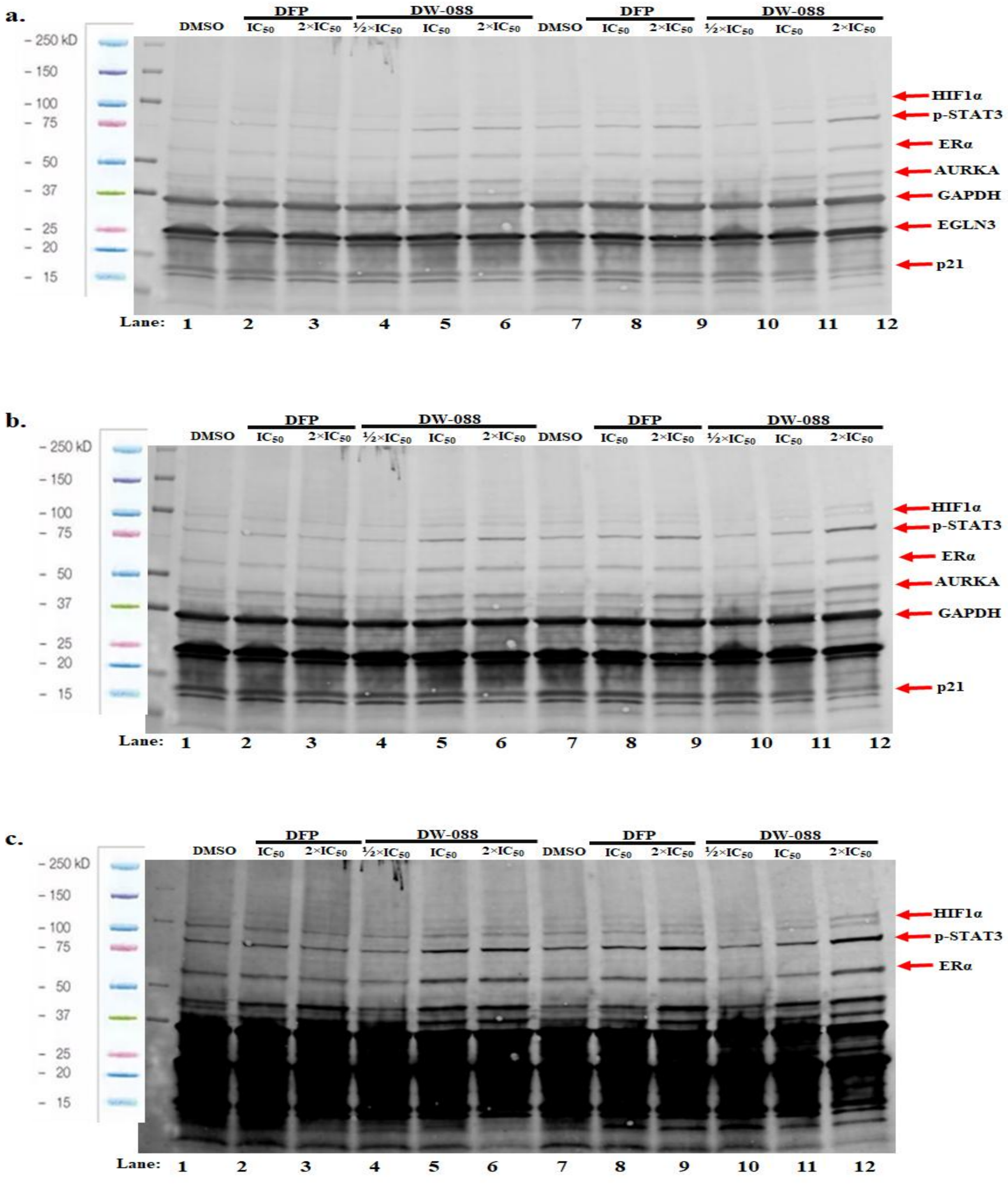


**Figure S8:** Representative full gels (**a.** – **c.**) of MCF-7 cells treated with deferiprone (DFP) and compound **DW-088** for 24 h, as indicated above.

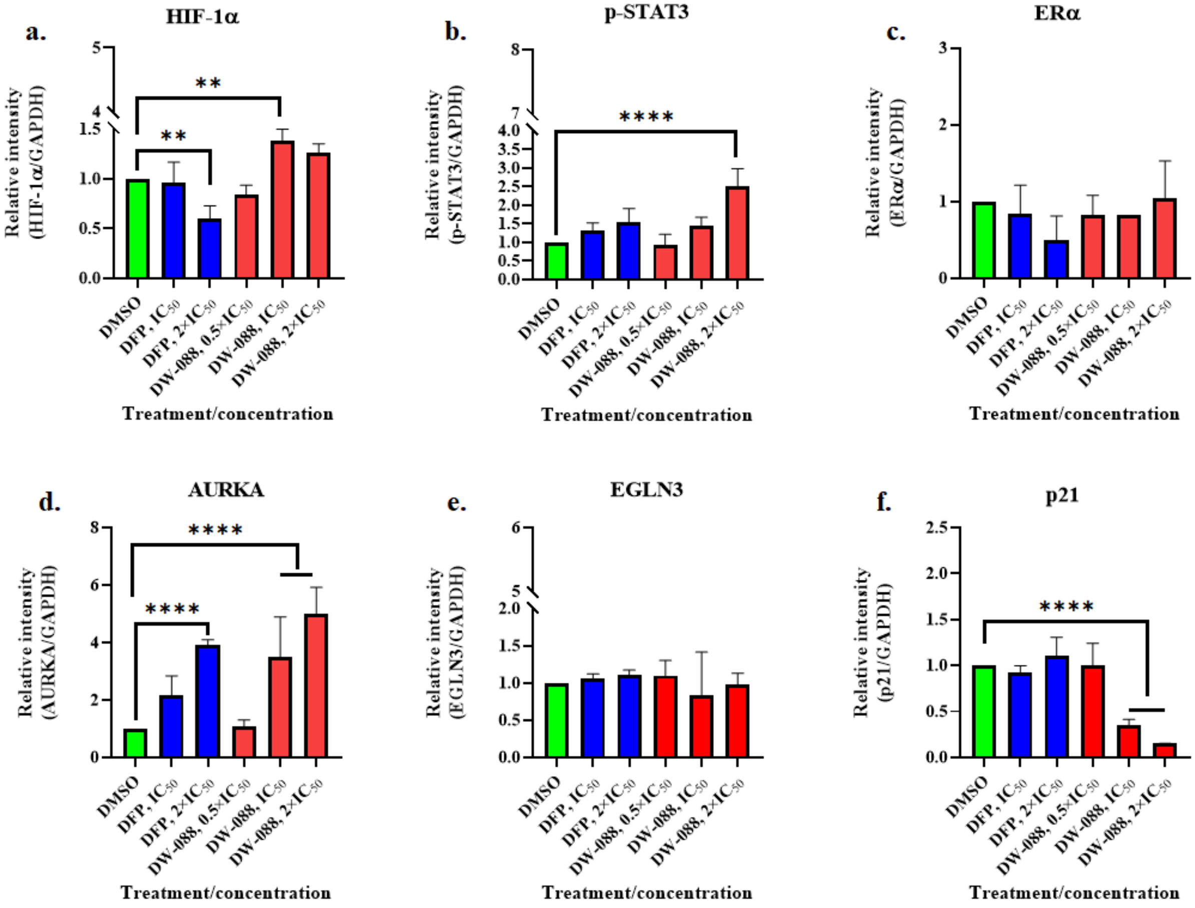


**Figure S9:** Western blot densitometric quantifications **(a-f.)** of MCF-7 breast cancer cells treated with deferiprone (DFP) and compound **DW-088** for 24 h. The quantifications are based on at least two independent experiments (bars show means plus standard deviations; **$p$ < 0.01; ****$p$ < 0.0001).

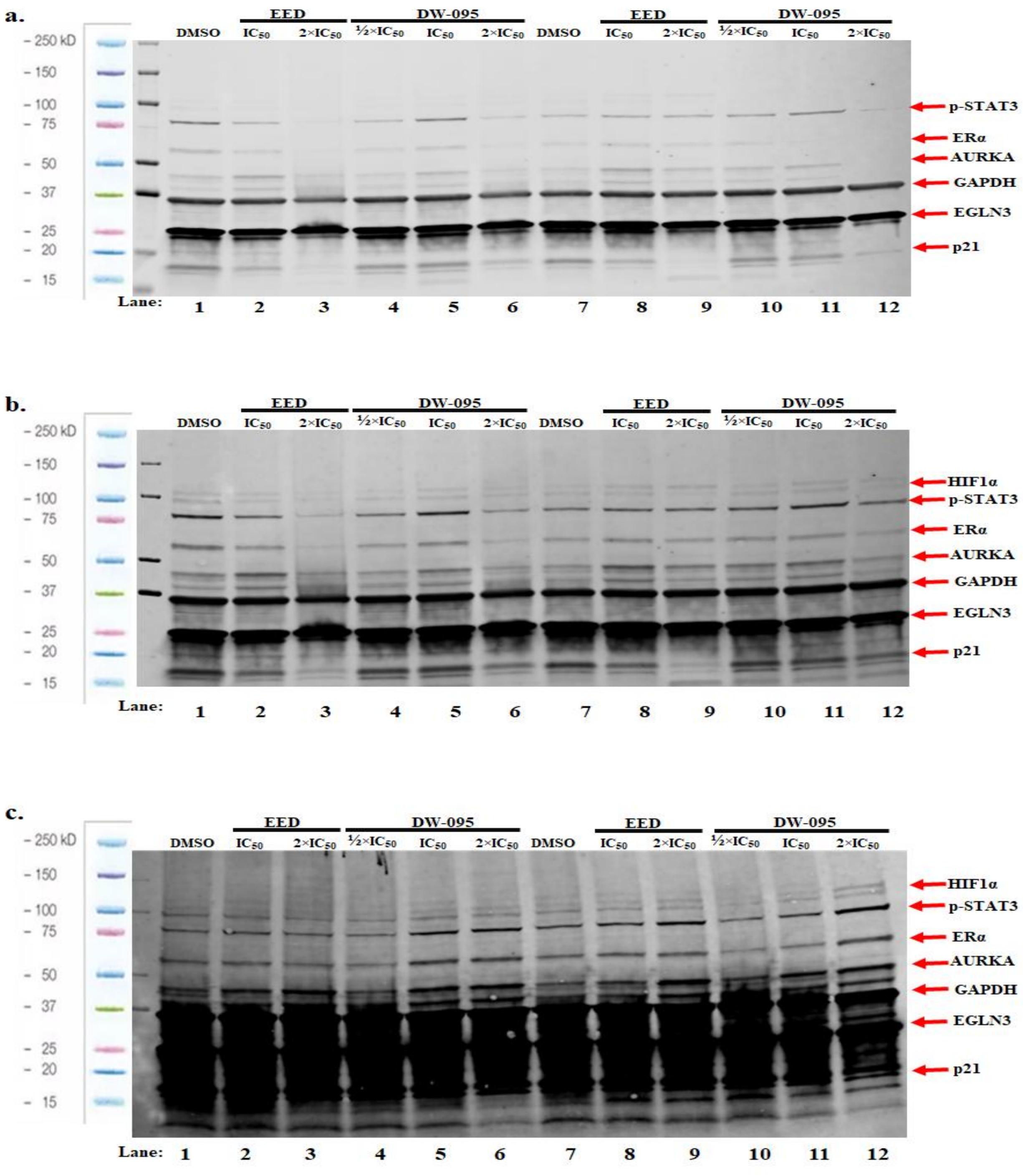


**Figure S10:** Representative full gels (**a**–**c**) of MCF-7 cells treated with ethinyl estradiol (EED) and compound **DW-095** for 24 h, as indicated above.

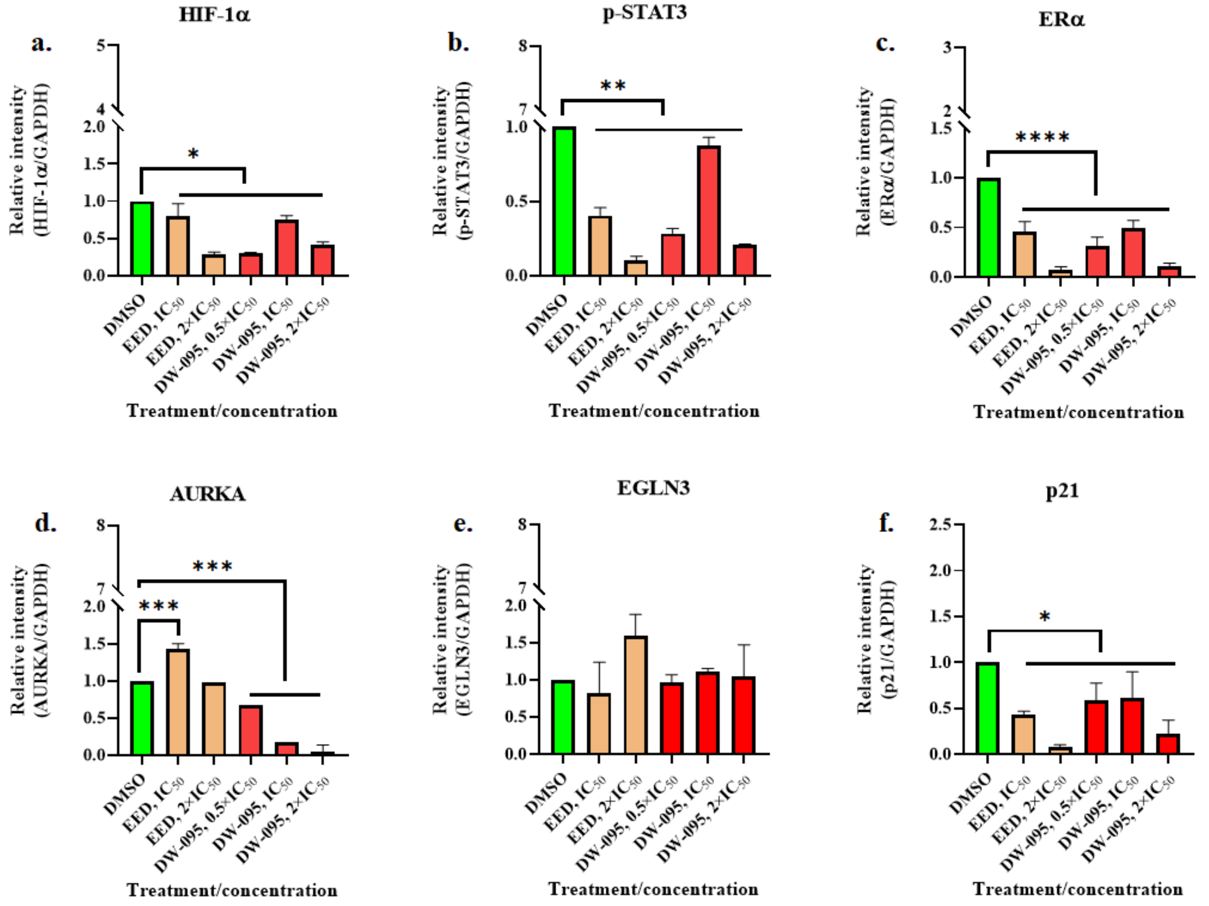


**Figure S11:** Western blot densitometric quantifications **(a-f)** of MCF-7 breast cancer cells treated with ethinyl estradiol (EED) and compound **DW-095** for 24 h. The quantifications are based on at least two independent experiments (bars show means plus standard deviations; *$p < 0.05$; **$p < 0.01$; ***$p < 0.001$; ****$p < 0.0001$).

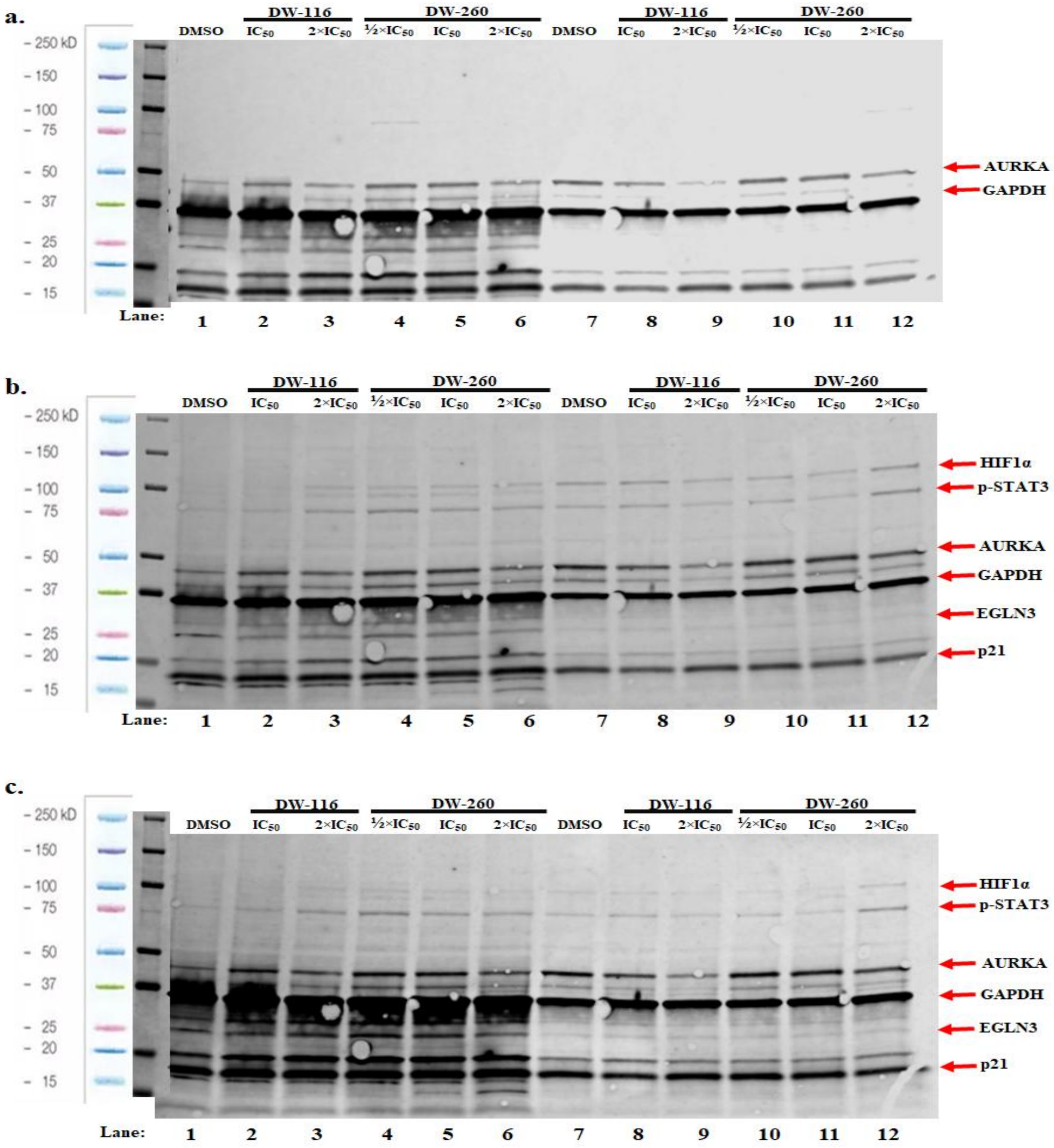


**Figure S12:** Representative full gels (**a**–**c**) of MDA-MB-231 cells treated with compounds **DW-116** and **DW-260** for 24 h, as indicated above.

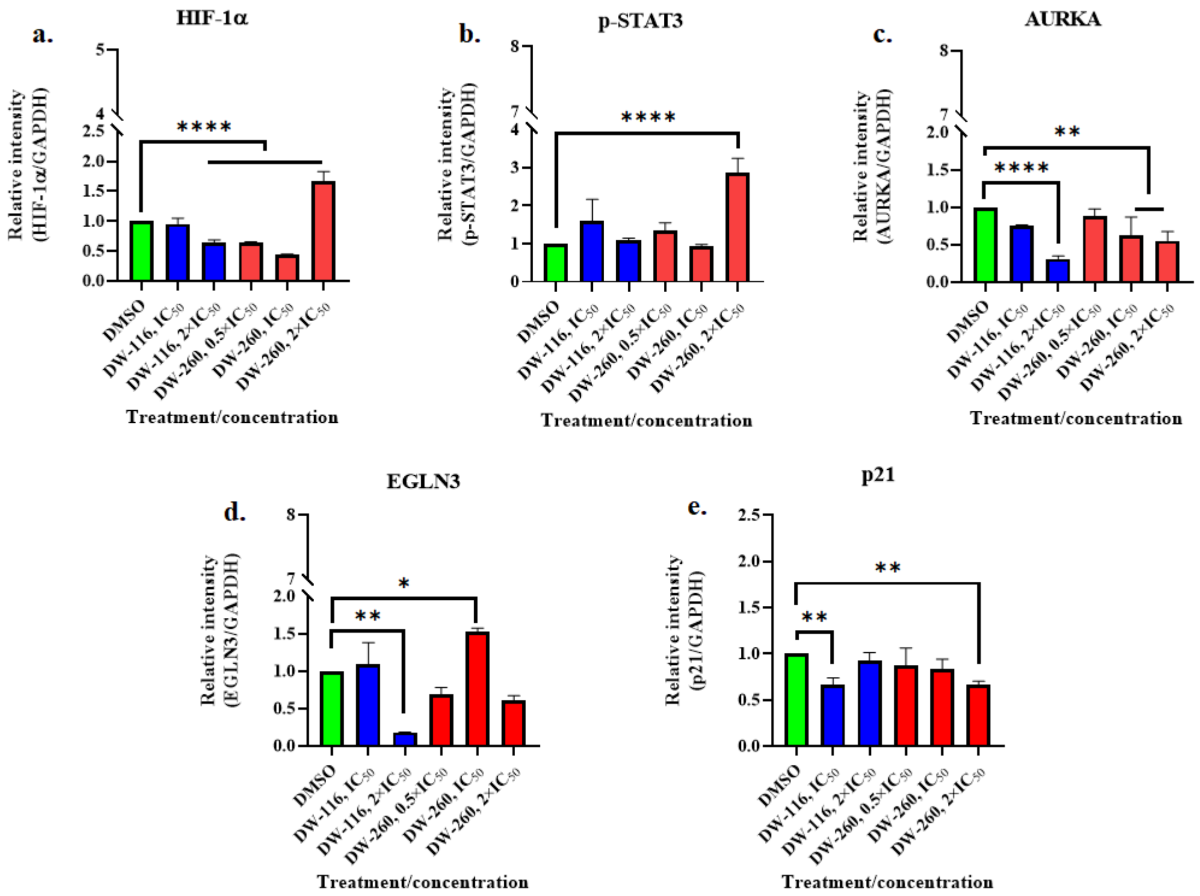


**Figure S13:** Western blot densitometric quantifications (**a**-**e**) of MDA-MB-231 breast cancer cells treated with compounds **DW-116** and **DW-260** for 24 h. The quantifications are based on at least two independent experiments (bars show means plus standard deviations; *$p < 0.05$; **$p < 0.01$; ****$p < 0.0001$).

**a.**

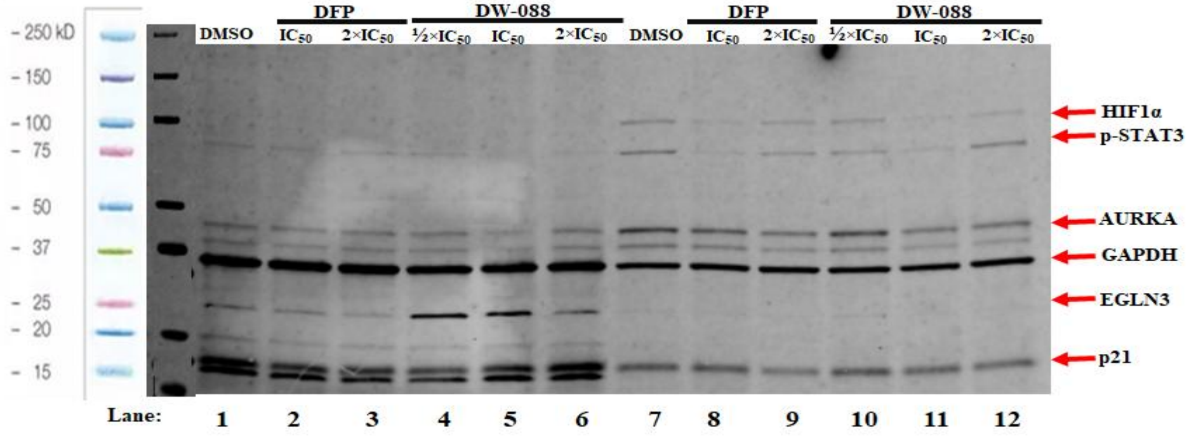


**b.**

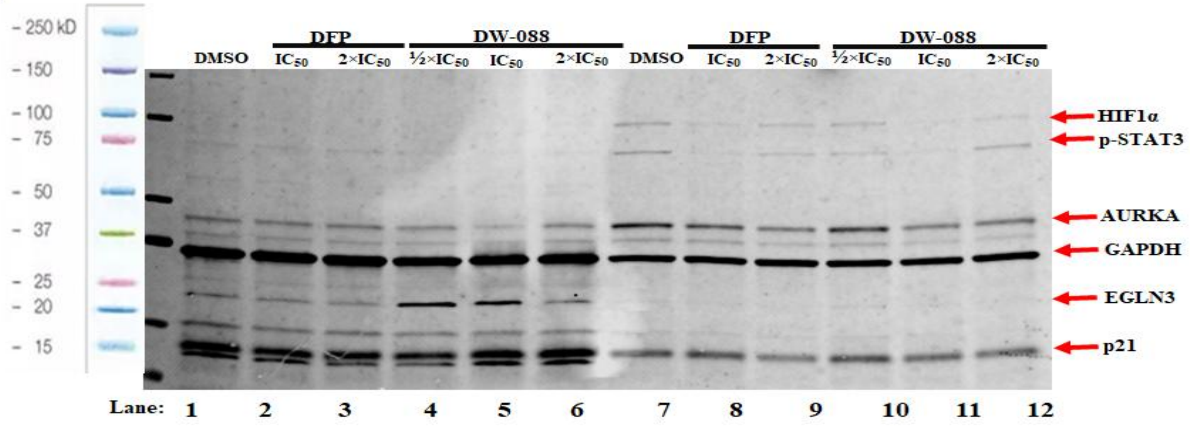


**Figure S14:** Representative full gels (**a.**–**b.**) of MDA-MB-231 cells treated with deferiprone (DFP) and compound **DW-088** for 24 h, as indicated above.

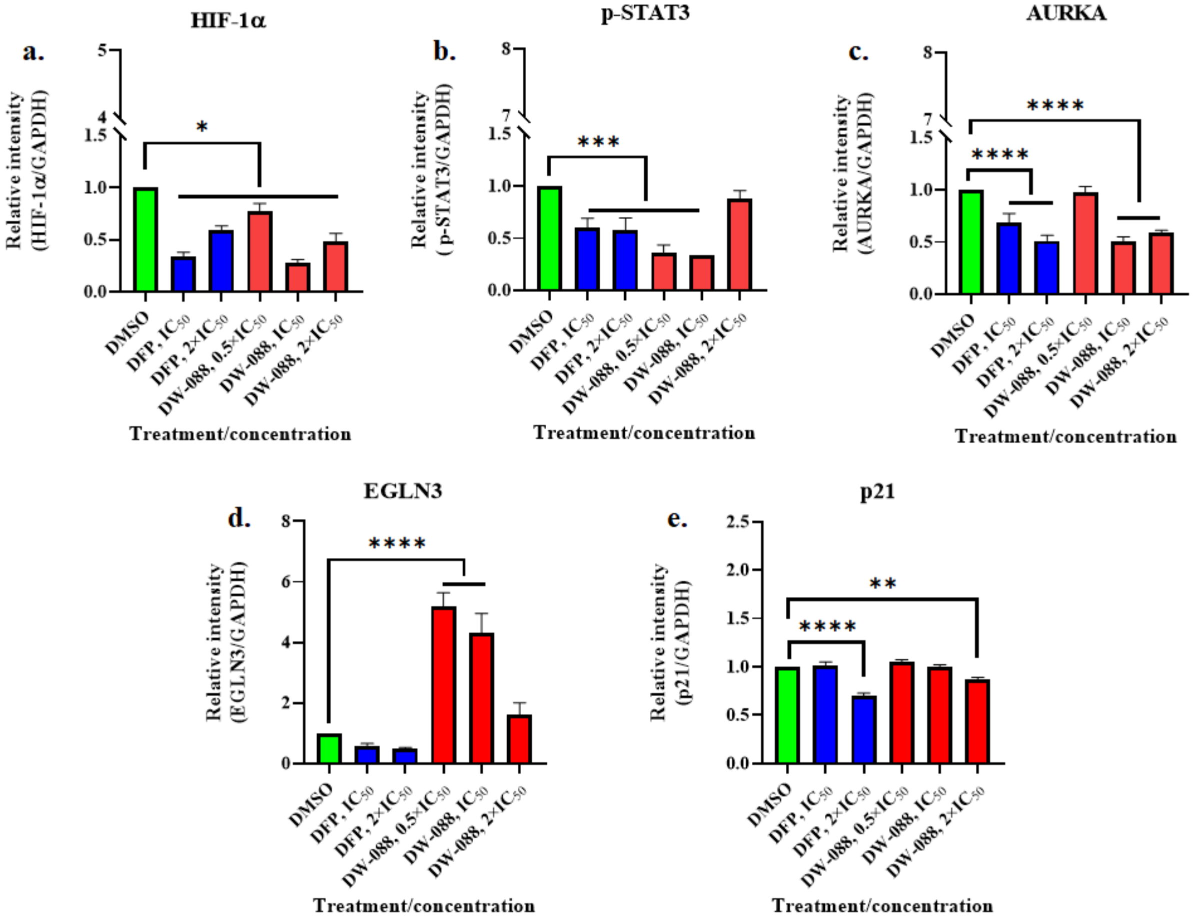


**Figure S15:** Western blot densitometric quantifications **(a.–e.**) of MDA-MB-231 breast cancer cells treated deferiprone (DFP) and compound **DW-088** for 24 h. The quantifications are based on at least two independent experiments (bars show means plus standard deviations; *$p < 0.05$; **$p < 0.01$; ***$p < 0.001$; ****$p < 0.0001$).

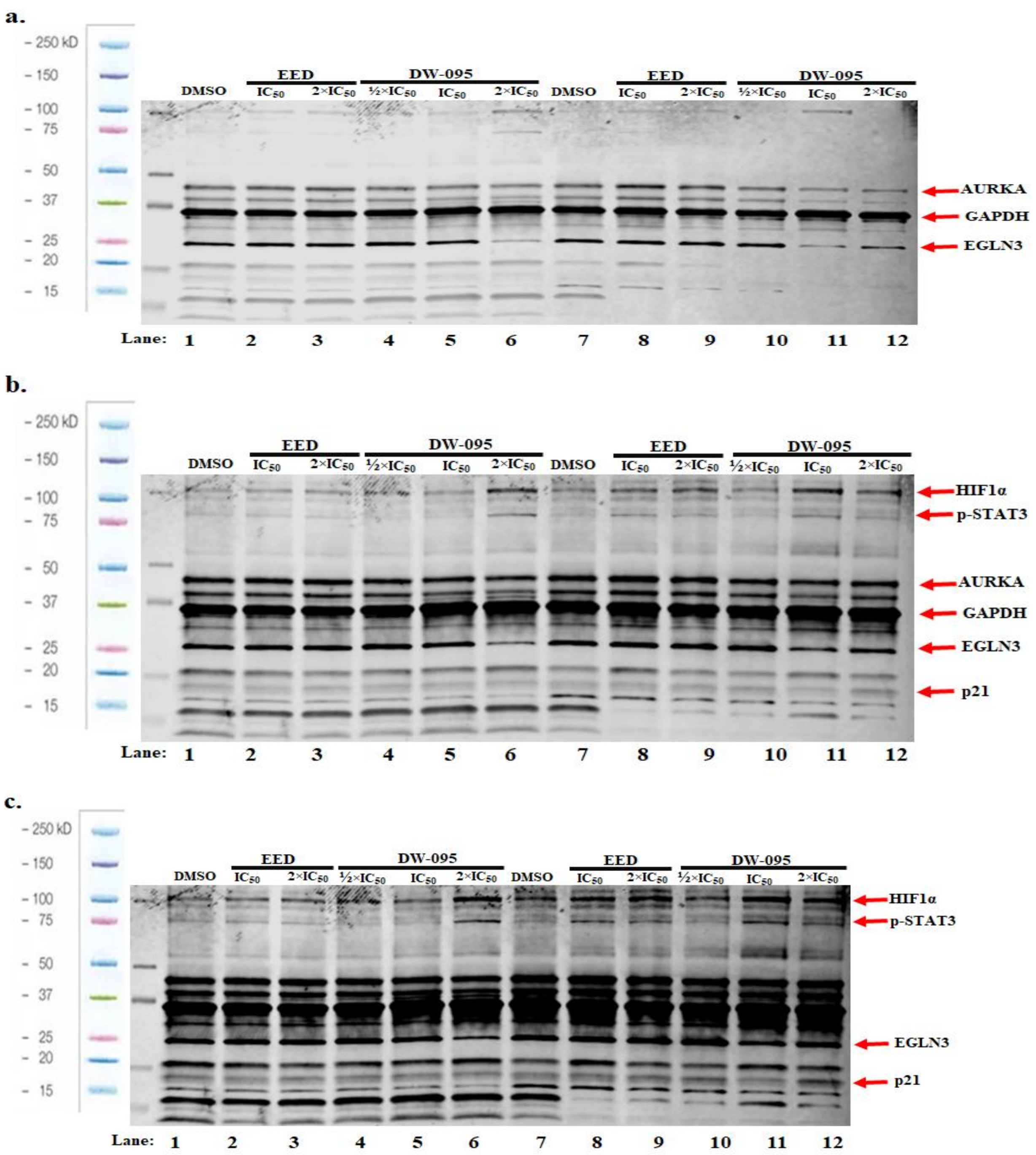


**Figure S16:** Representative full gels (**a.–c.**) of MDA-MB-231 cells treated with ethinyl estradiol (EED) and compound **DW-095** for 24 h, as indicated above.

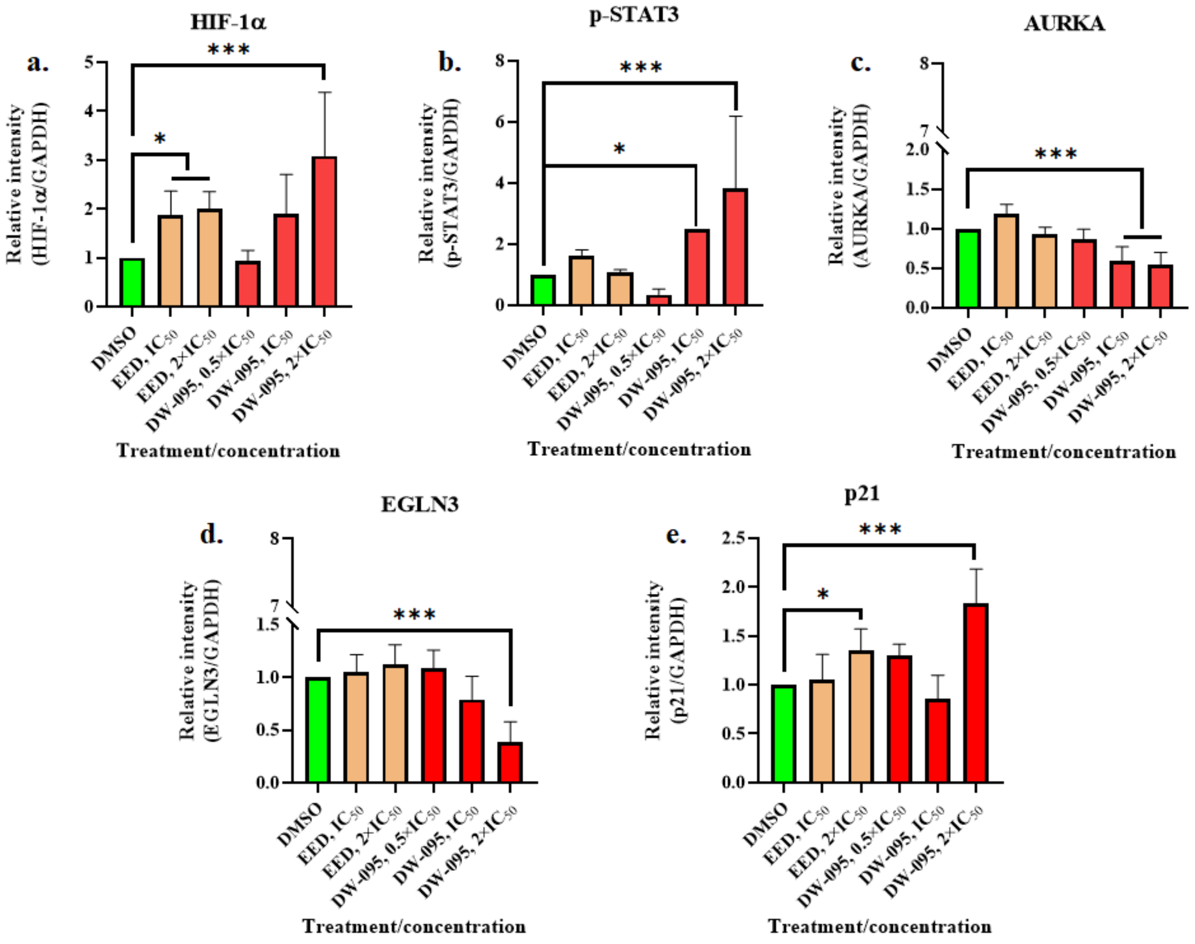


**Figure S17:** Western blot densitometric quantifications **(a.–e.)** of MDA-MB-231 breast cancer cells treated ethinyl estradiol (EED) and compound **DW-095** for 24 h. The quantifications are based on at least two independent experiments (bars show means plus standard deviations; $*p < 0.05$; $***p < 0.001$).

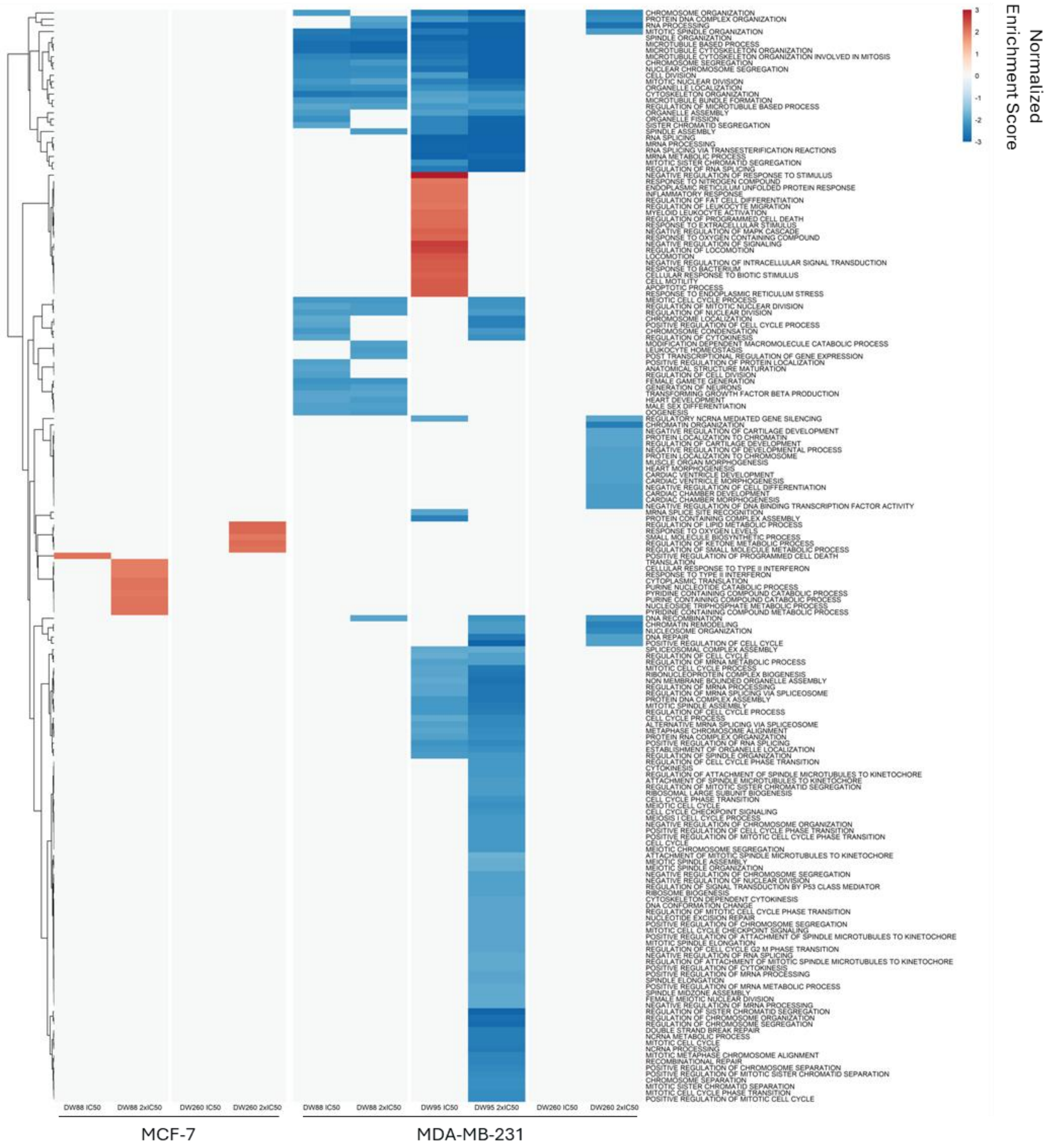


**Figure S18:** Gene Ontology Biological Processes (GOBP) Gene Set Enrichment Analysis (GSEA) of EED-KDMi and Tam-KDMi treatment to MCF-7 and MDA-MB-231 cells. Heatmap of normalized enrichment scores (NES) ($p < 0.01$, FDR $< 0.1$). Minimal effect was seen with **DW-088** and **DW-260** in MCF-7 cells. In MDA-MBA-231 cells, mitotic spindle organization (NES= -2.7, **DW-088**, -3.4, **DW-095**, -2.1, **DW-260**), chromosome organization (NES= -4.1, **DW-095**, -2.4, **DW-260**), and DNA recombination (NES= -1.9, **DW-088**, -2.2, **DW-095**, -2.2, **DW-260**) were among the gene sets negatively enriched by **DW-088, DW-095**, and **DW-260** at 2x $IC_{50}$.

### NMR Spectra

#### $^{1}H$ (700 MHz, $CDCl_3$) and $^{13}C$ (176 MHz, $CDCl_3$) NMR spectrum of compound 1

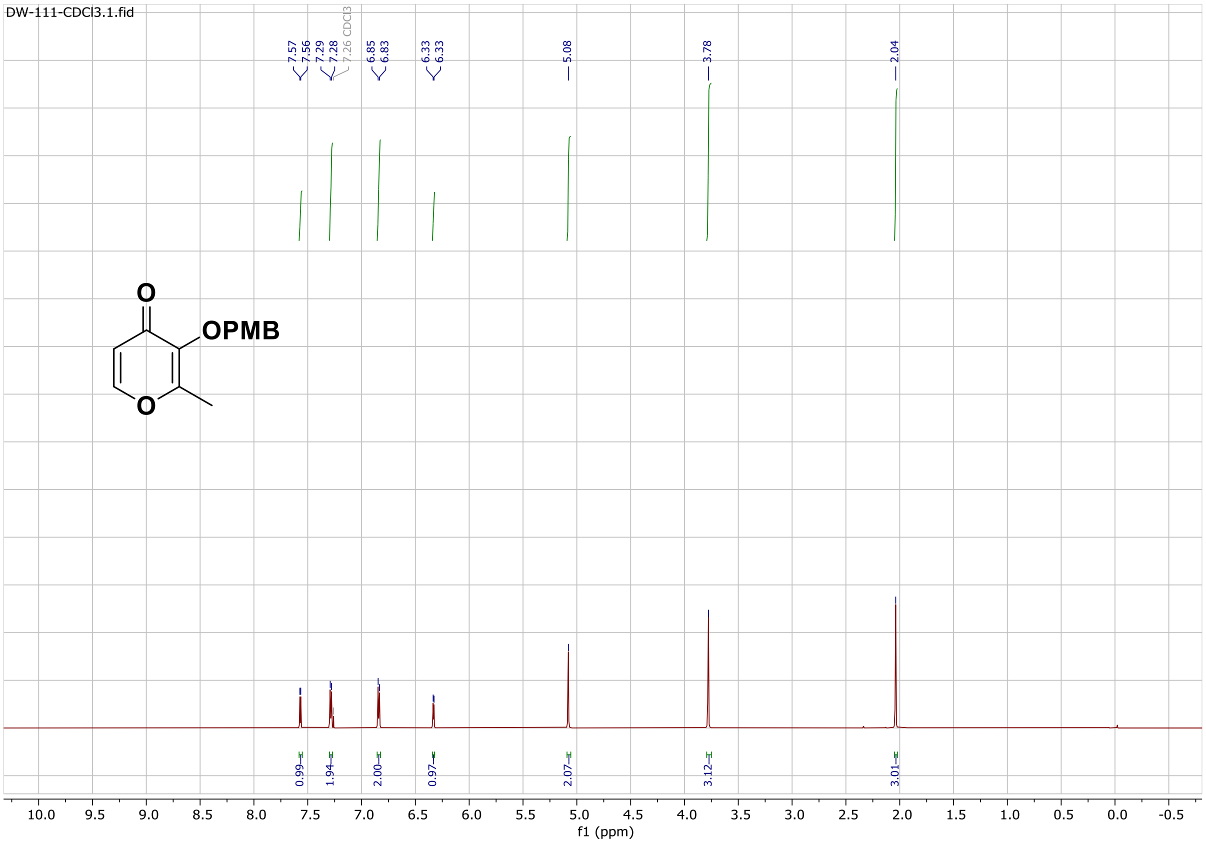


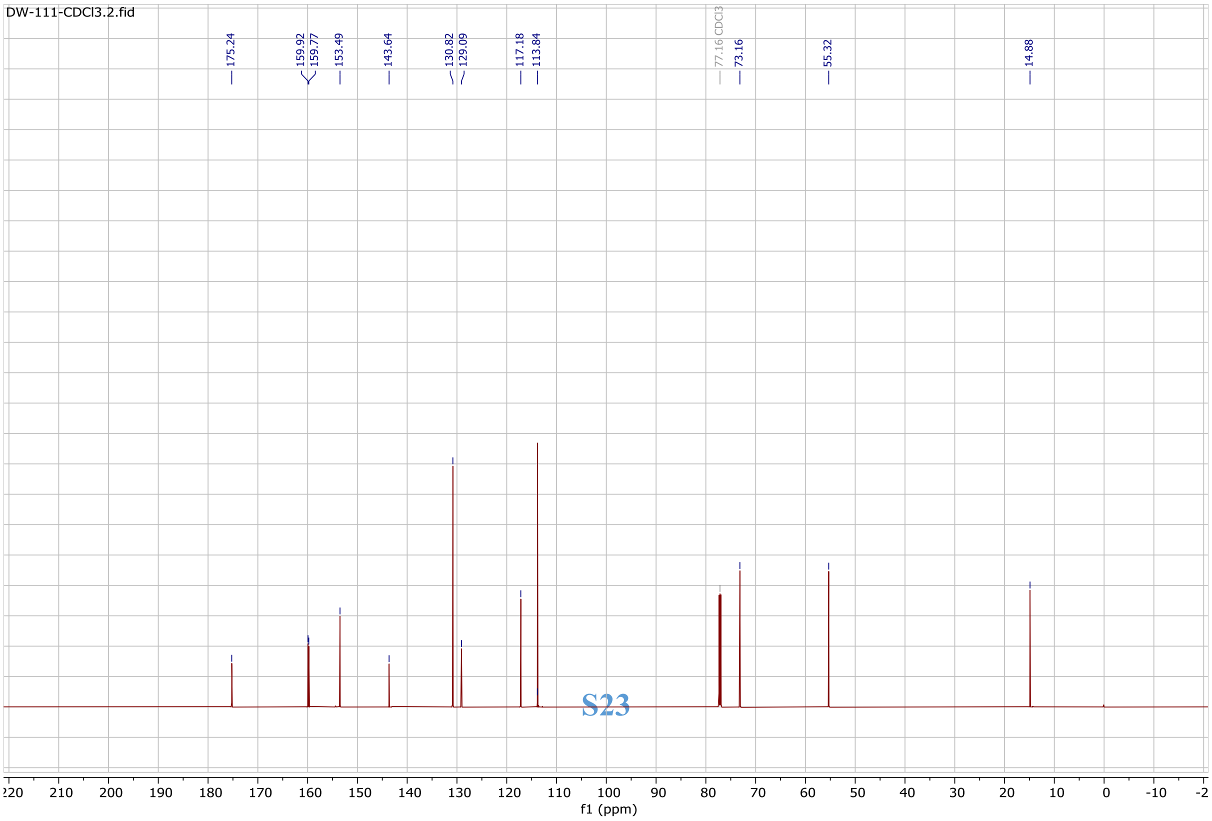

#### $^1$H (400 MHz, $CDCl_3$) and $^{13}$C (176 MHz, $CDCl_3$) NMR spectrum of compound 3a

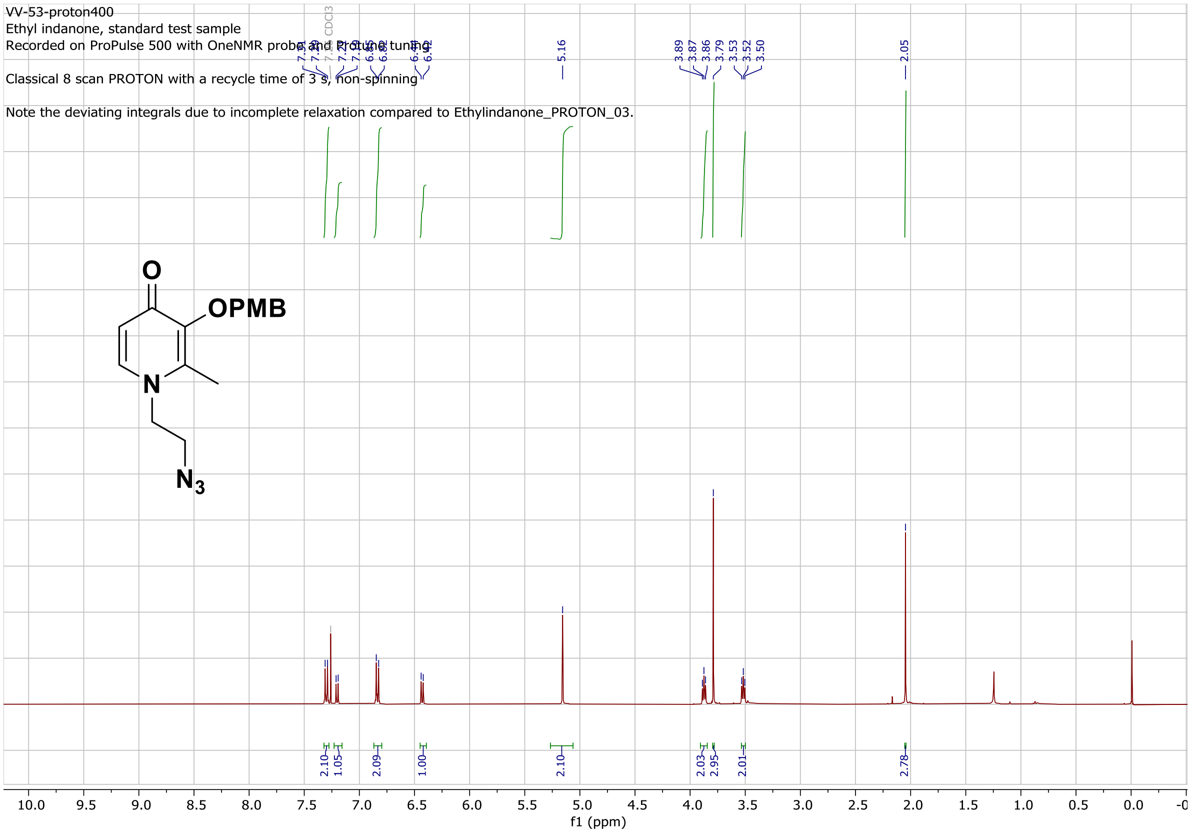


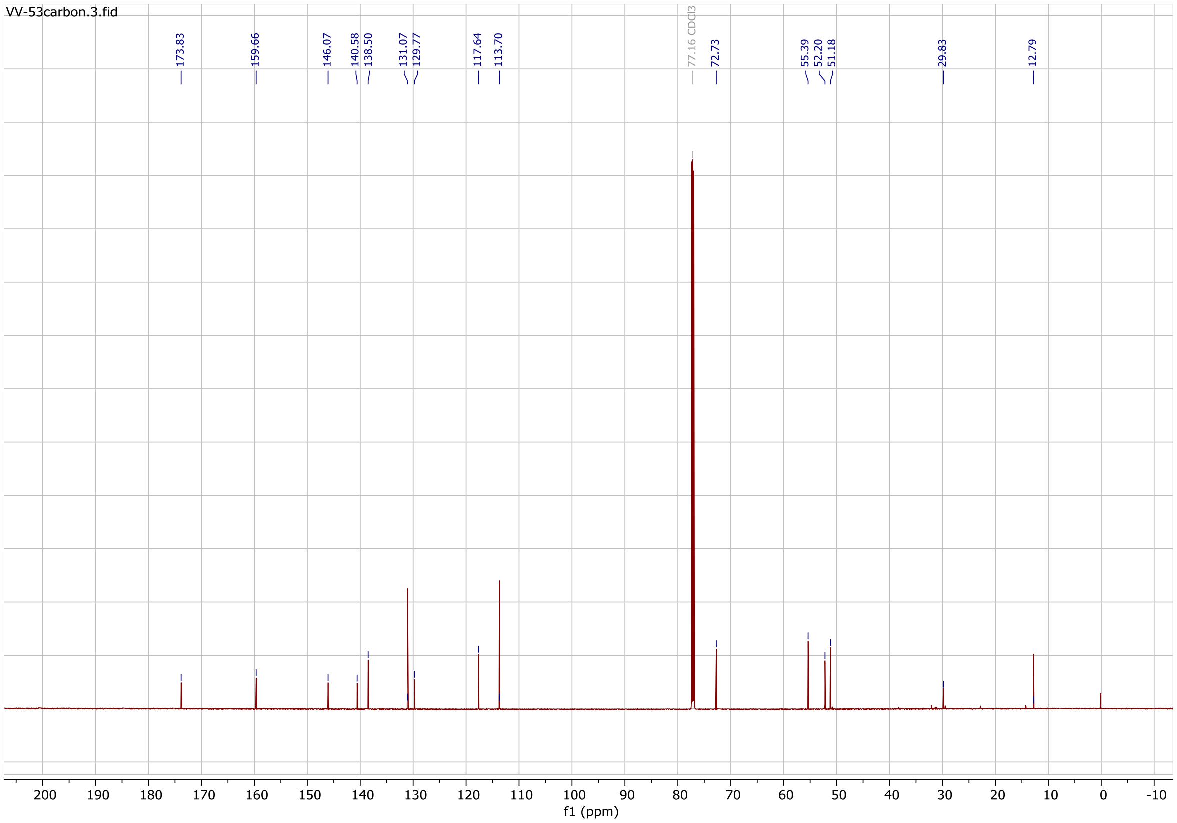

### 1H (400 MHz, CDCl3) and 13C (176 MHz, CDCl3) NMR spectrum of compound 3b

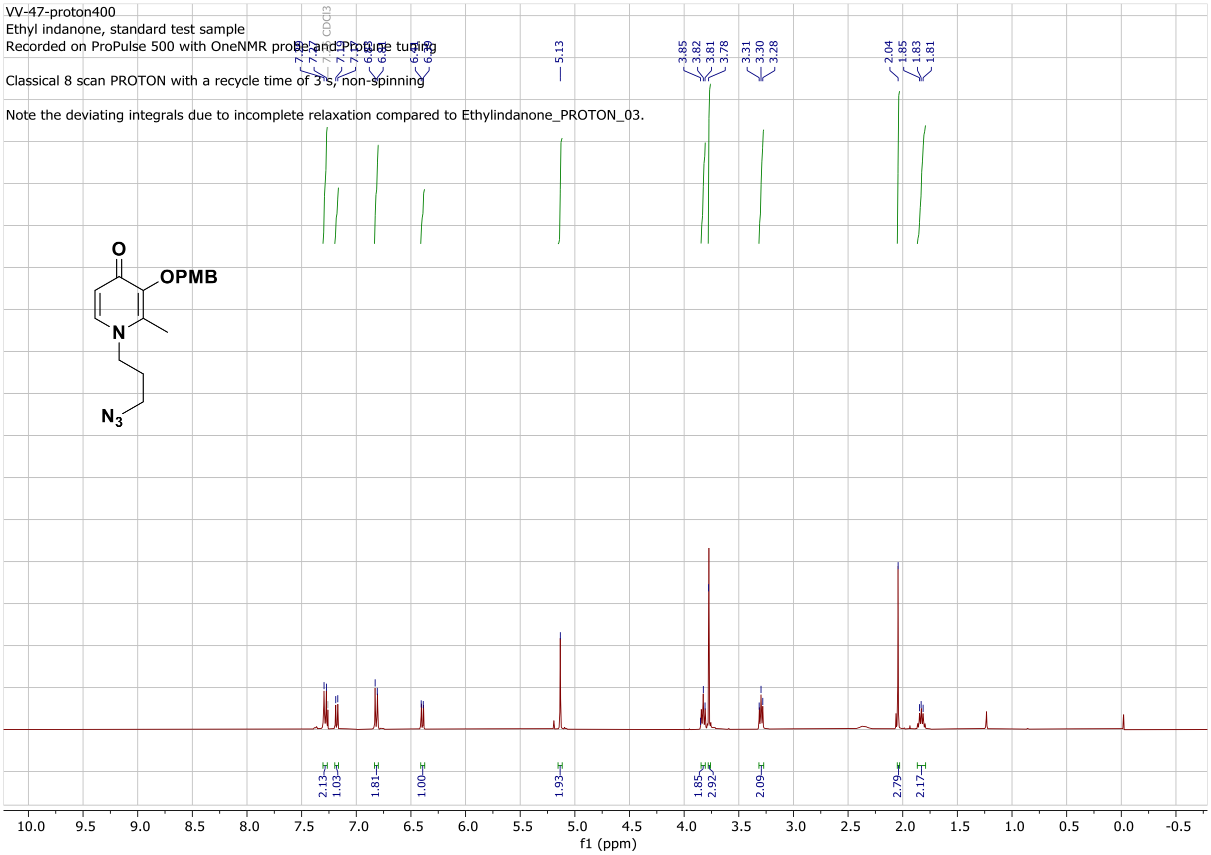


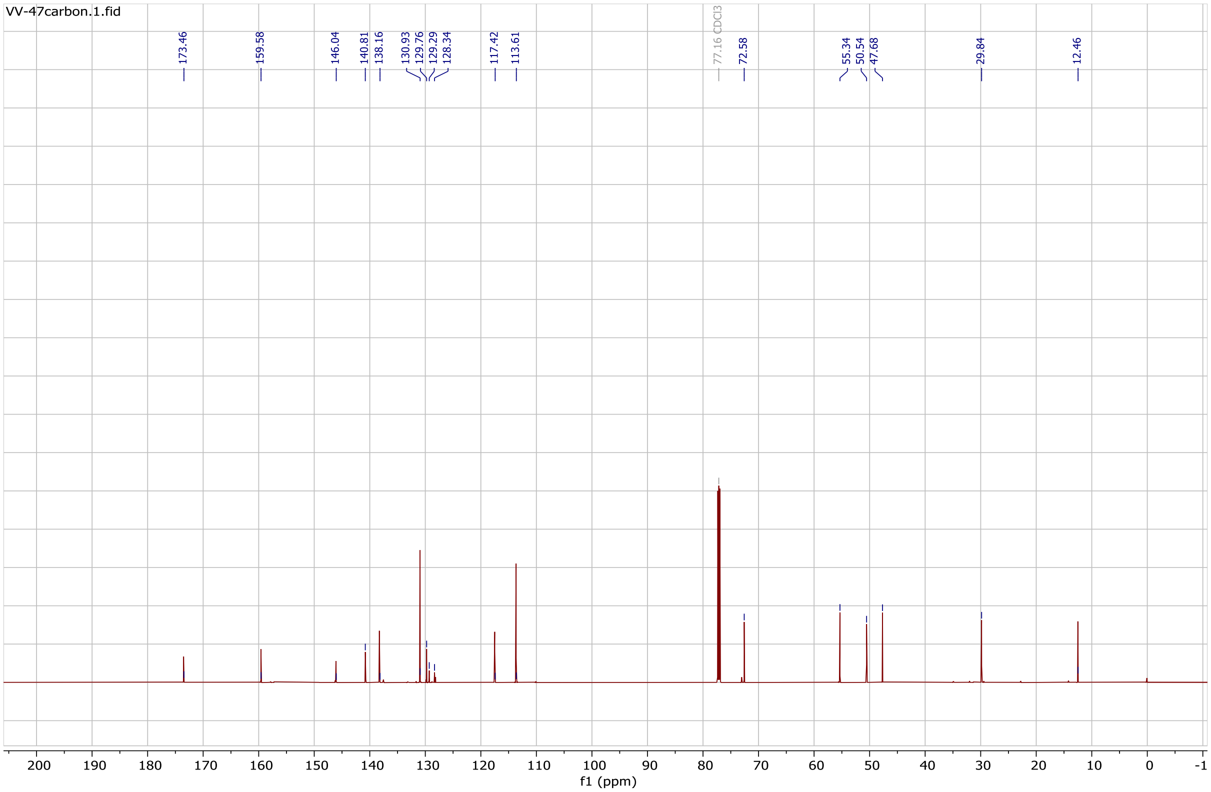

#### $^{1}H$ (400 MHz, $CDCl_3$) and $^{13}C$ (176 MHz, $CDCl_3$) NMR spectrum of compound 3c

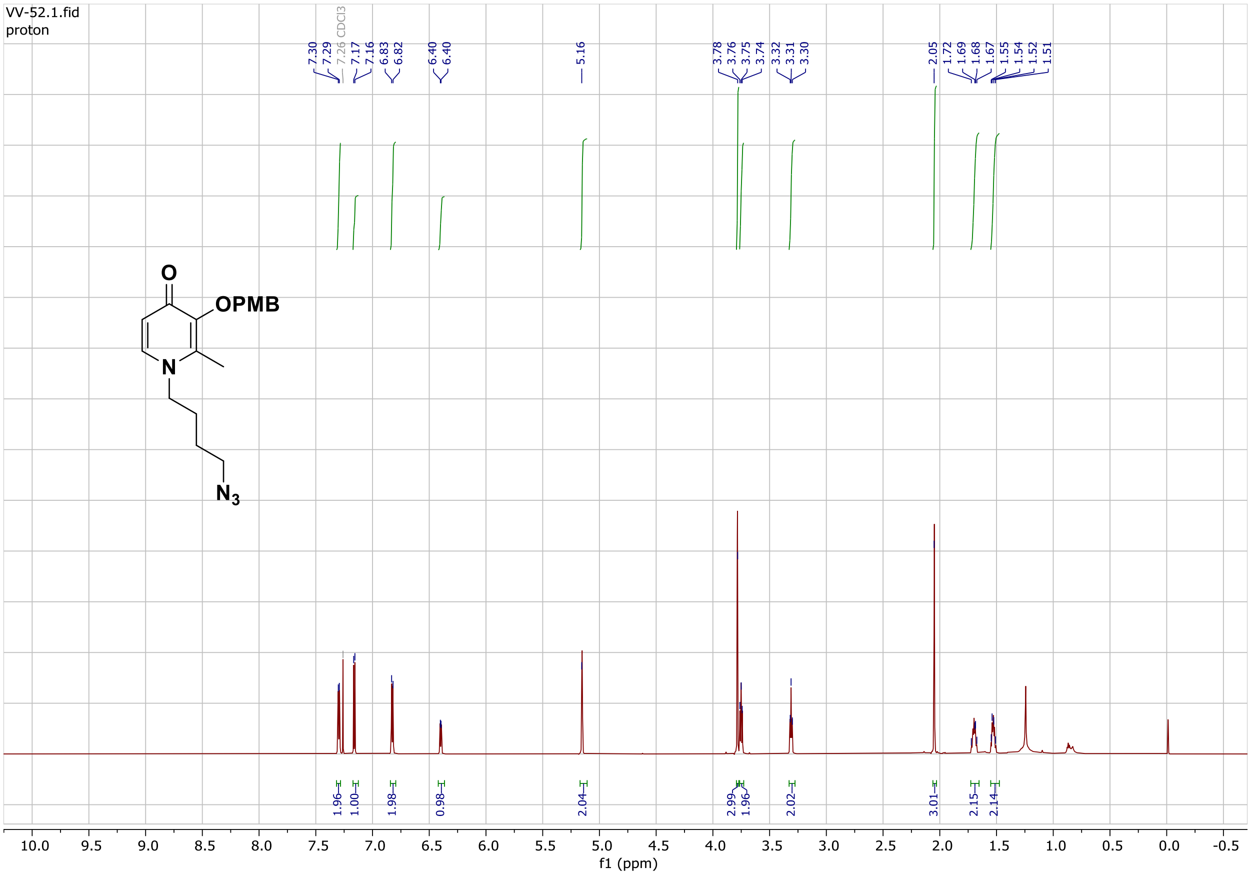


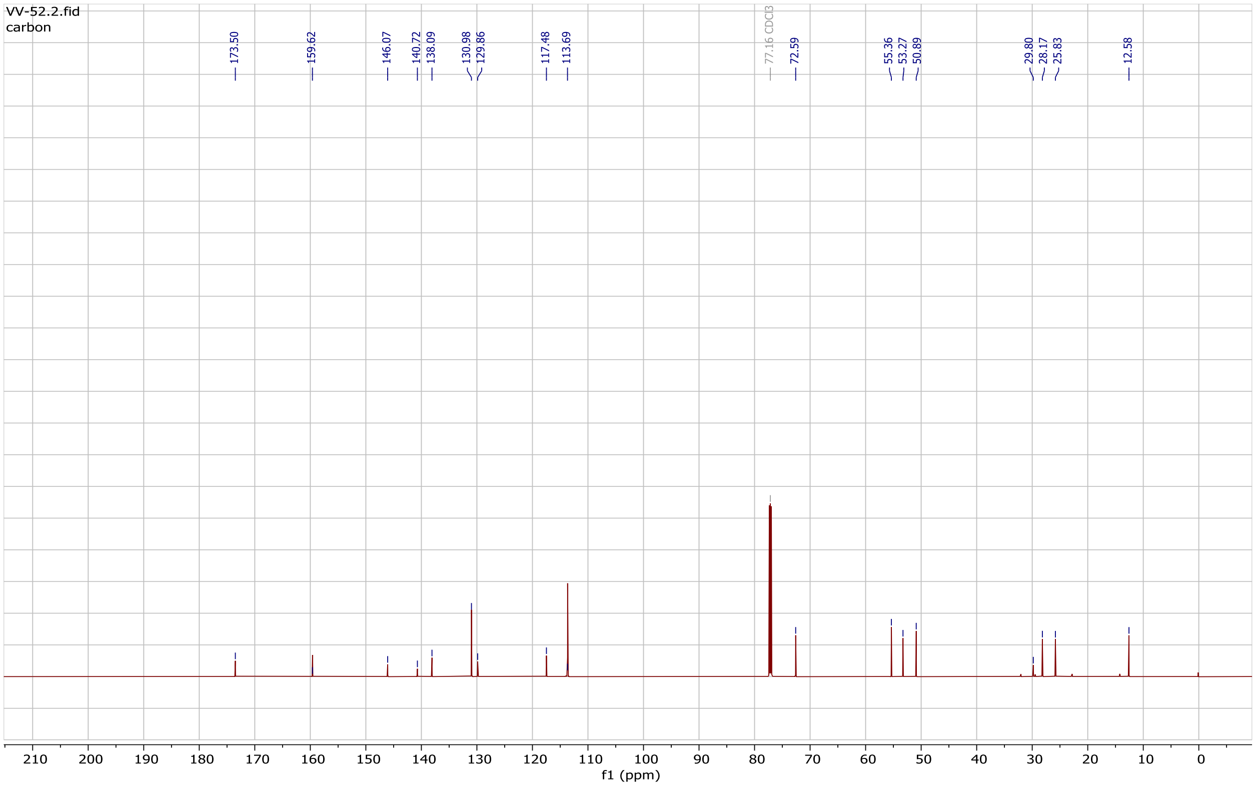

#### $^{1}H$ (400 MHz, $CDCl_3$) and $^{13}C$ (176 MHz, $CDCl_3$) NMR spectrum of compound 3d

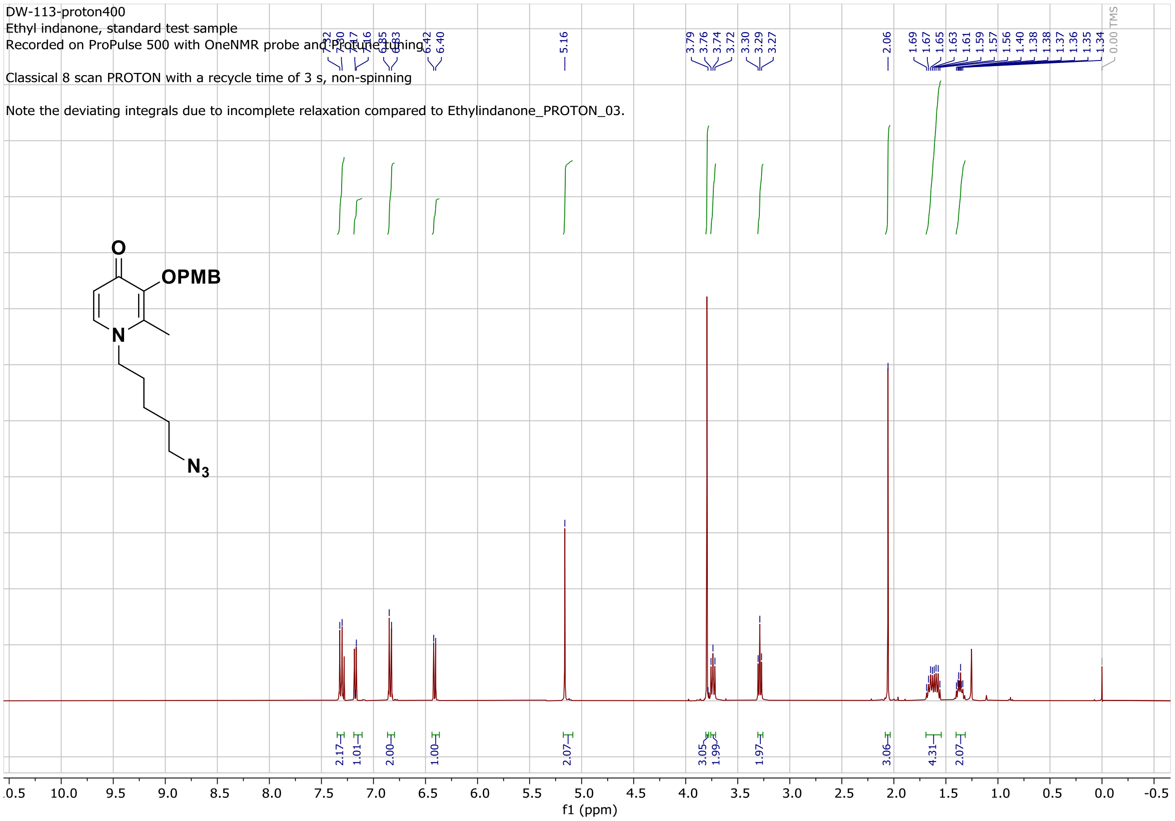


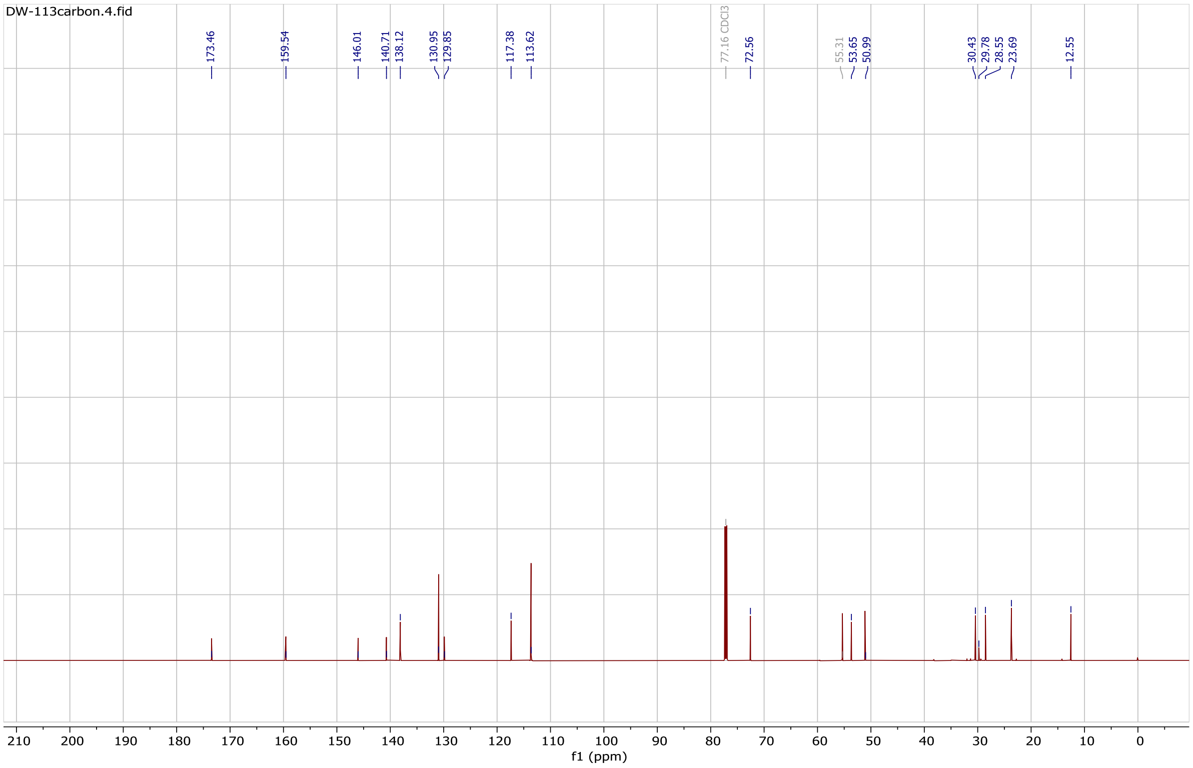

### $^{1}H$ (400 MHz, $CDCl_3$) and $^{13}C$ (176 MHz, $CDCl_3$) NMR spectrum of compound 3e

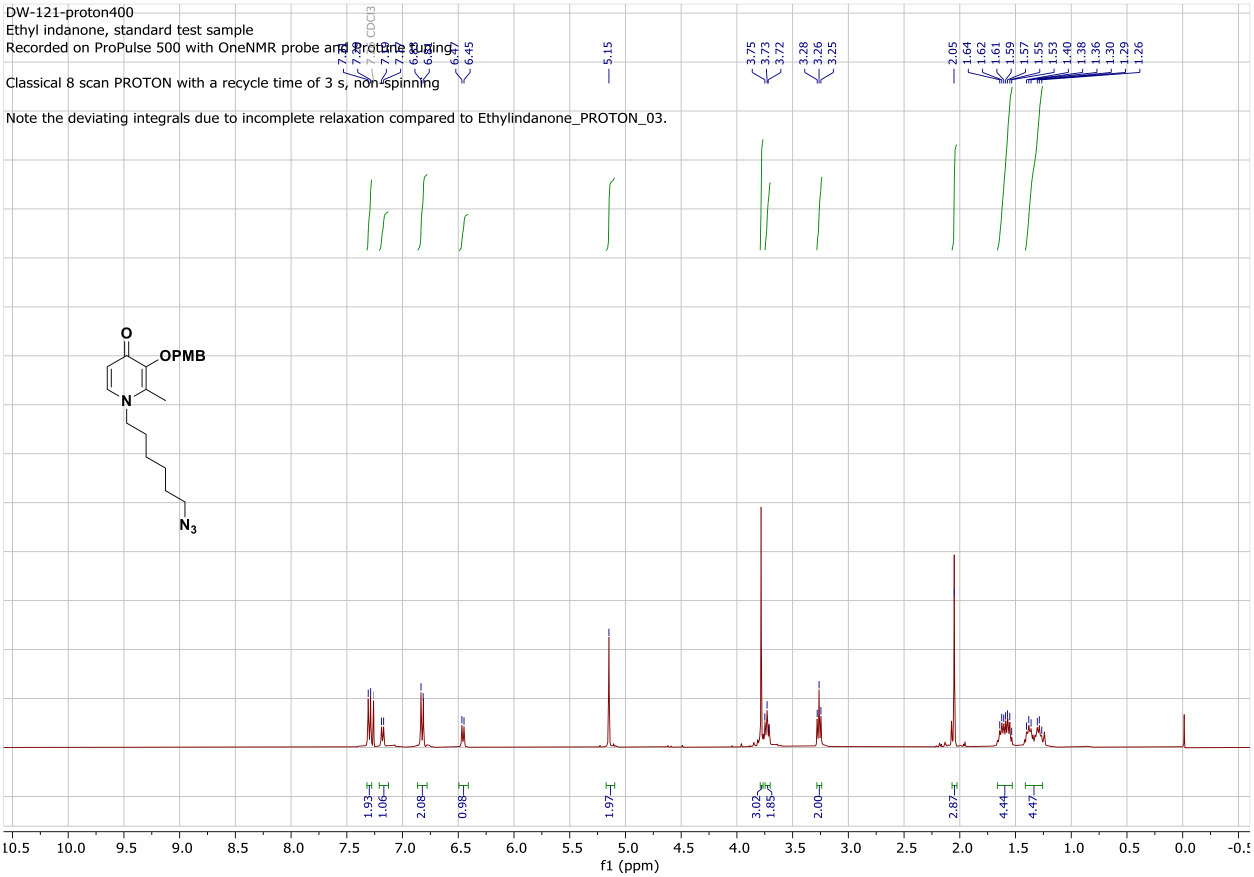


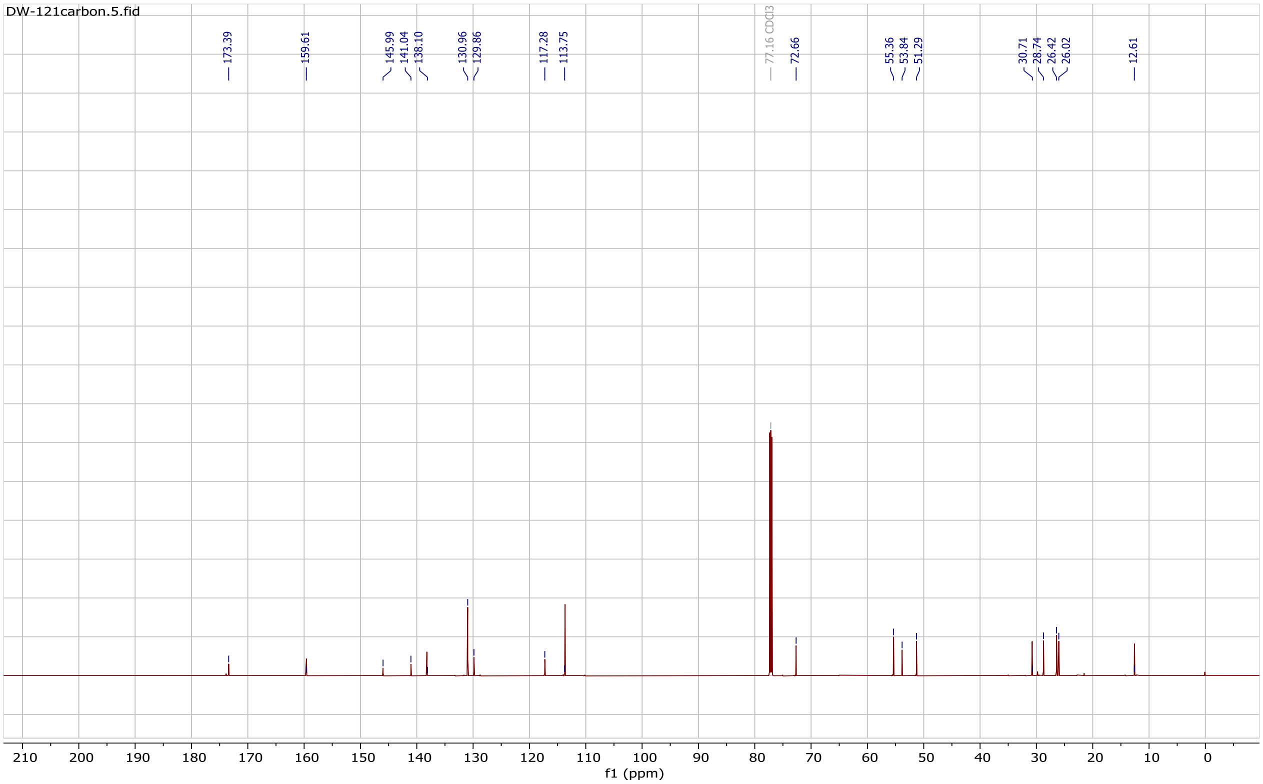

#### $^{1}H$ (400 MHz, $CDCl_3$) and $^{13}C$ (176 MHz, $CDCl_3$) NMR spectrum of compound 3f

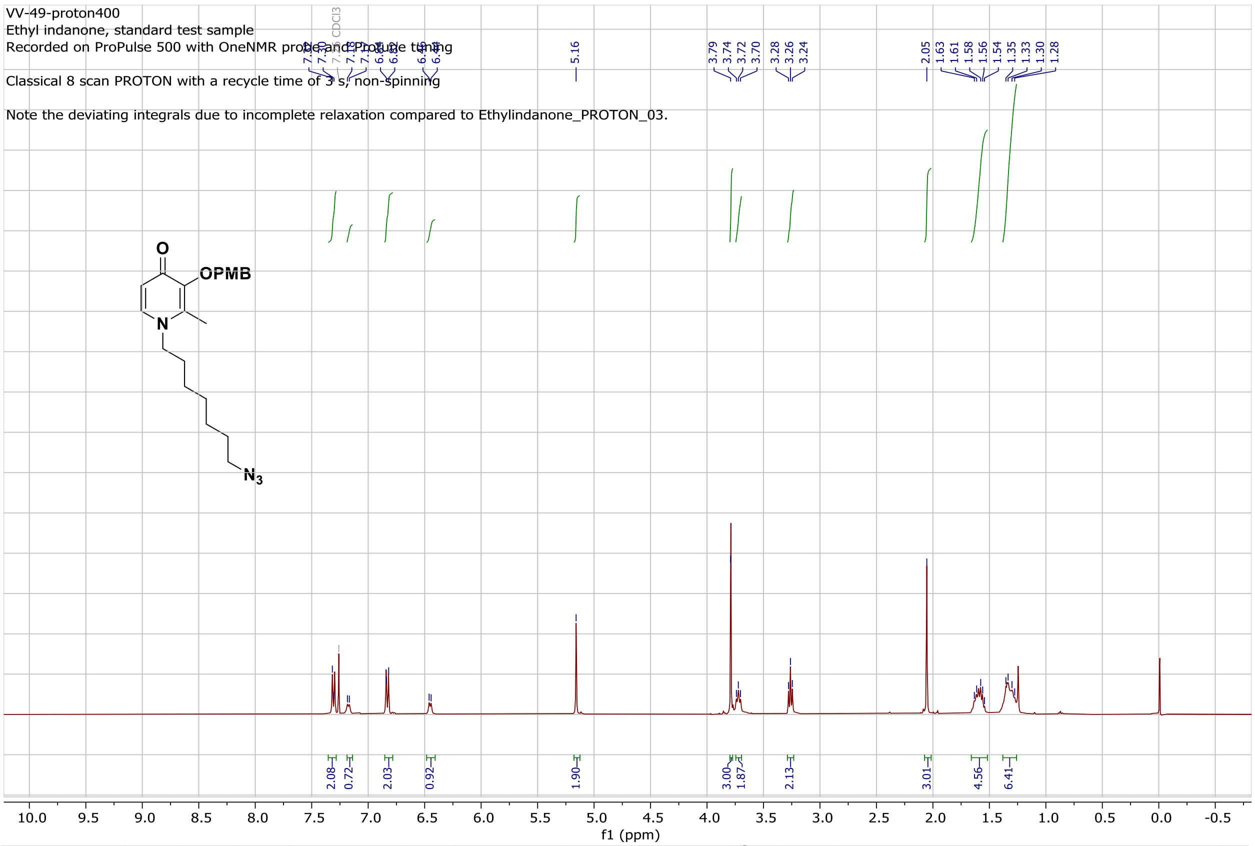


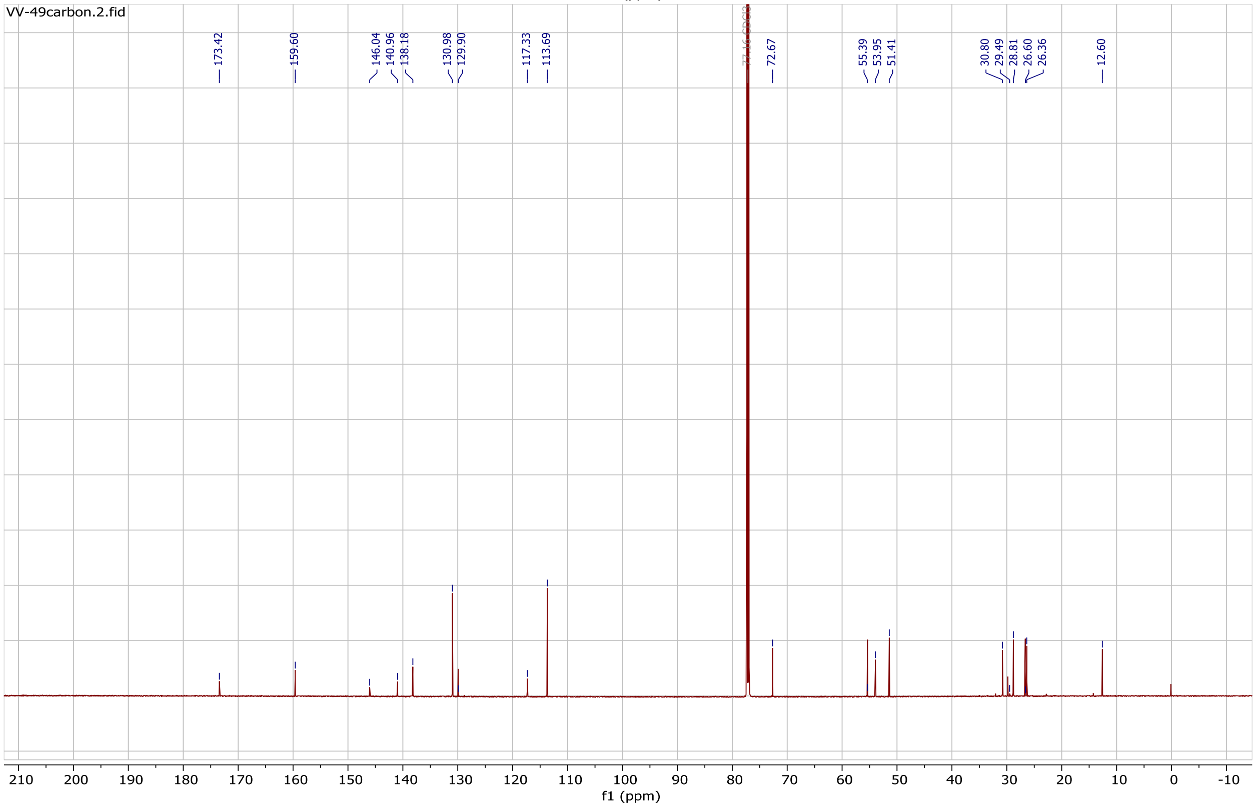

#### $^1H$ (700 MHz, $CDCl_3$) and $^{13}C$ (176 MHz, $CDCl_3$) NMR spectrum of compound 3g

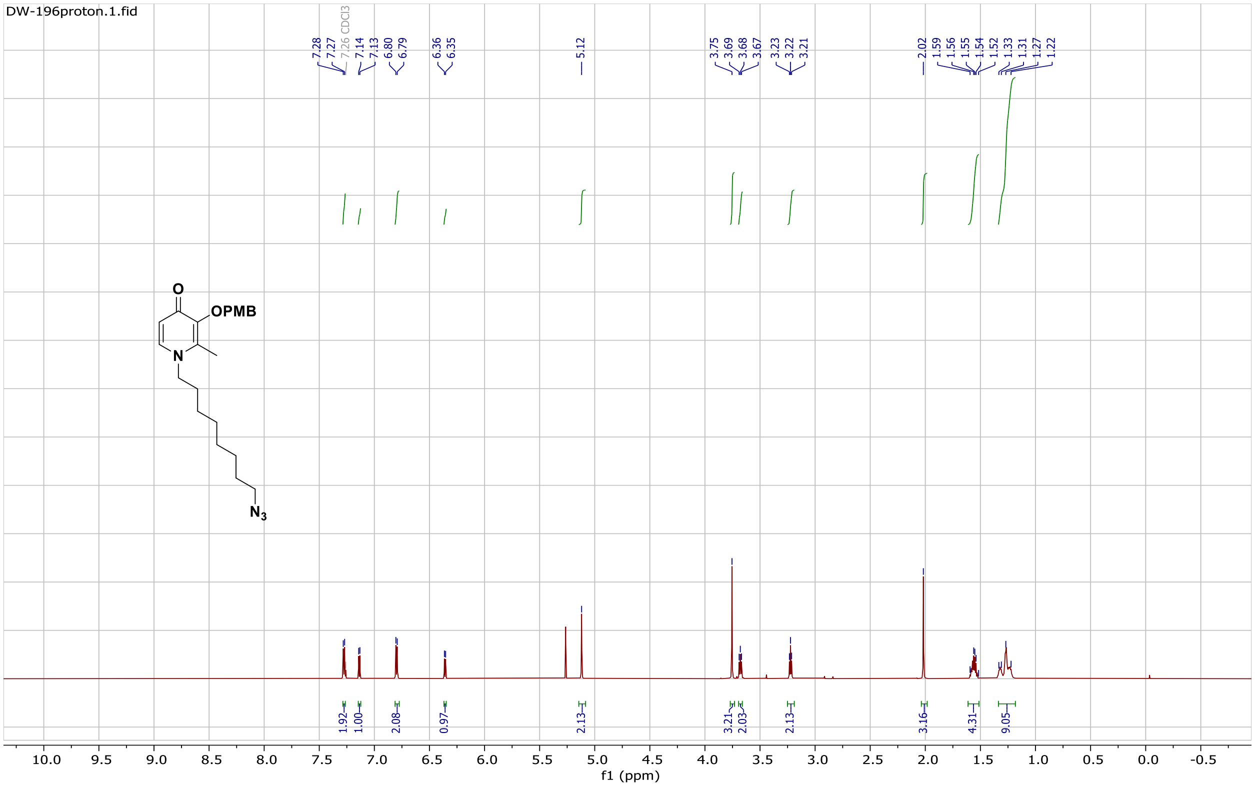


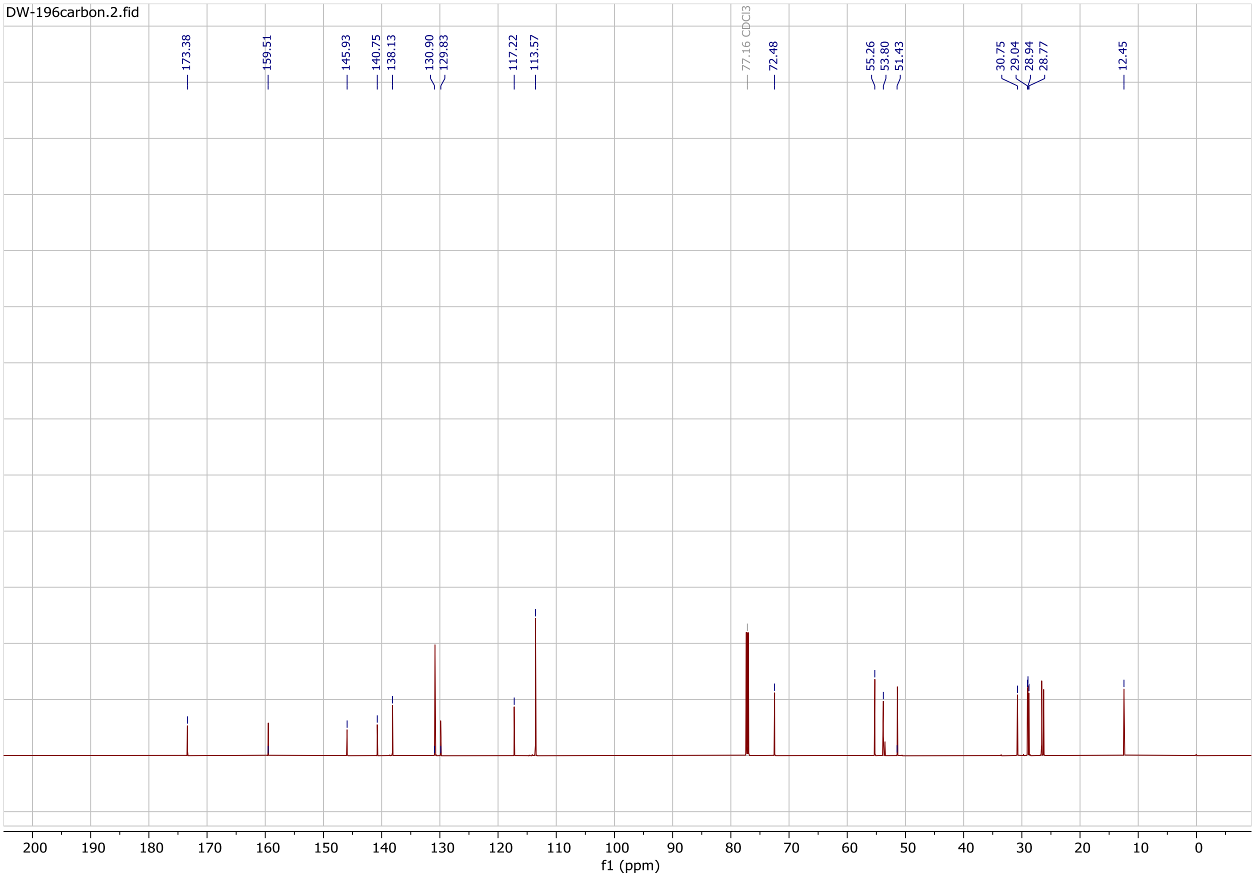

#### $^{1}H$ (700 MHz, $CDCl_3$) and $^{13}C$ (176 MHz, $CDCl_3$) NMR spectrum of compound 5

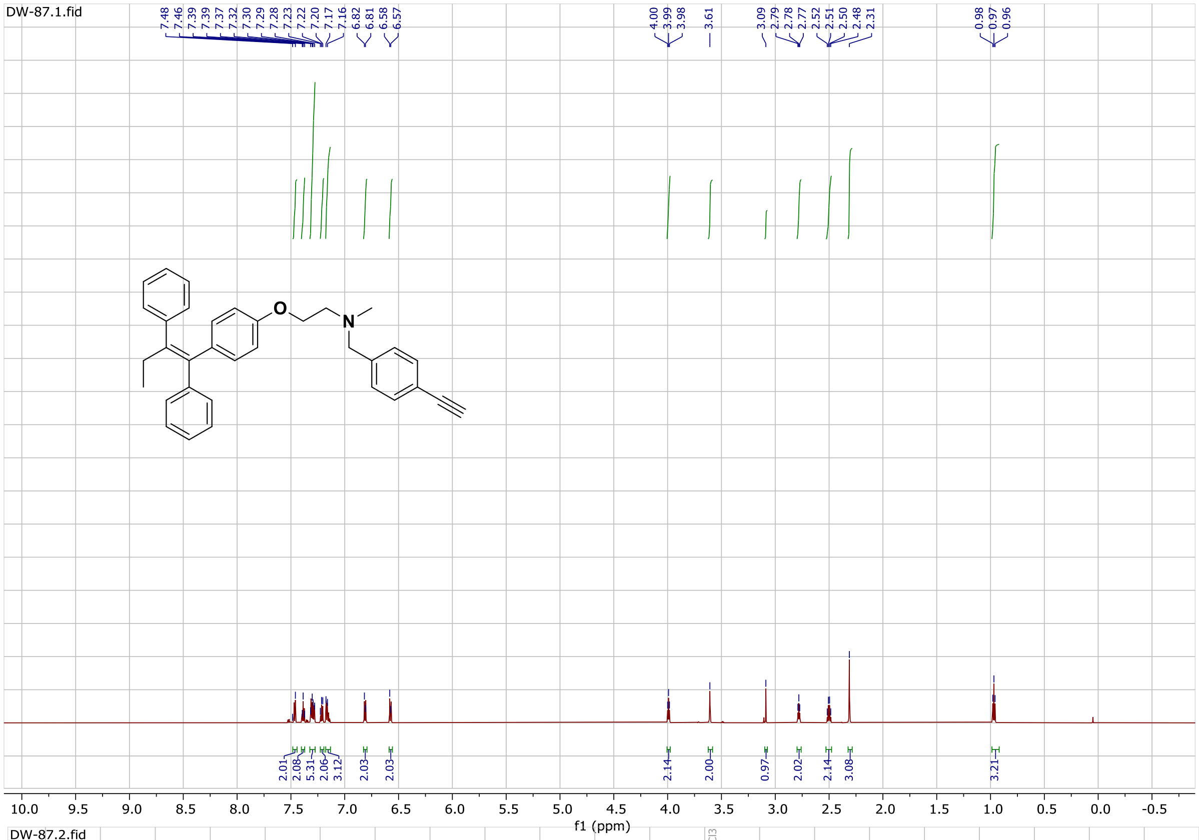


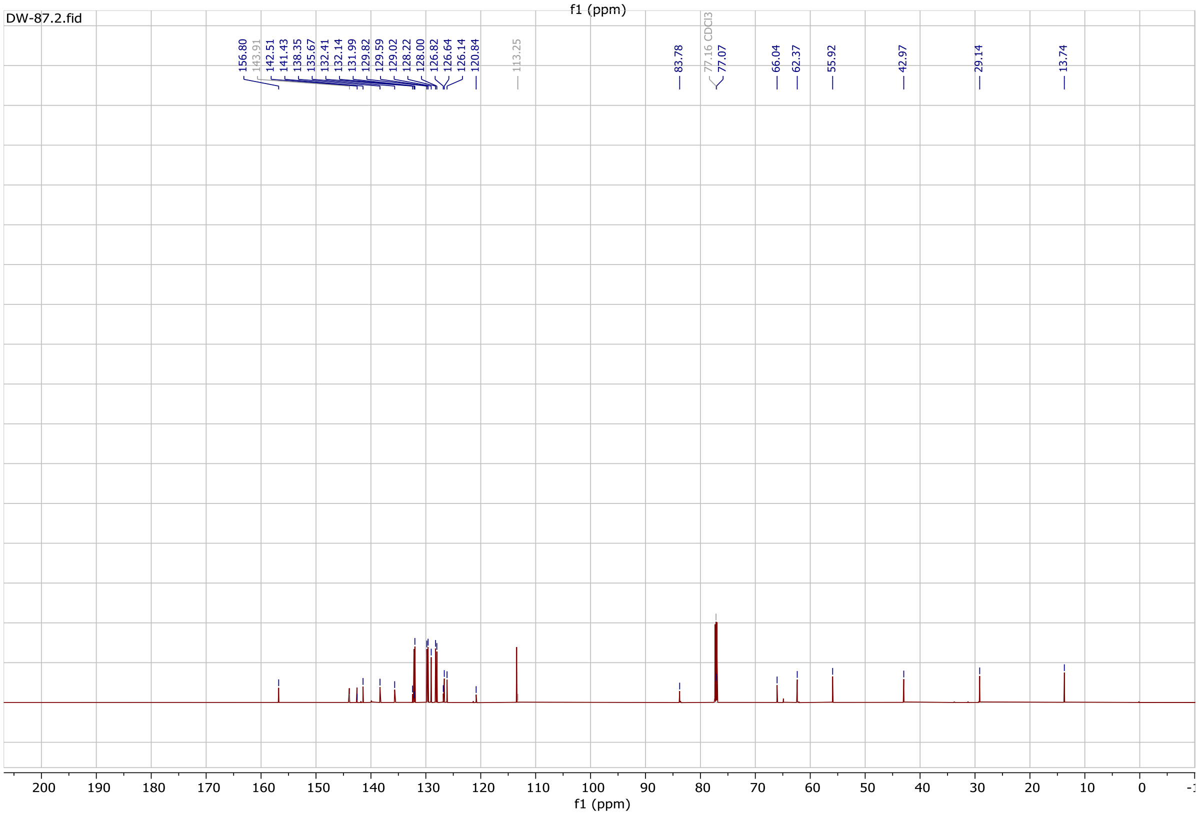

**$^1$H (700 MHz, $CDCl_3$) and $^{13}$C (176 MHz, $CDCl_3$) NMR spectrum of compound DW-54**

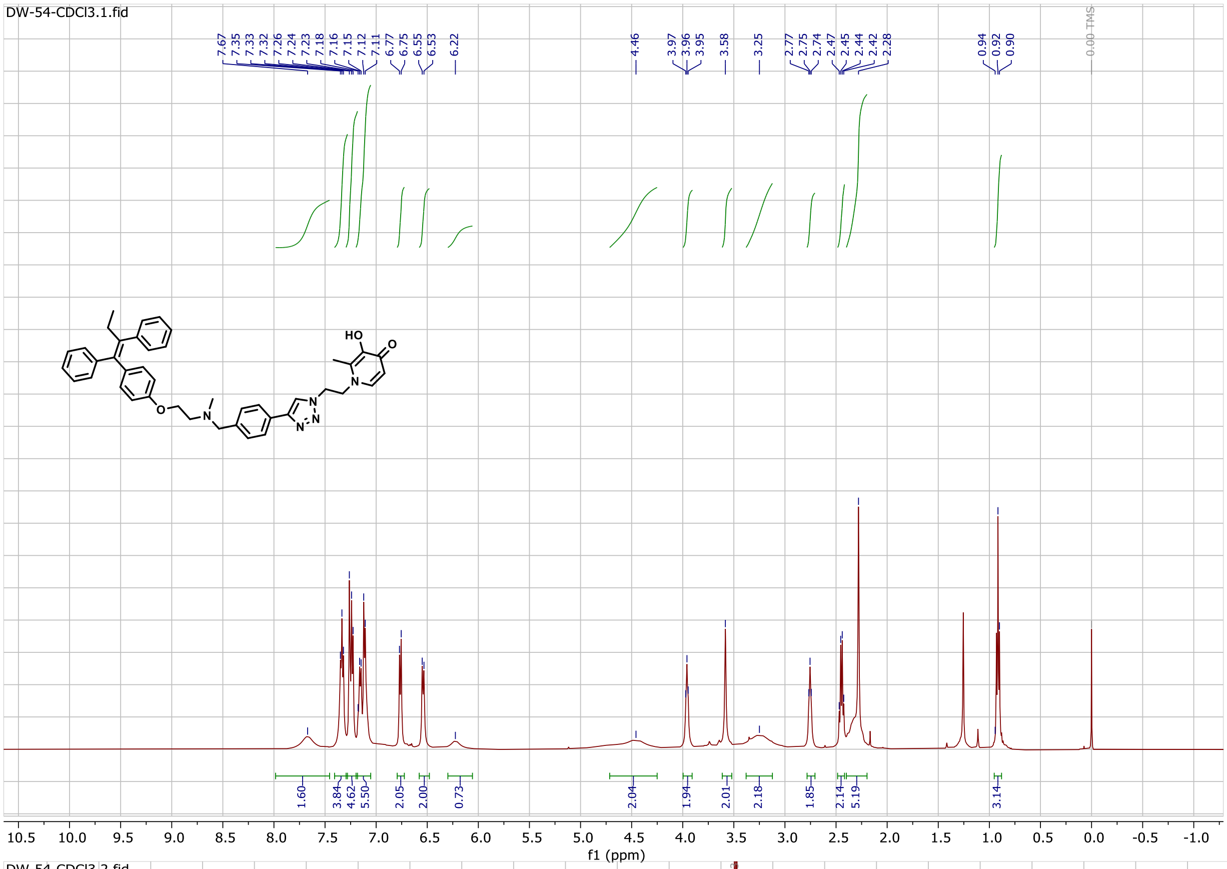

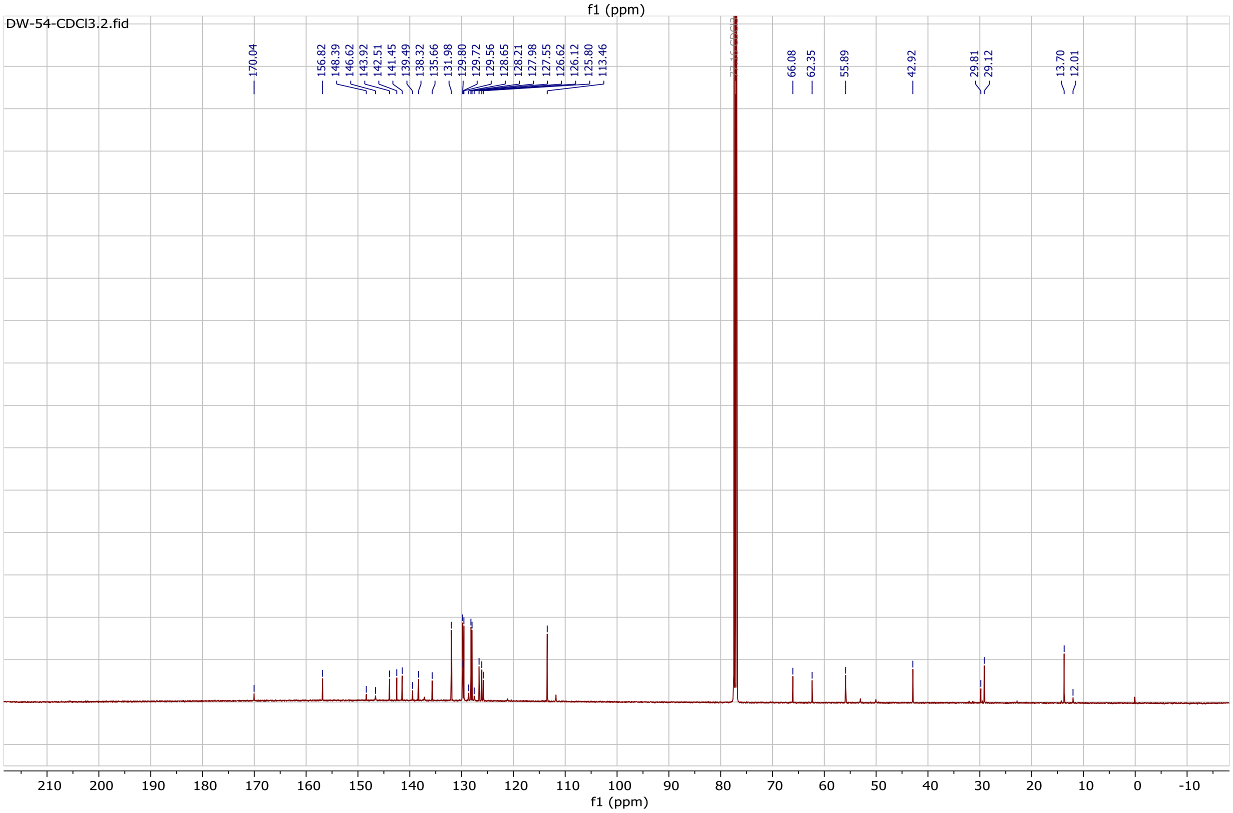

#### $^{1}H$ (700 MHz, $CDCl_3$) and $^{13}C$ (176 MHz, $CDCl_3$) NMR spectrum of compound DW-143

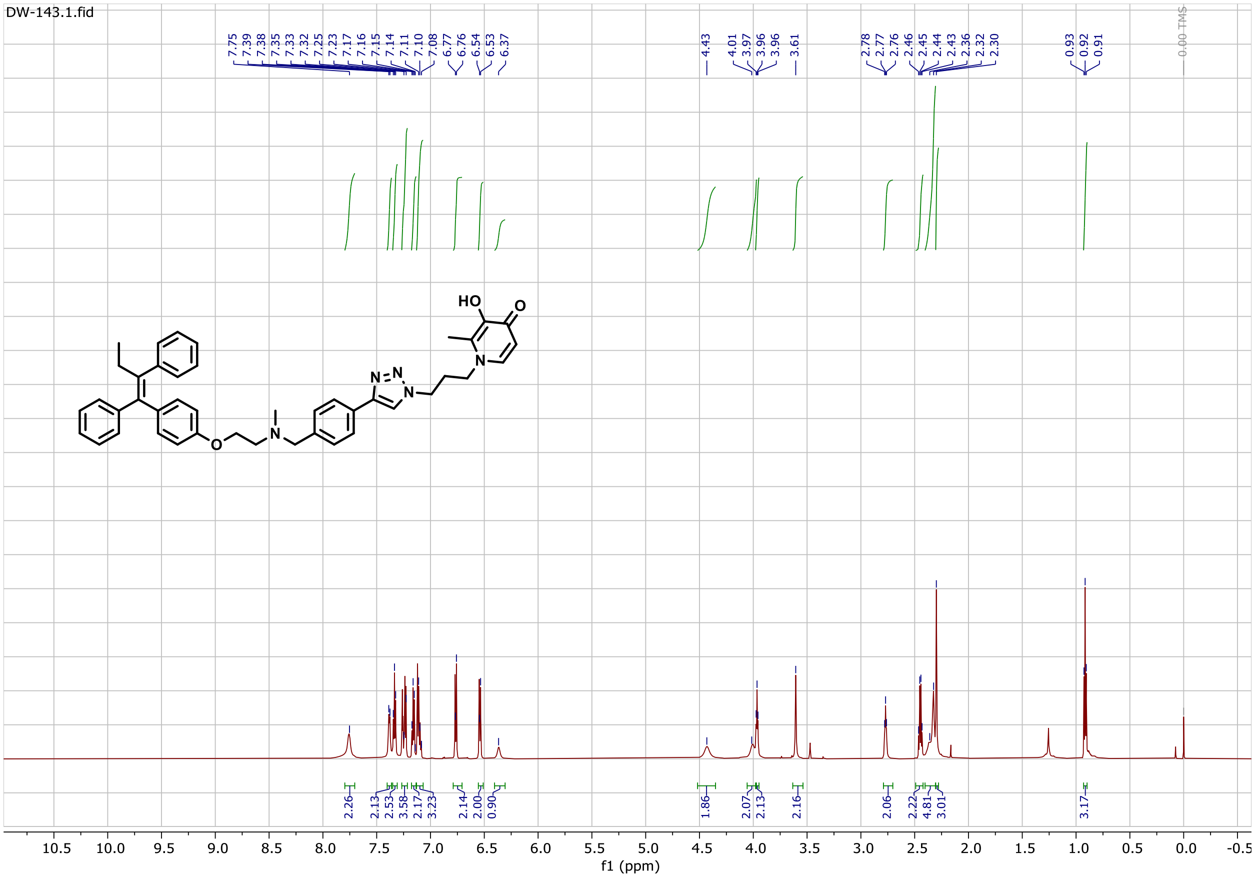


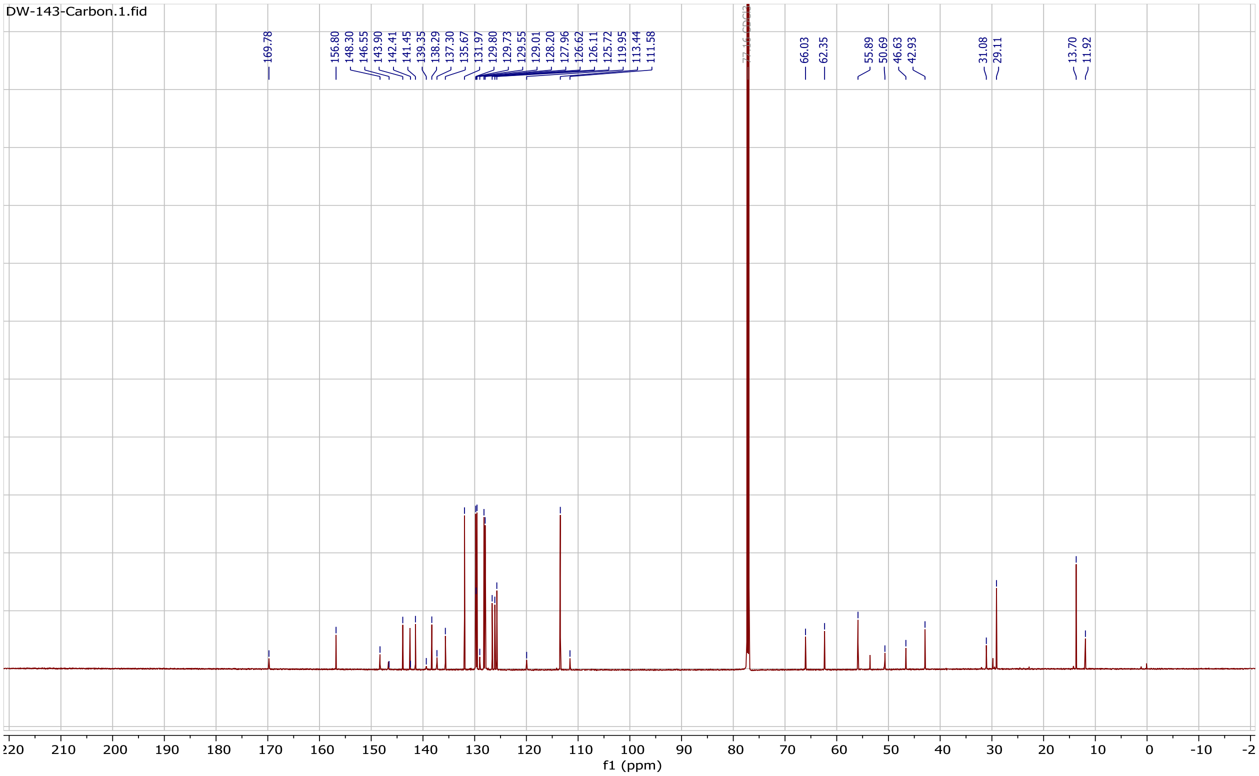

#### ¹H (700 MHz, $CDCl_3$) and ¹³C (176 MHz, $CDCl_3$) NMR spectrum of compound DW-136

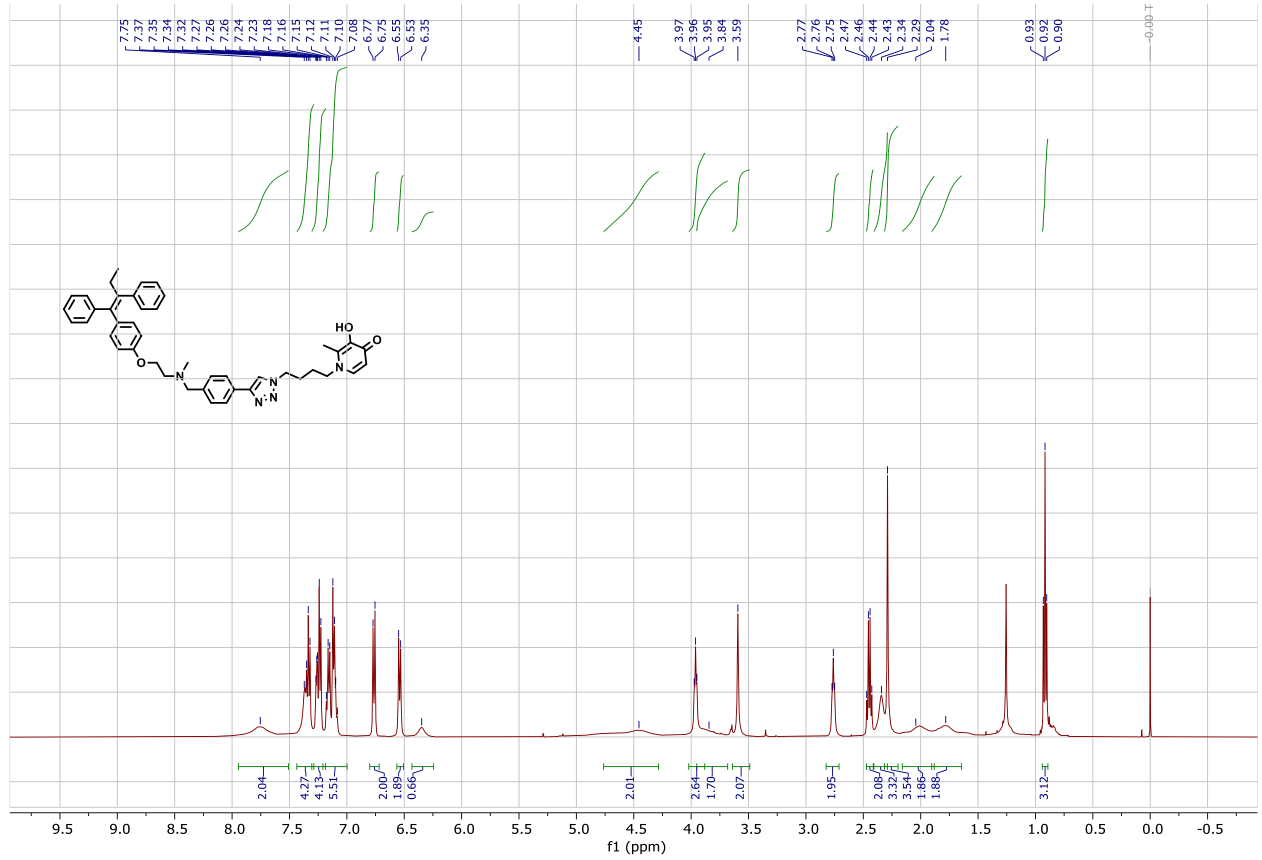

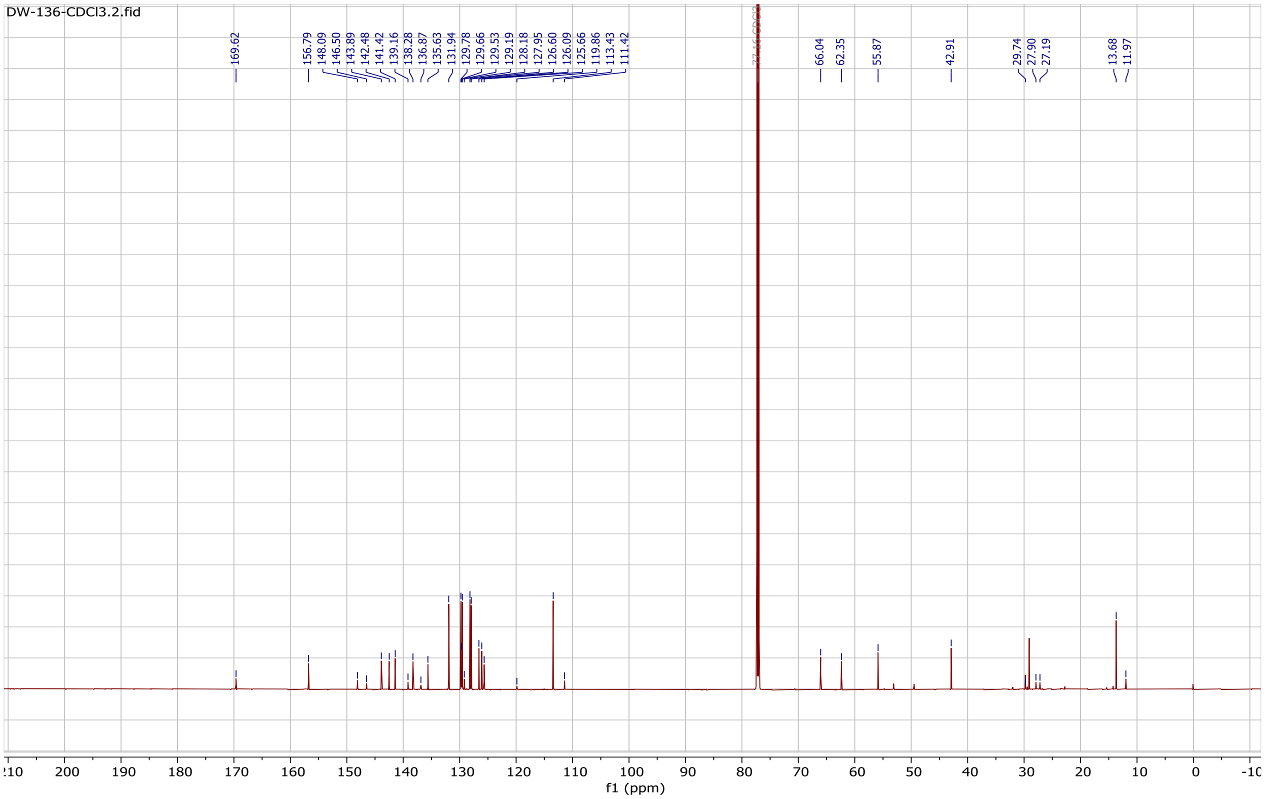

**[1]H (700 MHz, $CDCl_3$) and [13]C (176 MHz, $CDCl_3$) NMR spectrum of compound DW-116**

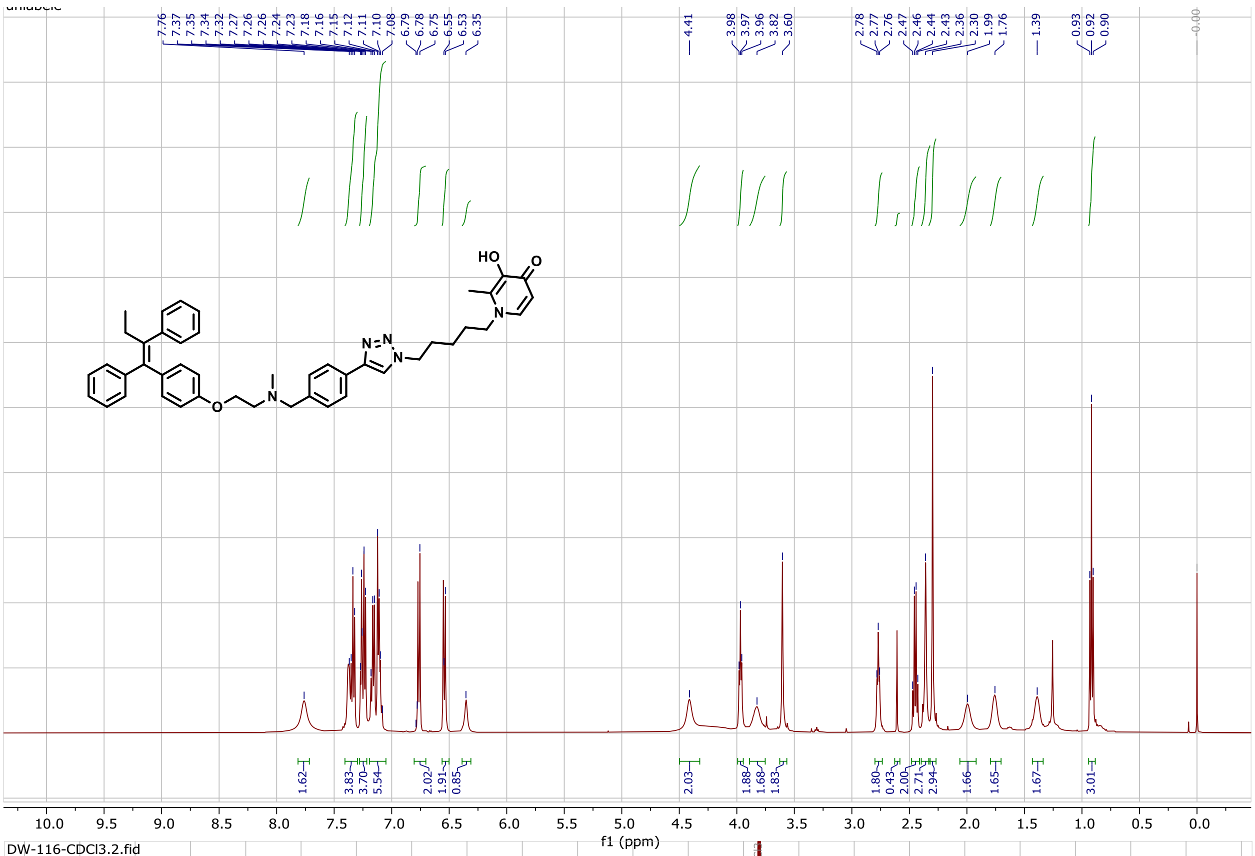


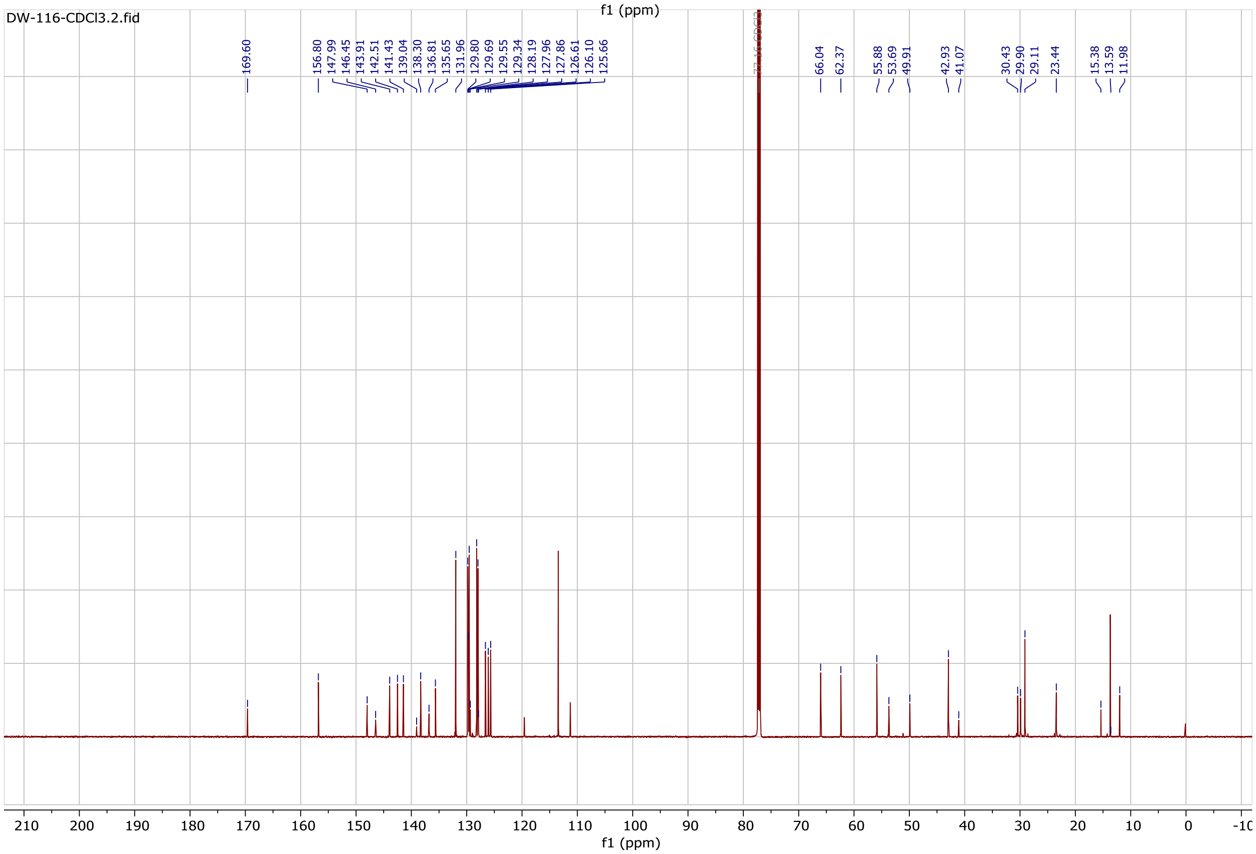

#### $^{1}$H (700 MHz, CD Cl$_3$) and $^{13}$C (176 MHz, CDCl$_3$) NMR spectrum of compound DW-122

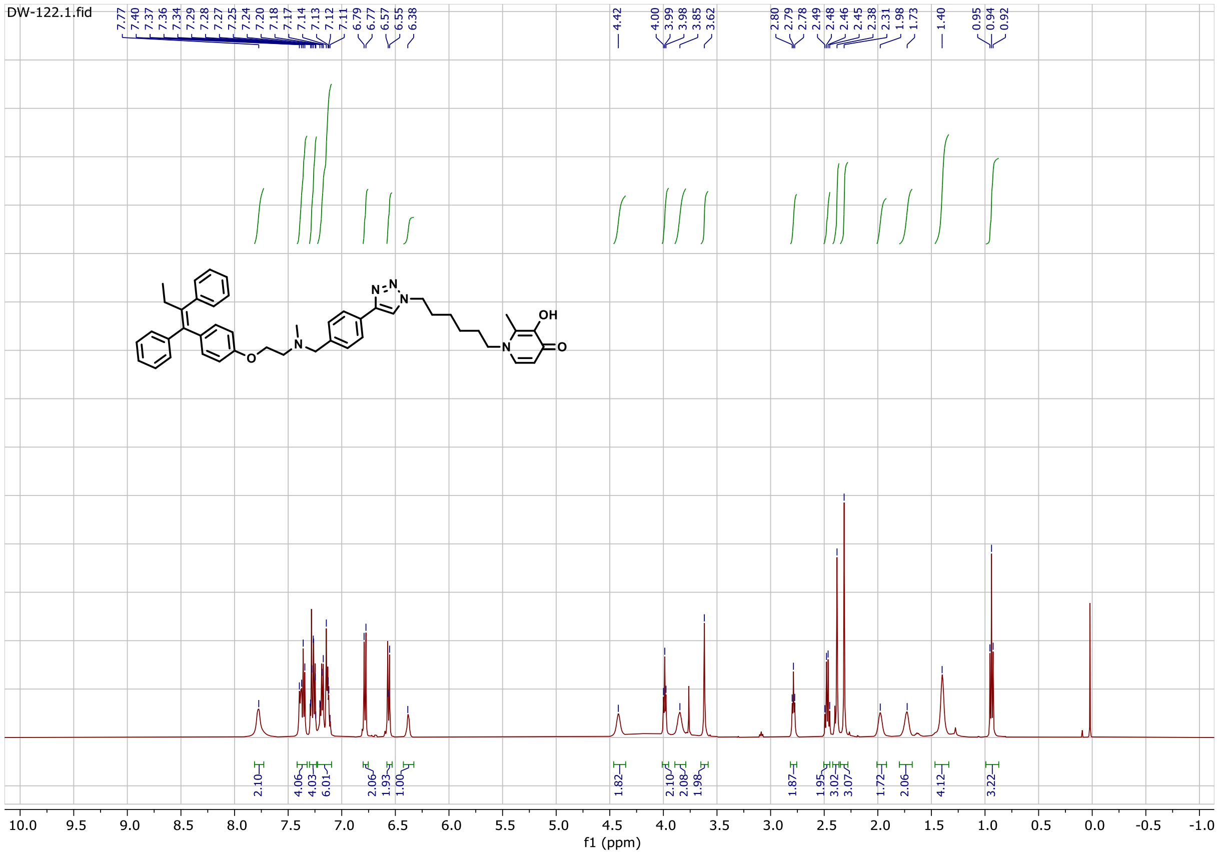

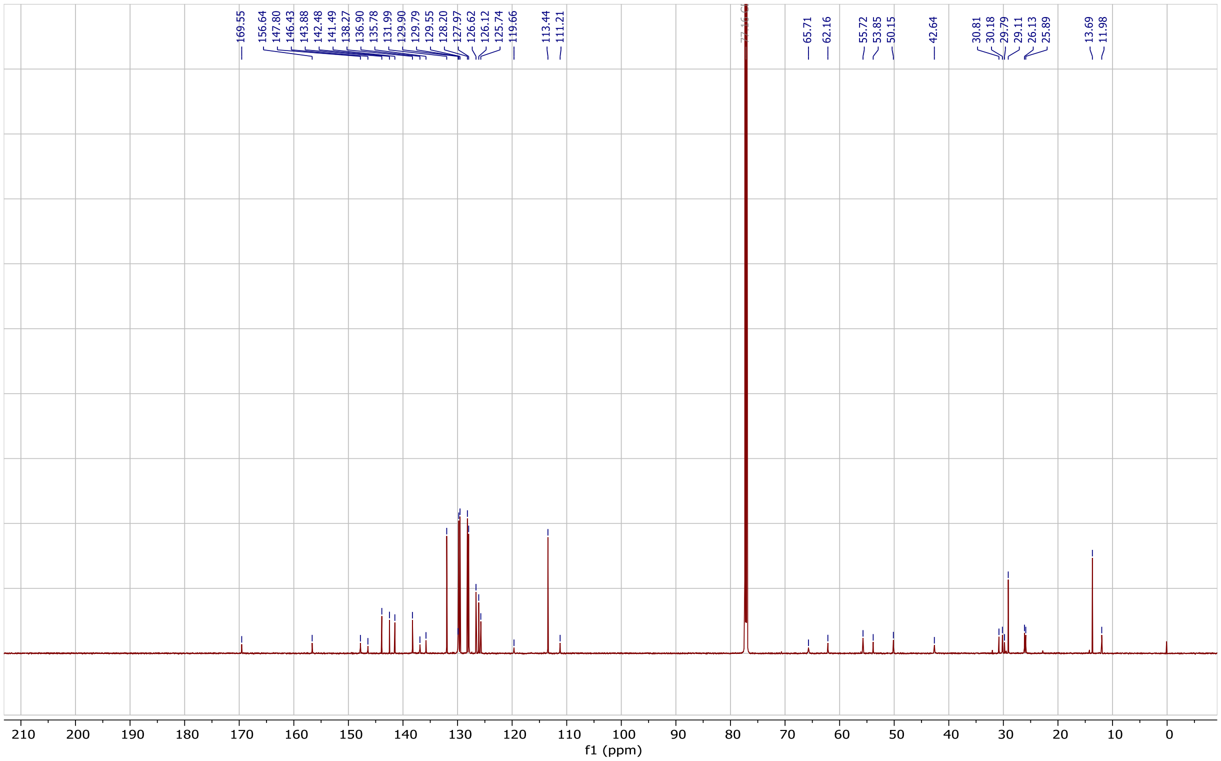

#### $^{1}H$ (700 MHz, $CDCl_3$) and $^{13}C$ (176 MHz, $CDCl_3$) NMR spectrum of compound DW-55

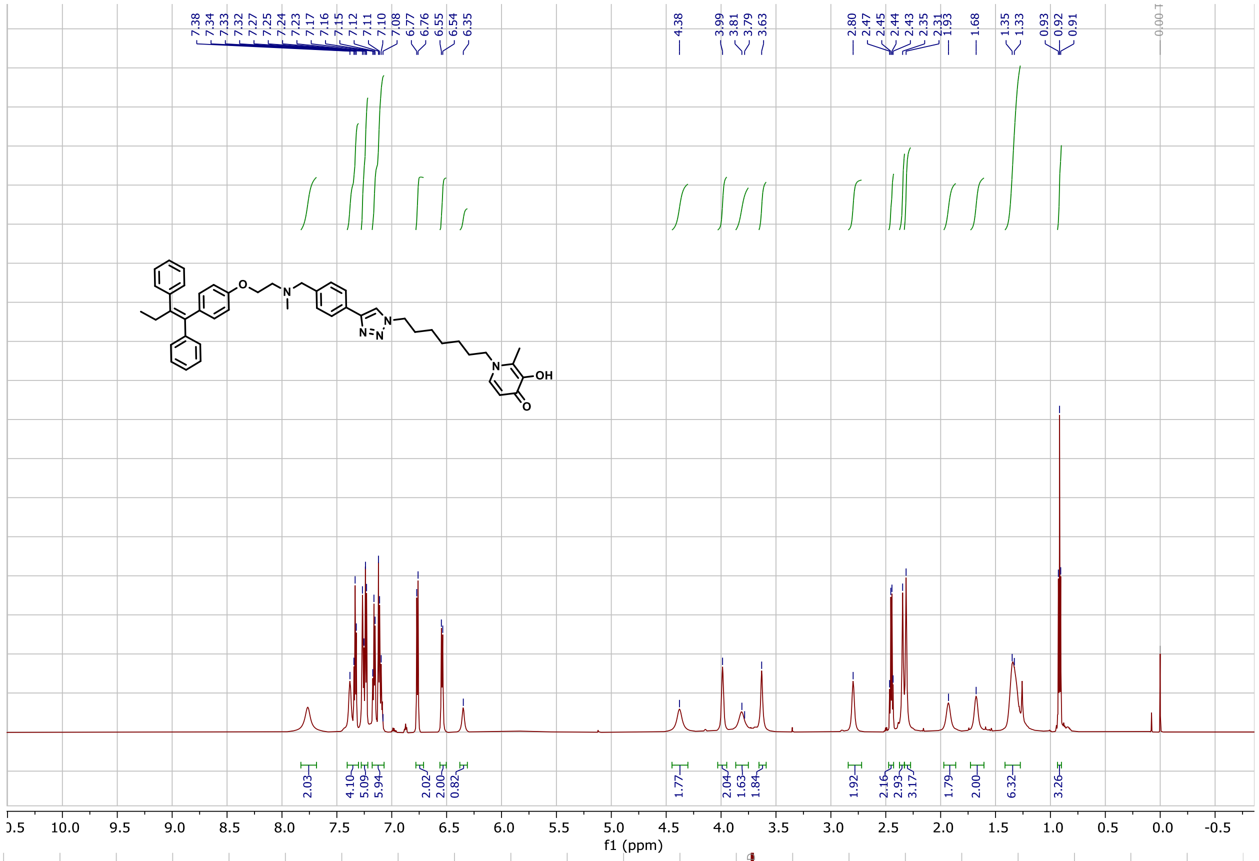


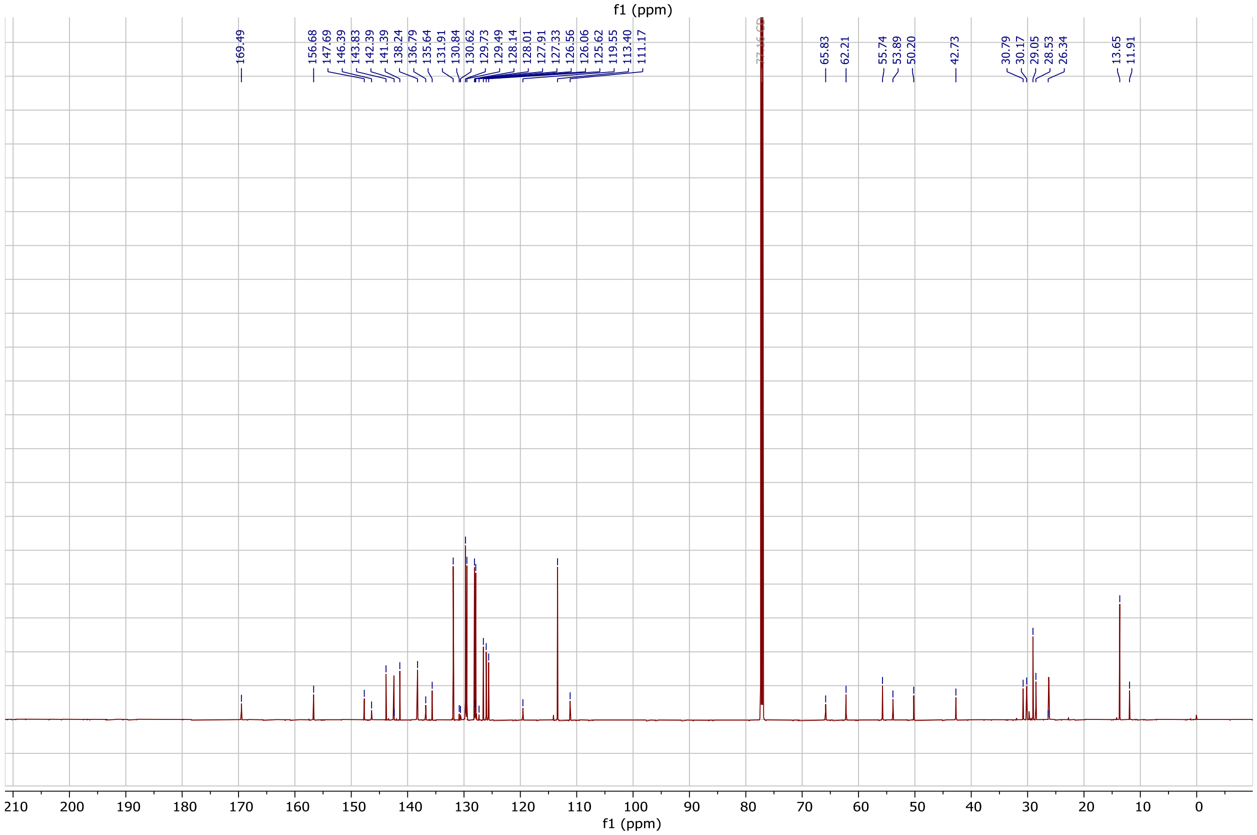

**$^{1}H$ (700 MHz, $CD_3OD+CDCl_3$) and $^{13}C$ (176 MHz, $CD_3OD+CDCl_3$) NMR spectrum of compound DW-554**

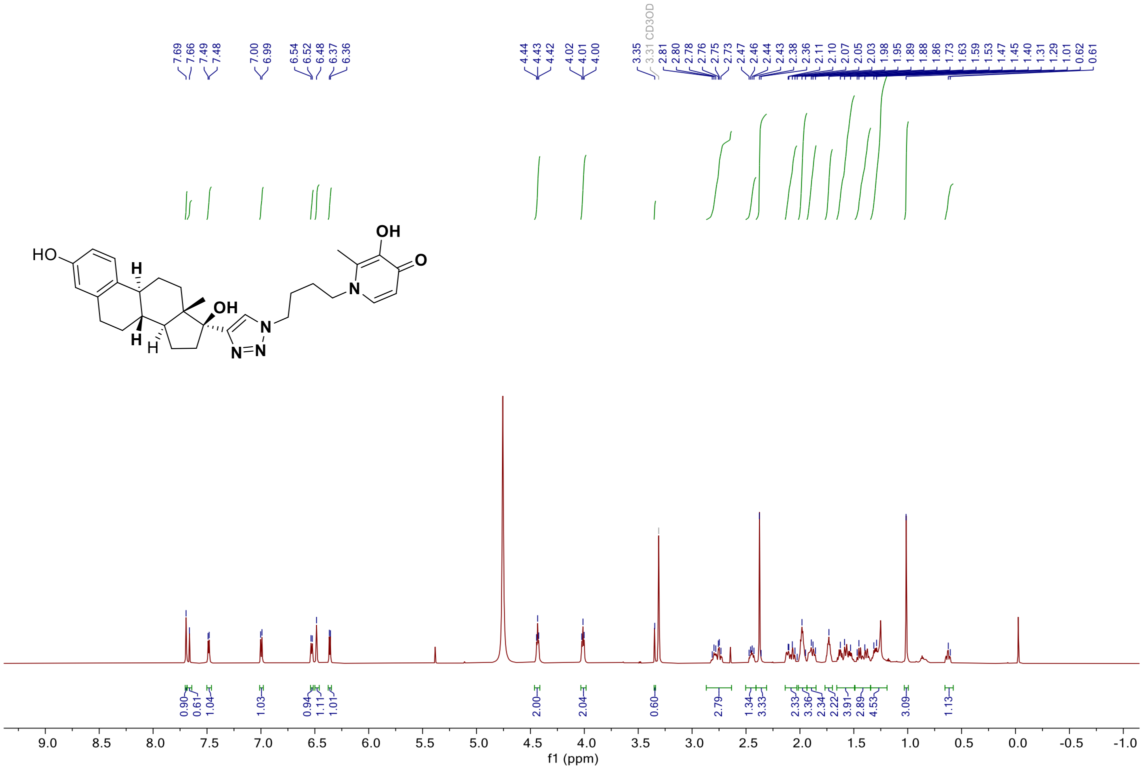


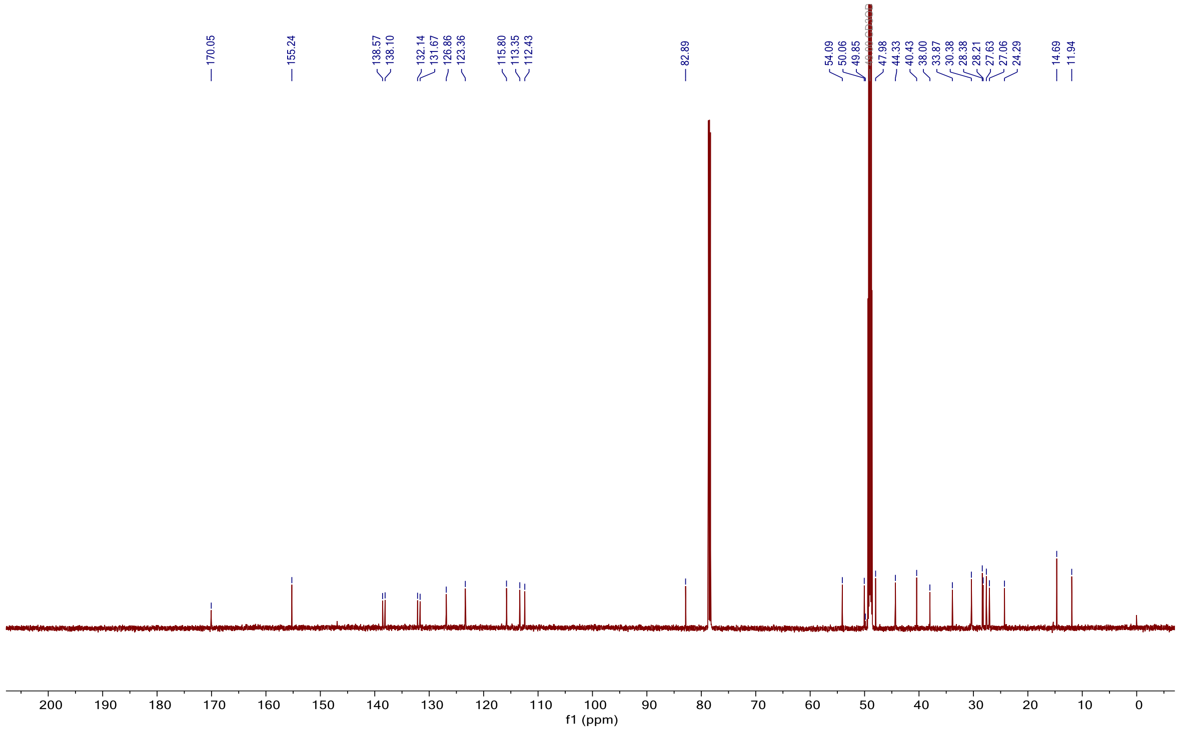

**[1]H (700 MHz, $CD_3OD+CDCl_3$) and [13]C (176 MHz, $CD_3OD+CDCl_3$) NMR spectrum of compound DW-613**

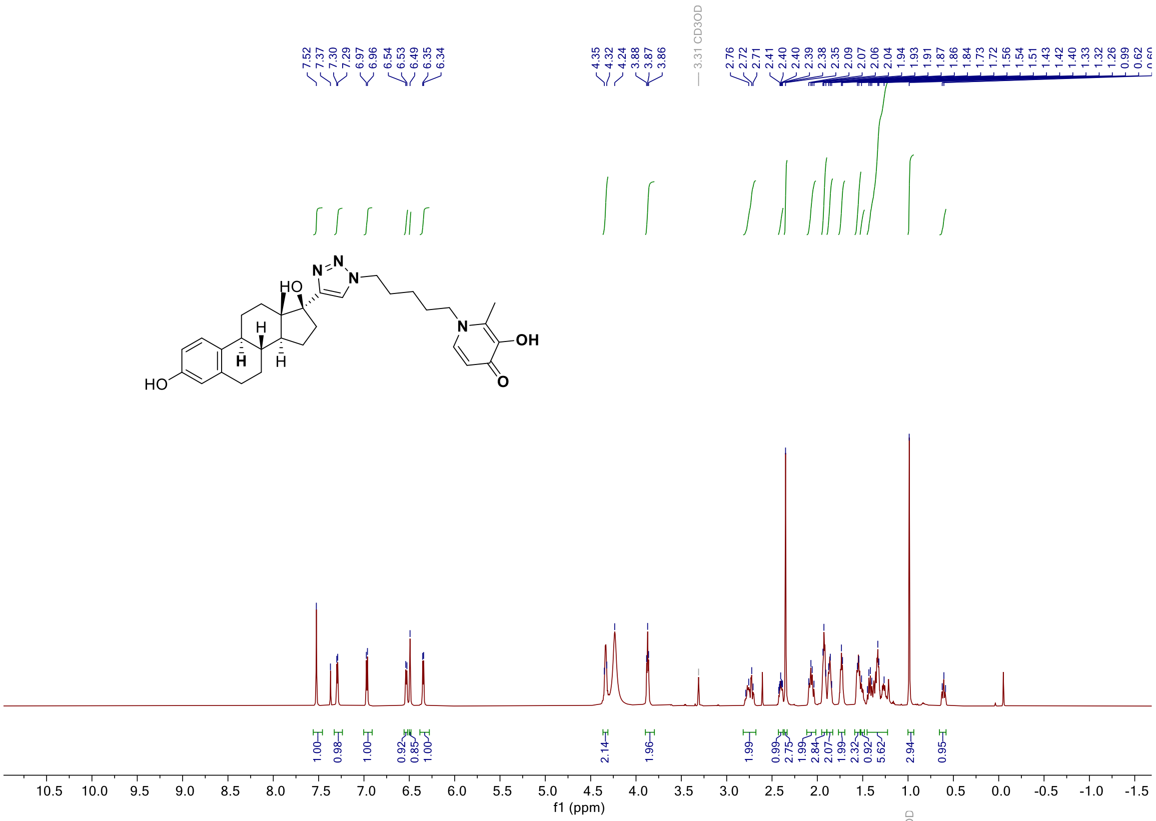


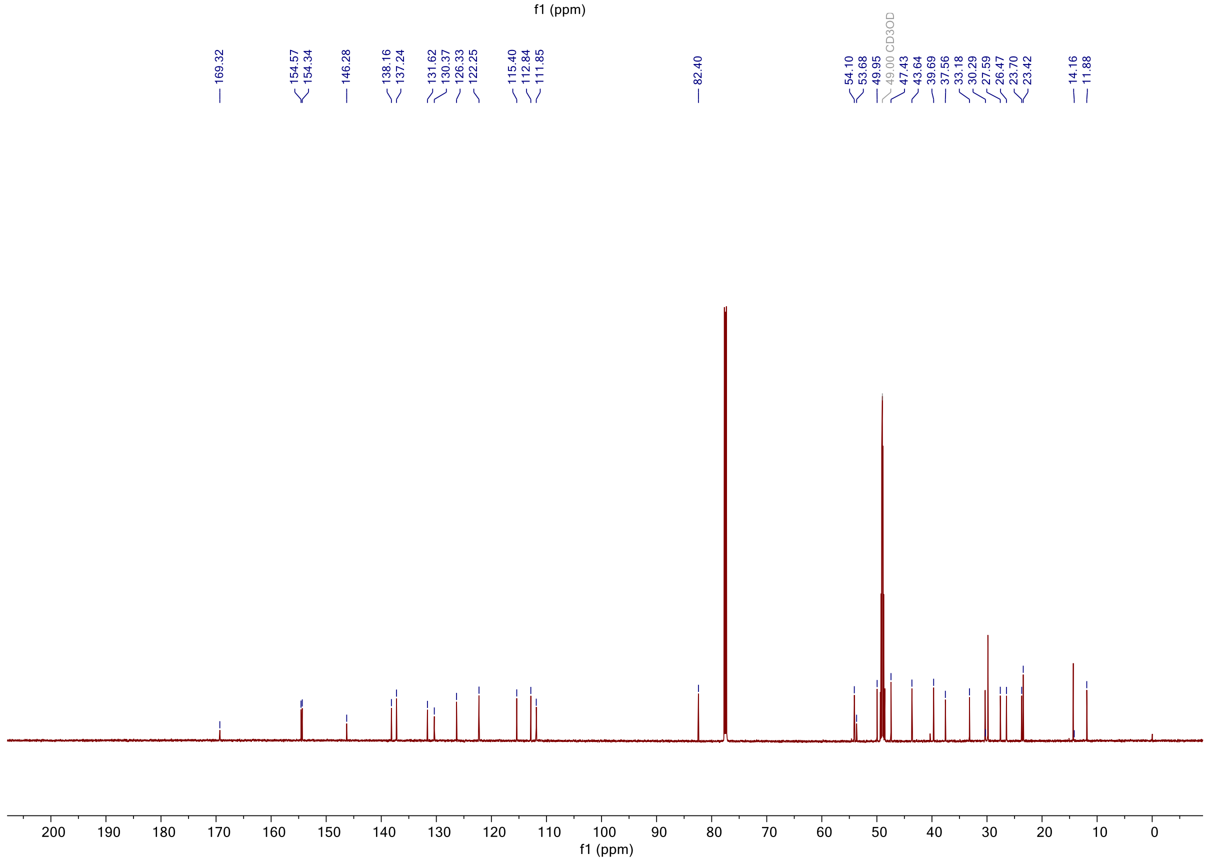

**$^{1}$H (700 MHz, $CD_3OD+CDCl_3$) and $^{13}$C (176 MHz $CD_3OD+CDCl_3$) NMR spectrum of compound DW-088**

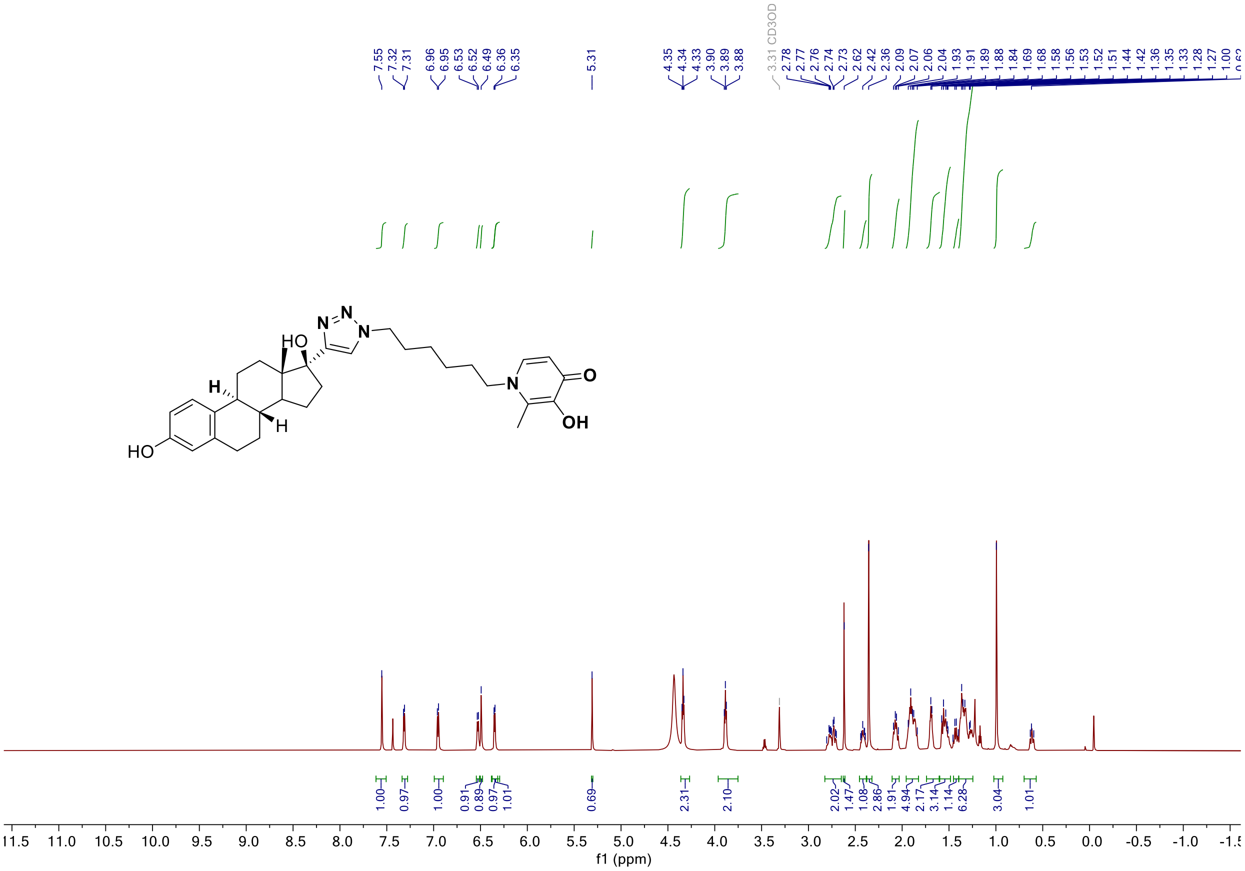

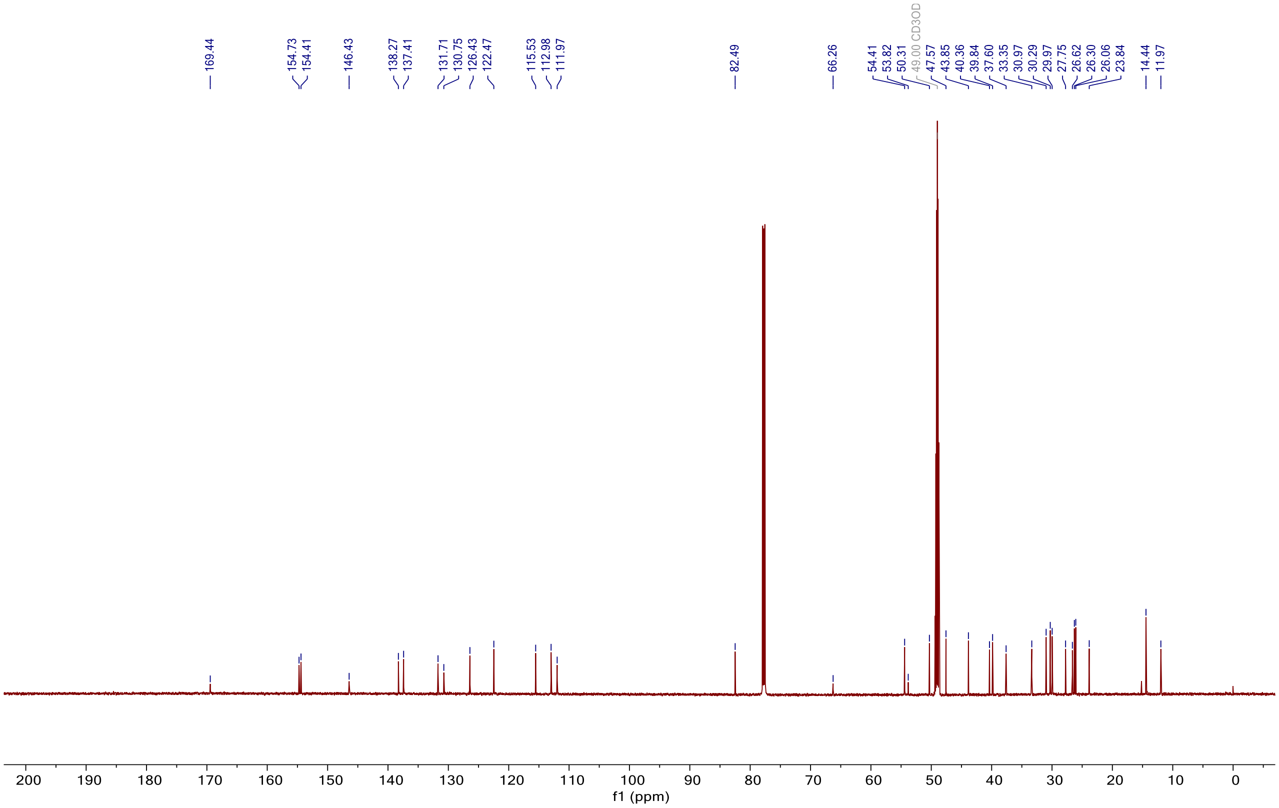

#### [1]H (700 MHz, $CD_3OD+CDCl_3$) and [13]C (176 MHz, $CD_3OD+CDCl_3$) NMR spectrum of compound DW-614

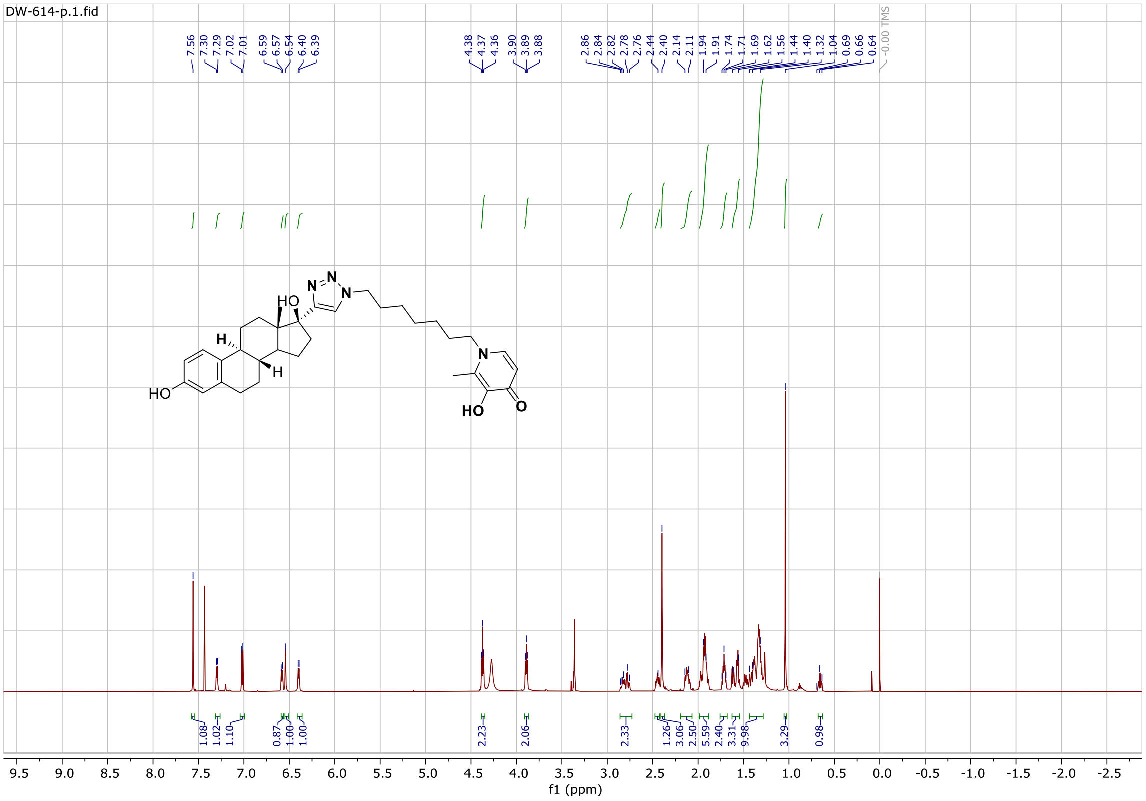

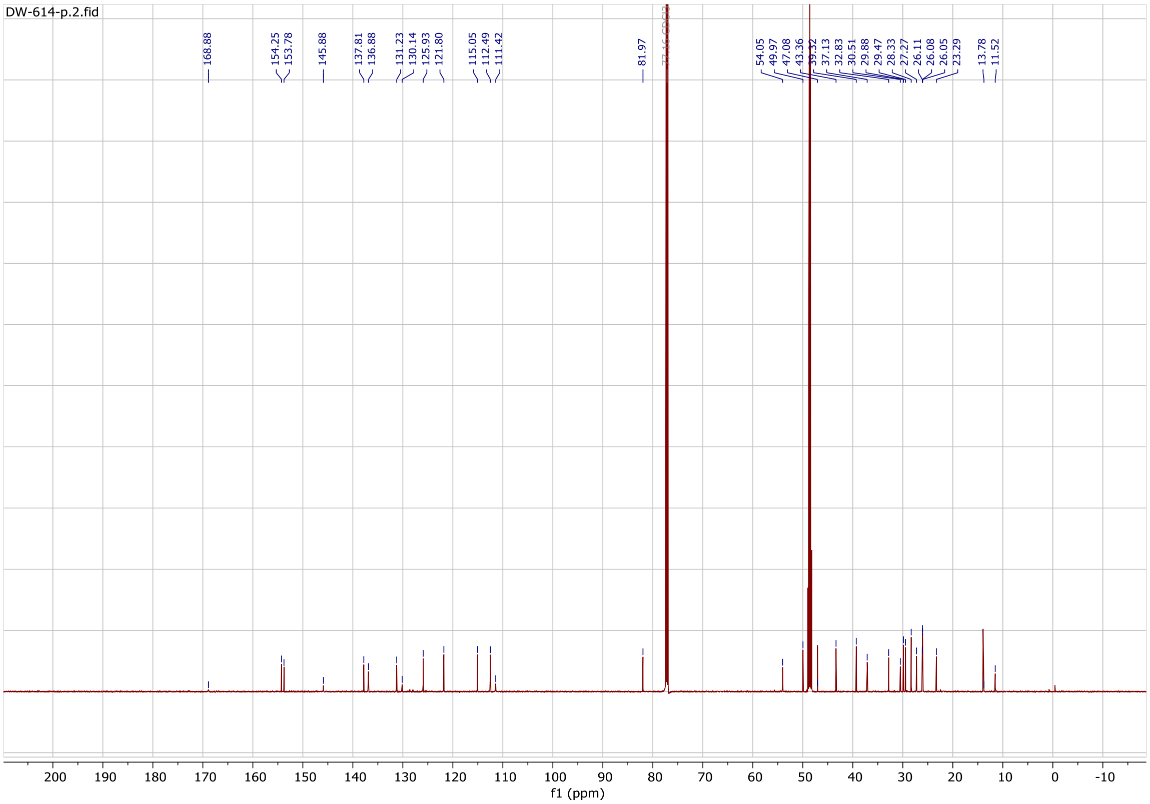

#### $^{1}H$ (700 MHz, $CD_3OD+CDCl_3$) and $^{13}C$ (176 MHz, $CD_3OD+CDCl_3$) NMR spectrum of compound DW-615

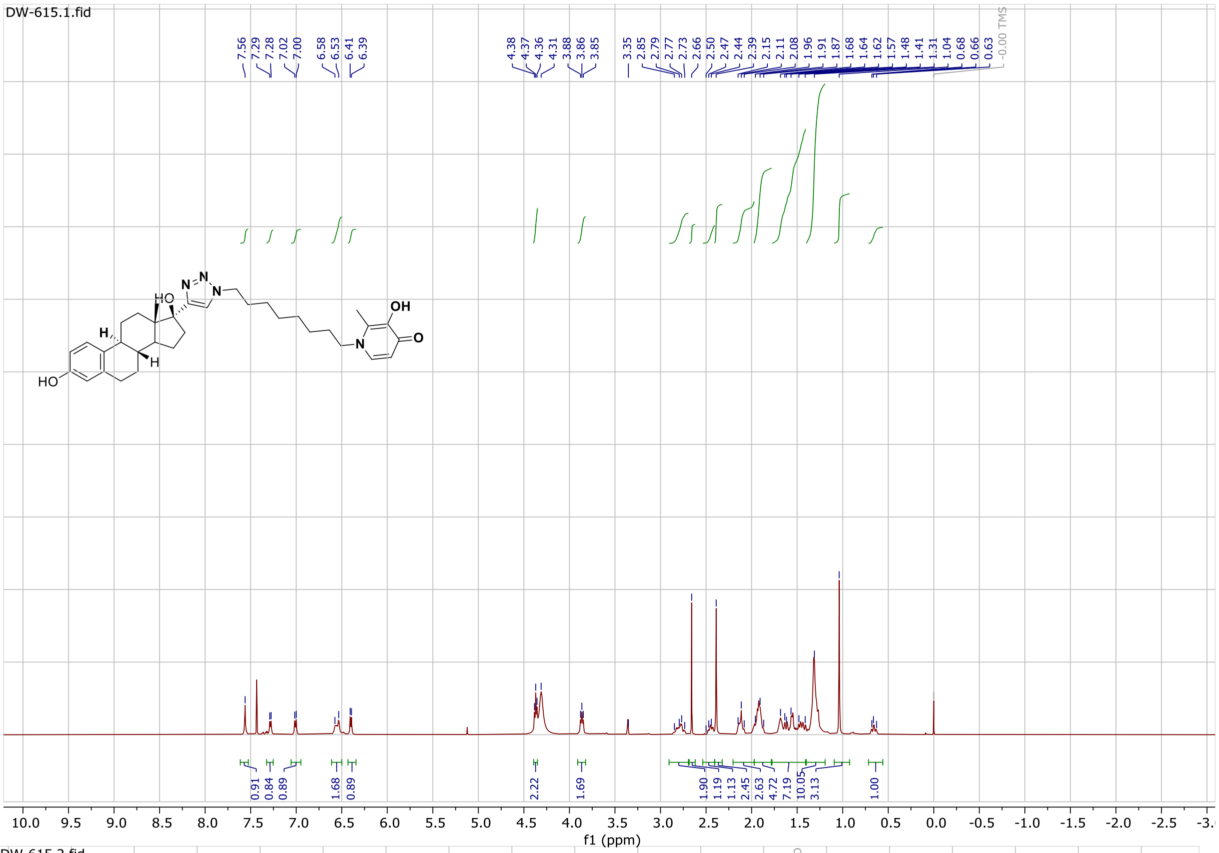


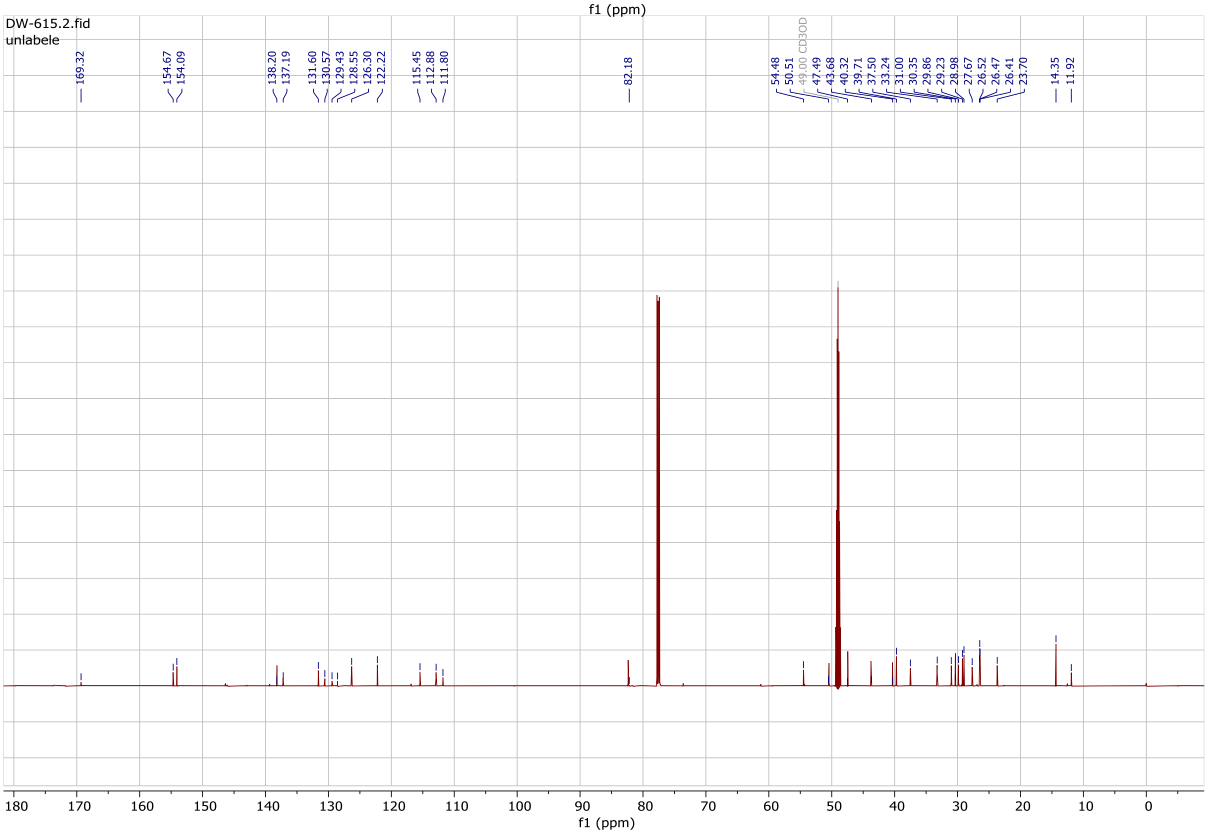

#### $^{1}H$ (700 MHz, $CDCl_3$) and $^{13}C$ (176 MHz, $CDCl_3$) NMR spectrum of compound DW-95

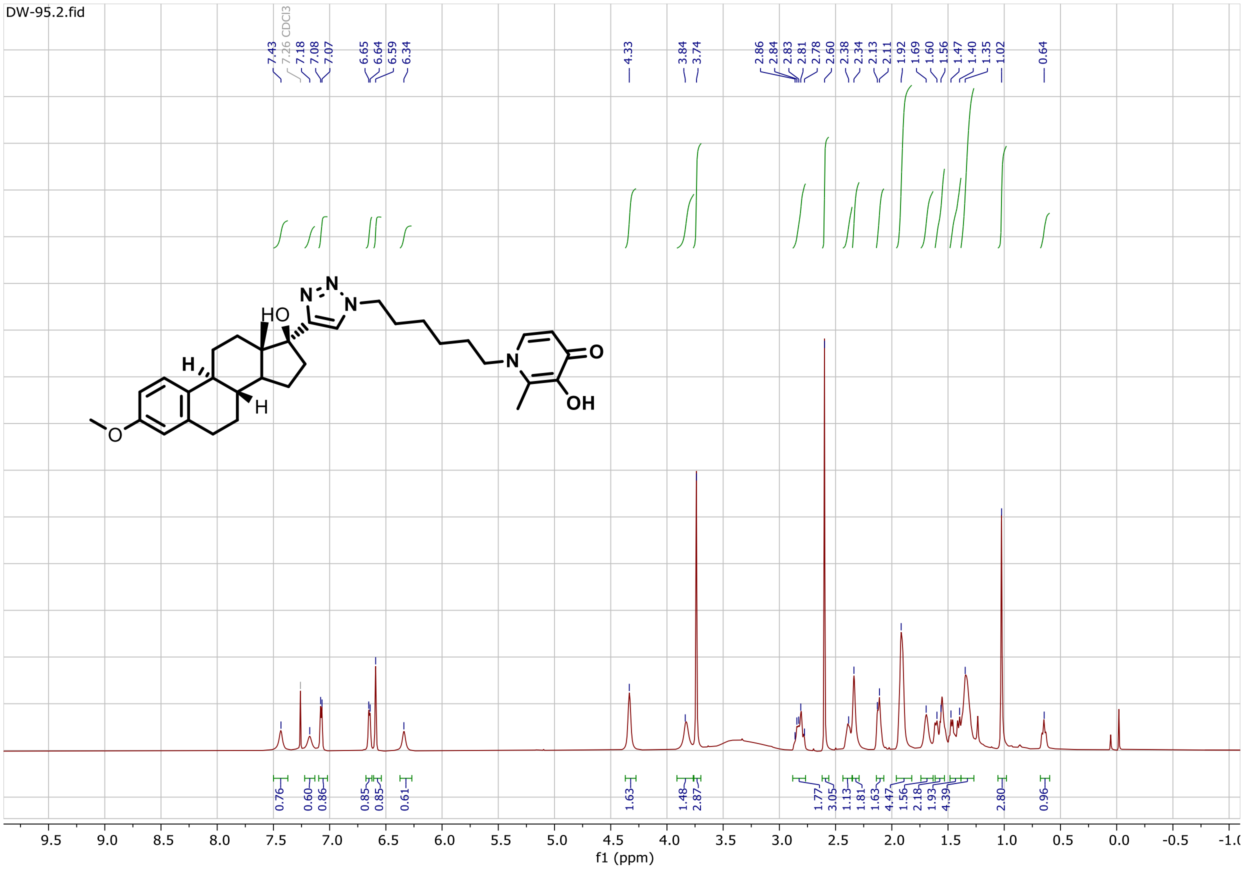


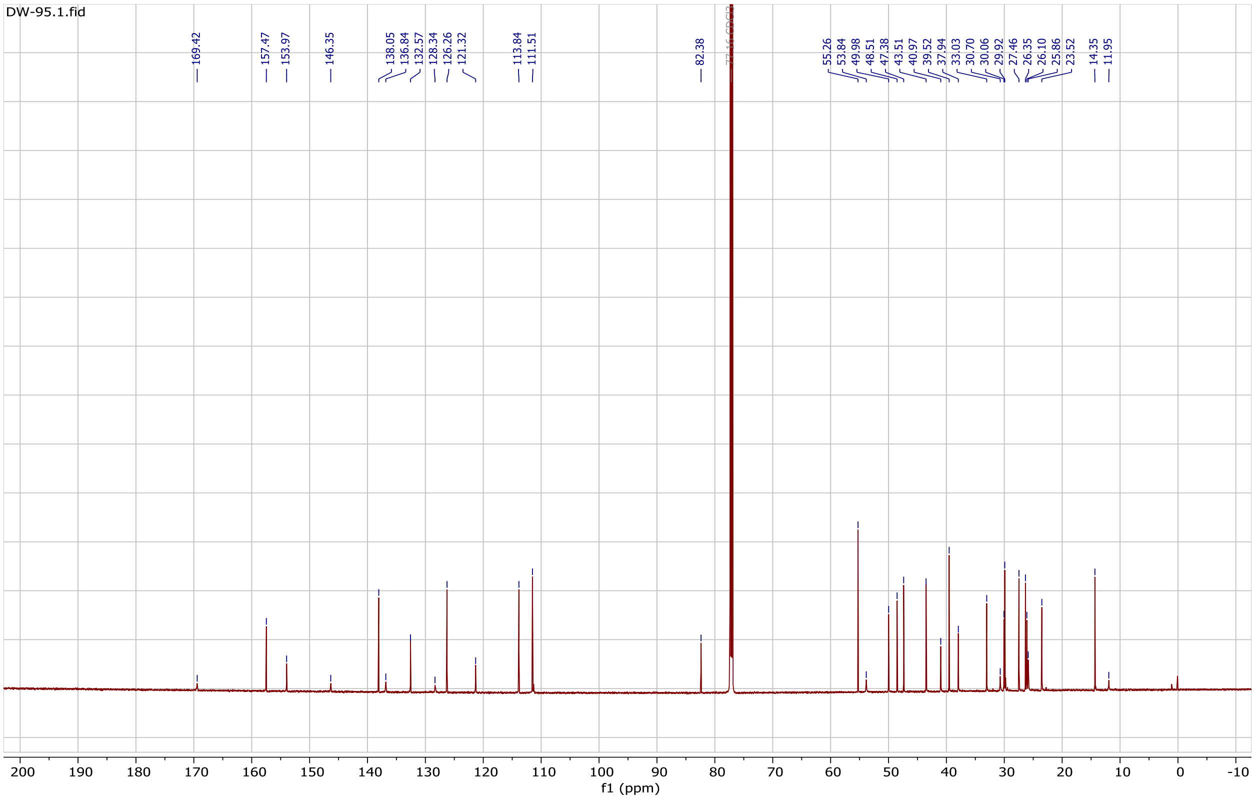

#### 1. HPLC Purity and UV peak at 254 nm of compound DW-54

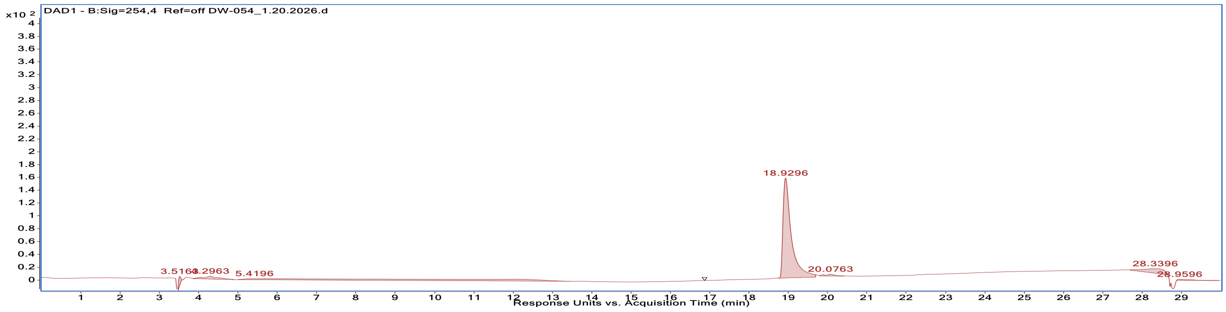


#### 2. HPLC Purity and UV peak at 254 nm of compound DW-55

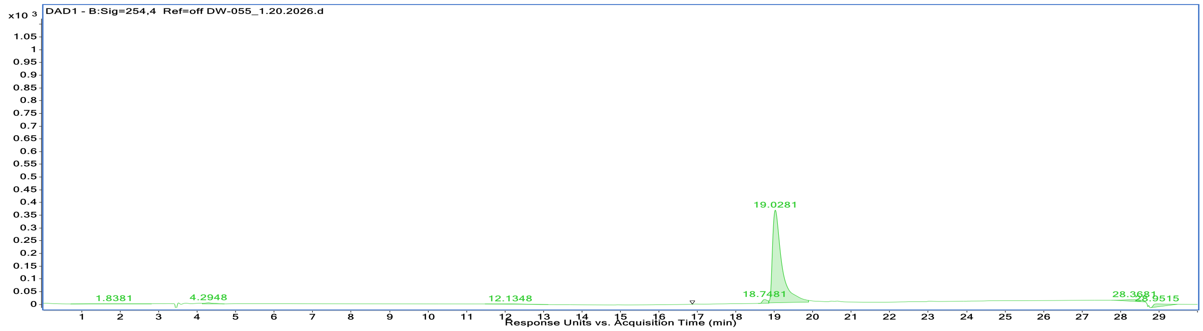


#### 3. HPLC Purity and UV peak at 254 nm of compound DW-136

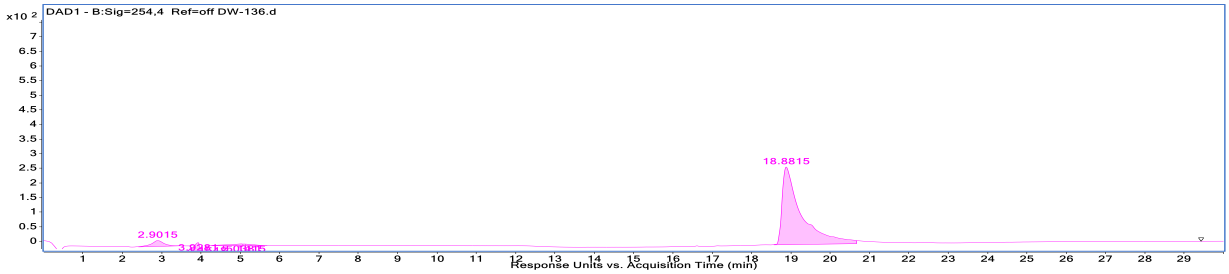

#### 4. HPLC Purity and UV peak at 254 nm of compound DW-143

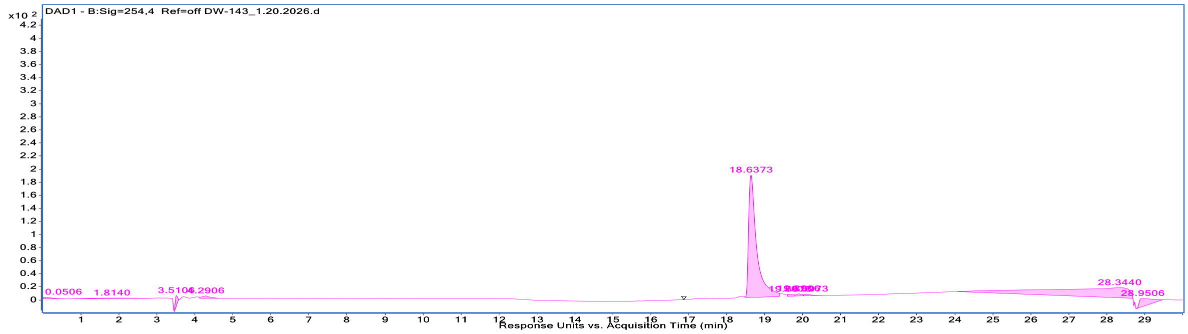


#### 5.HPLC Purity and UV peak at 254 nm of compound DW-116

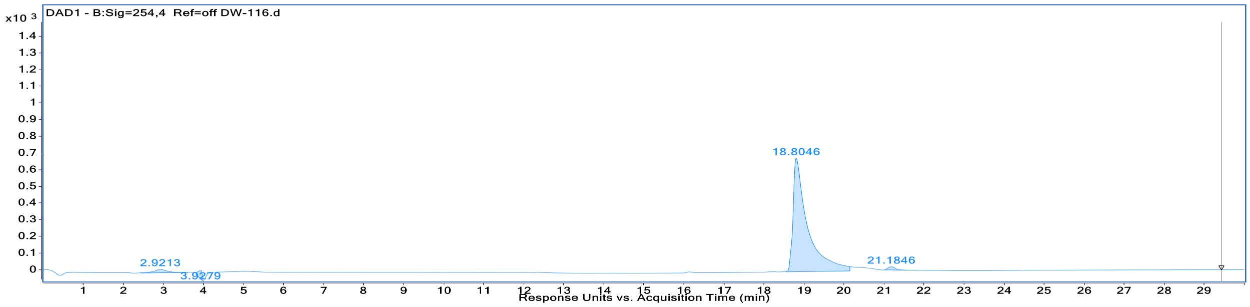


#### 6 .HPLC Purity and UV peak at 254 nm of compound DW-122

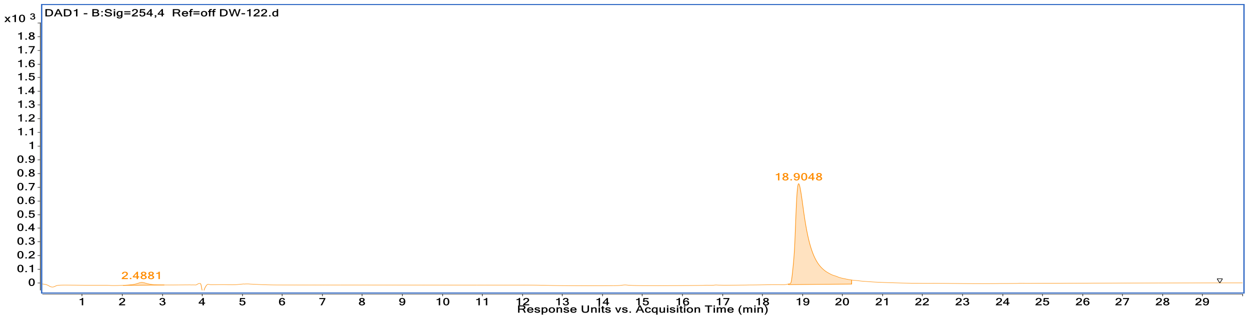

#### 7.HPLC Purity and UV peak at 254 nm of compound DW-088

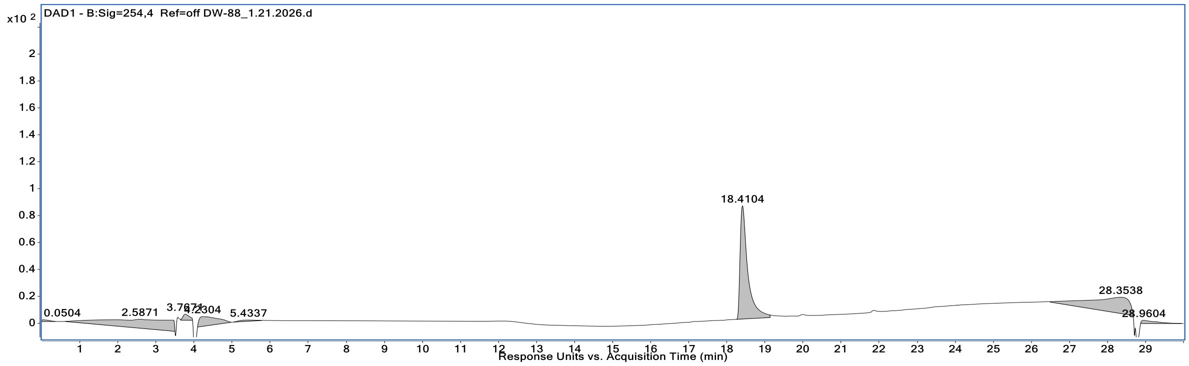


#### 8.HPLC Purity and UV peak at 254 nm of compound DW-95

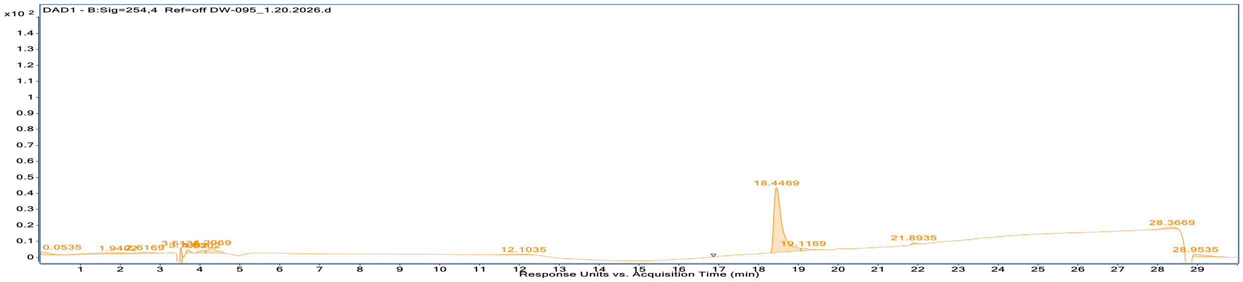


#### 9.HPLC Purity and UV peak at 254 nm of compound DW-554

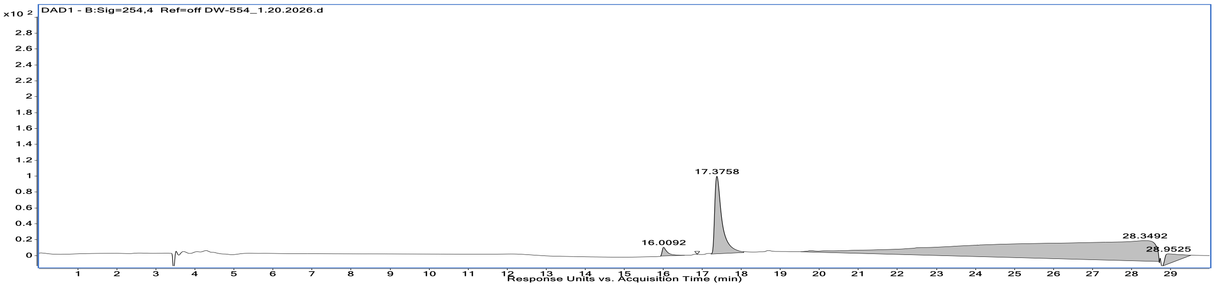

#### 10. HPLC Purity and UV peak at 254 nm of compound DW-613

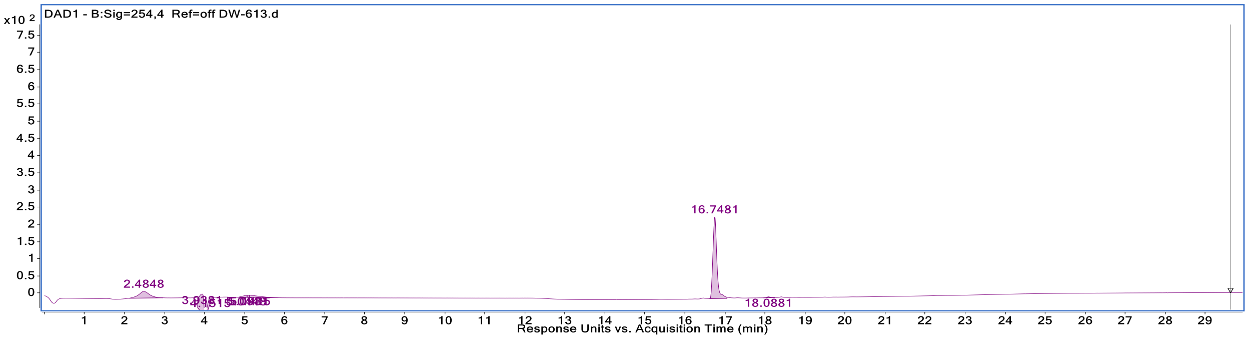


#### 11. HPLC Purity and UV peak at 254 nm of compound DW-614

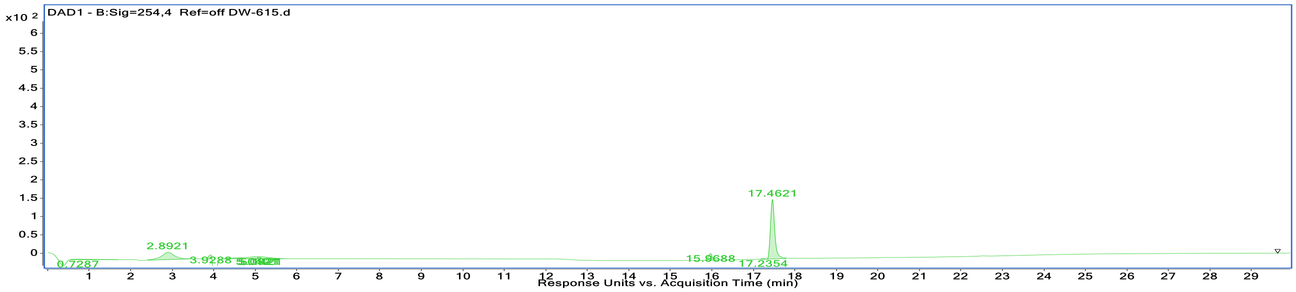


#### 12. HPLC Purity and UV peak at 254 nm of compound DW-615

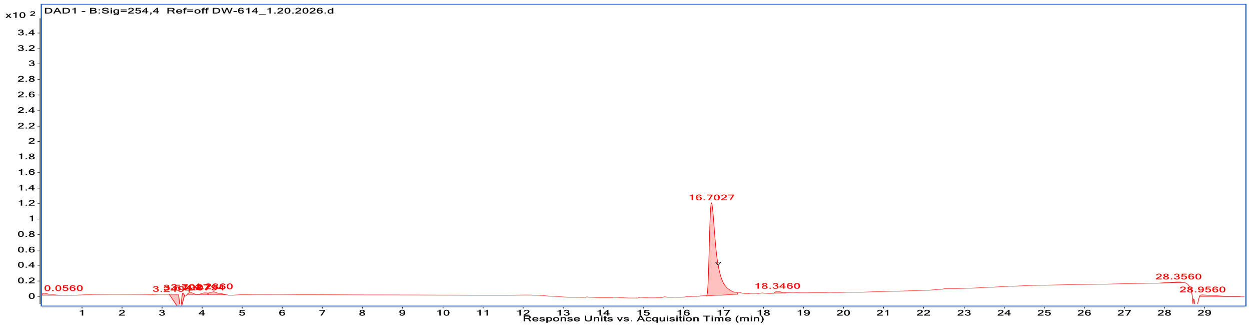
